# *APOE* E4 Alzheimer’s Risk Converges on an Oligodendrocyte Subtype in the Human Entorhinal Cortex

**DOI:** 10.1101/2025.11.20.689483

**Authors:** Louise A. Huuki-Myers, Heena R. Divecha, Svitlana V. Bach, Madeline R. Valentine, Nicholas J. Eagles, Bernard Mulvey, Rahul A. Bharadwaj, Ruth Zhang, James R. Evans, Melissa Grant-Peters, Ryan A. Miller, Joel E. Kleinman, Shizhong Han, Thomas M. Hyde, Stephanie C. Page, Daniel R. Weinberger, Keri Martinowich, Mina Ryten, Kristen R. Maynard, Leonardo Collado-Torres

## Abstract

The entorhinal cortex (ERC) is implicated in early progression of Alzheimer’s disease (AD). Here we investigated the impact of established biological risk factors for AD, including *APOE* genotype (E2 versus E4 alleles), sex, and ancestry, on gene expression in the human ERC. We generated paired spatially-resolved transcriptomics (SRT) and single-nucleus RNA sequencing data (snRNA-seq) in postmortem human ERC tissue from middle aged brain donors with no history of AD. *APOE*-dependent changes in gene expression predominantly mapped to a transcriptionally-defined oligodendrocyte subtype, which varied substantially with ancestry, and suggested differences in oligodendrocyte differentiation and myelination. Integration of SRT and snRNA-seq data identified a common gene expression signature associated with *APOE* genotype, which we localized to the same oligodendrocyte subtype and a white matter spatial domain. This suggests that AD risk in ERC may be associated with disrupted oligodendrocyte function, potentially contributing to future neurodegeneration.

**Lay Summary:** Alzheimer’s disease (AD) is a neurodegenerative disorder that accounts for 60-80% dementia cases. Apolipoprotein E (*APOE)* genotype is the strongest genetic risk factor for AD, and the entorhinal cortex (ERC) is a brain region implicated in its earliest progression. Our study investigated how *APOE* genotype impacts gene expression in the ERC. We identified genotype-dependent effects on oligodendrocytes with different transcriptional profiles related to maturation that may help explain how *APOE* genotype mediates its effects on AD risk.

## 1| Introduction

Alzheimer’s disease (AD) is a neurodegenerative disorder that accounts for 60-80% of dementia cases, constituting a major health challenge^1^. AD is characterized by accumulation of extracellular amyloid-beta (Aß) plaques and hyperphosphorylated tau (pTau) neurofibrillary tangles, which are associated with inflammation, neuronal death, and brain atrophy^1,2^. Since neuronal loss becomes rapid and largely irreversible once substantial AD pathology has accumulated, current therapeutic strategies emphasize identifying biomarkers and drug targets at the earliest stages of disease, before widespread neurodegeneration occurs. The recent approval of the first disease-modifying treatments that act on underlying pathology^3^ further underscores the critical importance of early detection and intervention in AD, as these treatments are only approved for individuals in early AD stages.

There are several documented epidemiological factors that impact risk for developing AD. Americans of African ancestry (AA) are more likely to develop AD than Americans of European ancestry (EA), even when controlling for environmental factors^4^. Females are at overall higher risk of developing AD, but the effect of sex on risk is modified by both age and ancestry^5,6^. However, the strongest genetic risk factor for AD is allelic variation at the Apolipoprotein E (*APOE)* locus, with *APOE* E4 increasing risk and *APOE* E2 reducing risk^7^. *APOE* E4 is associated with higher levels of amyloid beta deposition and pTau pathology, while *APOE* E2 delays pathology^7^. Importantly, ancestry also interacts with the effects of *APOE* allelic risk. For instance, E2 and E4 alleles are more common in AA compared to EA populations, but the protective and risk effects are stronger in EA^4^. Additionally *APOE* E4 is a major risk factor for treatment adverse effects known as amyloid-related imaging abnormalities (ARIA)^8^. Understanding the impacts of *APOE* genotype, sex and ancestry on biological processes in the human brain prior to disease onset could be key to developing preventive AD treatments.

AD pathology progressively appears in brain regions in a specific order, with the entorhinal cortex (ERC) being one of the first cortical regions impacted in AD^9,10^. Part of the periallocortex, the ERC serves as the interface between the hippocampus and neocortex, playing a critical role in integrating positional and temporal information for episodic memory formation^11–13^. The anatomical and cytoarchitectural organization of the human ERC is important to understanding the early stages of AD. Molecular characterization of AD pathology in ERC has identified pTau tangles in Layer 2 ERC neurons and revealed AD-associated changes in white matter^9,14,15^.

AD is also associated with dysfunction of several specific cell types^7,16^, including various glial populations. Astrocytes are linked to increased lipid metabolism and brain cholesterol synthesis^16^. Oligodendrocytes are linked to disease via observations of decreased myelination in AD patients and in iPSC models of APOE4 Oligos^16,17^. Microglia mount an inflammatory response in AD characterized by increased expression of triggering receptor expressed on myeloid cells 2 (*TREM2*), which has been hypothesized to trigger the phagocytosis of amyloid beta plaques with supporting evidence from a *Trem2* knockout mice study^16^. Single-nucleus RNA-seq (snRNA-seq) studies in postmortem human prefrontal and entorhinal cortex demonstrate that many AD-related gene expression changes are cell type-specific, with oligodendrocytes showing many transcriptional differences^18, 19^. Studies have also identified changes in cell states in AD, specifically microglia activation and disease-associated astrocytes and oligodendrocytes^20,21^.

Given that specific cell types and cortical layers have been implicated in AD, it is critical to understand the laminar organization of ERC cell type populations in the context of AD risk factors. To investigate spatially-localized cell type vulnerability related to AD risk, we generated paired snRNA-seq and spatially-resolved transcriptomics (SRT) data in postmortem human ERC tissue from middle aged brain donors with no clinical signs of AD. Donors were stratified across *APOE* genotype (E2+ versus E4+ carriers), ancestry (AA versus EA), and sex. We have targeted the cellular and molecular landscape of risk to identify factors that might be targeted before the emergence of clinical illness. We characterized human ERC spatial domains and cell types, focusing on subpopulations of glial cell types. We identified differentially expressed genes (DEGs) between E2+ and E4+ carriers, with most identified in a specific oligodendrocyte subcluster showing a transcriptional signature consistent with a more immature, non-myelinating state. We also found ancestry-specific DEGs between E2+ and E4+ carriers, which mostly did not overlap, hinting at ancestry-specific mechanisms of *APOE* risk contribution to AD.

## 2| Results

### 2.1| Evaluating *APOE* risk in a diverse population

This study was designed to evaluate gene expression changes across cell types and spatial domains mediated by biological risk factors for AD, including allelic variation at the *APOE* locus, ancestry and sex. We generated paired spatially-resolved transcriptomics (SRT) and snRNA-seq data in postmortem human entorhinal cortex (ERC) tissue from adult brain donors with no clinical signs of AD (**Figure 1A, Table S1**). The final analyzed cohort included 30 donors with diverse risk of AD, including reduced risk in E2+ *APOE* carriers (n=14) and increased risk in E4+ *APOE* carriers (n=16, **Figure 1A, Table S1**)^7,22,23^. *APOE* allele carrier groups contained both hetero- and homozygote *APOE* genotypes across 21 male and 9 female donors (**Fig S1A-B**). Age at time of death for the cohort ranged from 30 to 68 years old (**Fig S1C**). Ancestry modifies the effect of *APOE* genotype, and accordingly, our cohort included 14 African ancestry (AA) and 16 European ancestry (EA) donors across *APOE* E2+ and E4+ genotypes^4^ (**Figure 1A, Fig S11B**). Ancestry was reported by legal next-of-kin and confirmed by death records, and DNA genotyping data was used to match predicted ancestry (**Fig S1D**, **Methods: Ancestry inference**). All donors had no clinical AD diagnosis and evidence of only minimal to no AD-associated pathology (**Fig S1E-F**).

**Figure 1.**
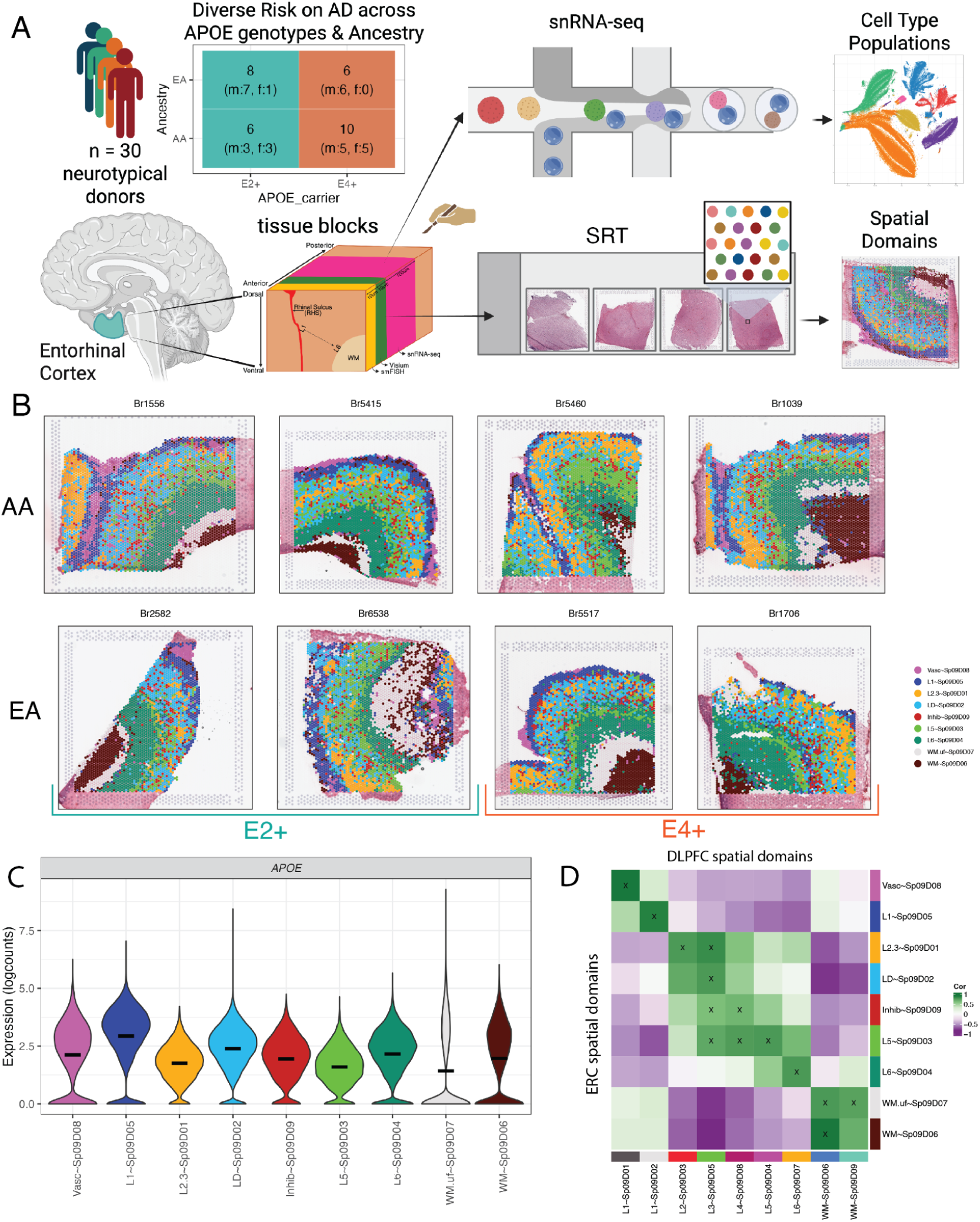
Experimental design and annotation of entorhinal cortex (ERC) spatial domains (SpDs). **A.** Schematic depicting cohort composition (n=30 donors) stratified across *APOE* genotype, sex and ancestry as well as experimental design for generation of single nucleus RNA-seq (snRNA-seq) and spatially-resolved transcriptomics (SRT) data from postmortem human ERC tissue. Created in BioRender. Lab, M. (2025) https://BioRender.com/6dkstuj **B.** Representative SRT tissue sections showing *BayesSpace* clusters at *k*=9 from donors with different *APOE* genotypes (E2+ versus E4+) and ancestries (African, AA and European, EA). **C.** Violin plot of *APOE* log-normalized expression (`logcounts`) across SpDs. **D.** *spatialLIBD* spatial registration heatmap^26–28^ of ERC SpDs compared to reference SpDs in the DLPFC^28^. Summarizes pairwise correlation analysis between enrichment t-statistics from each dataset, high correlation (green) suggests similar patterns of gene expression. High confidence matches (cor > 0.5, merge ratio = 0.1) are marked with an “X” Related to Figure 2, **Fig S1-Fig S14**, **Fig S43**, **Table S1-Table S3**.

### 2.2| Human ERC contains both laminar and non-laminar spatial domains

To characterize the topography of gene expression in the ERC, we generated SRT data using the Visium spatial gene expression platform (10x Genomics) on a single tissue section from each donor (**Table S2**). Following quality control, we obtained a median of 3,964 high-quality spots per sample and a total of 122,742 spots across donors. While ERC samples showed some variation in orientation, patterns of transcriptional activity and expression of layer-associated marker genes confirmed that each tissue section included white matter (WM) and layered gray matter (GM, **Fig S2**, **Fig S3A-C**). Following batch correction using *harmony*^24^ (**Fig S4**, **Fig S5**), we performed spatially aware clustering with *BayesSpace*^25^ and identified transcriptionally distinct spatial domains (SpDs), with *k*=9 being the optimal number of SpDs (**Figure 1B**, **Fig S3D**, **Fig S6**). We observed the expected number of reads, detected genes and mitochondrial rate across SpDs indicating that clustering was not driven by technical factors, but rather accurately represented the organization of gene expression across the ERC architecture (**Fig S7**).

To annotate SpDs, we leveraged anatomical landmarks and data driven marker genes (**Fig S8**, **Table S3**). We characterized the 9 SpDs using our previously developed spatial registration framework^26–28^ by integrating our ERC SRT data with SRT data from the human dorsolateral prefrontal cortex (DLPFC)^28^, and to an ERC snRNA-seq^29^. We also evaluated expression of established human and mouse cortical layer marker genes^27,30,31^, including ERC-specific gene sets in rodents^32,33^. Spatial domains were named with the following syntax: *L*∼Sp_k_D_d_ where *L* was the layer annotation associated to domain *d* at clustering resolution *k*. Vasc∼Sp_9_D_8_ was the outermost layer and had high expression of blood-associated genes (*HBA1*, *HBA2*, *HBB*, **Fig S8**), suggesting that this SpD represented a vasculature meninges layer. Vasc∼Sp_9_D_8_ also showed high correlation with the vascular SpD in human DLPFC, and vascular cell types from Franjic et al. (**Figure 1D, Fig S9**). The next most superficial SpD, L1∼Sp_9_D_5_, had high expression of *AQP4*, a known marker of astrocytes enriched in L1 in ERC and other cortical regions (**Fig S8**, **Figure 1D, Fig S9**). L1∼Sp_9_D_5_ also showed the highest expression of *APOE*, consistent with the well-recognized high expression of *APOE* in astrocytes^34^ (**Figure 1C, Fig S10A**). Middle SpDs, including L2.3∼Sp_9_D_1_, LD∼Sp_9_D_2_, and Inhib∼Sp_9_D_9,_ showed less discrete laminar organization. While L2.3∼Sp_9_D_1_ mapped to L2/3 in DLPFC (**Figure 1D**), a key structural difference between ERC and DLPFC is a low cell density L4 or “laminar desicans” (LD) in ERC. LD∼Sp_9_D_2_ was the most abundant SpD (**Fig S11A**), and given its position between the GM layers, enrichment in low nuclei spots (**Fig S7D**), and weak association with ERC cell types (**Fig S9**), we annotated this SpD as the laminar desicans. Consistent with the notion that lamination of the ERC is less well-defined than that of neocortical regions^35^, ERC SpDs representing L2-4 did not specifically register with histological layers of DLPFC (**Figure 1D**). In contrast to superficial ERC layers, deeper layers annotated as L5∼Sp_9_D_3_ and L6∼Sp_9_D_4_ were more continuously laminar. L5∼Sp_9_D_3_ showed high expression of *PCP4*, a classic L5 marker in human cortex^27,30,31^ (**Fig S8**, **Figure 1D**), as well as *ETV1,* a L5a marker in rodent ERC^32^. We did not see strong evidence for L5 sublayers at this clustering resolution (**Fig S12**). L6∼Sp_9_D_4_ was located between L5∼Sp_9_D_3_ and WM, and registered with DLPFC L6 (**Figure 1D**). Registration to ERC Layer annotated neurons from Franjic et al. supported L2, L5, and L6 identities (**Fig S9**). Finally, we identified an inhibitory neuron SpD (Inhib∼Sp_9_D_9_) with high expression of *GAD1* and *GAD2*, and registration to ERC inhibitory neurons, interspersed throughout the middle GM layers (**Fig S8**, **Fig S9**).

Beyond GM domains, we identified two WM domains. We annotated a more superficial WM U-fiber layer (WM.uf∼Sp_9_D_7_) aligned with known fiber tracts in ERC^36^ and defined by low transcriptional activity (**Fig S7A**). We also defined a deeper WM layer (WM∼Sp_9_D_6_) with relatively higher expression of canonical WM marker genes, such as *MOBP* (**Fig S8**). WM.uf∼Sp_9_D_7_ in ERC spatially registered to the two WM SpDs in the DLPFC, while WM∼Sp_9_D_6_ in ERC matched the superficial DLPFC WM domain, suggesting unique WM organization in the ERC (**Figure 1D**). Compared to mouse ERC^33^, human ERC contained more SpDs and substantially different gene expression in L3 and L5 (**Fig S13**). Having characterized SpDs in ERC, we next examined differences between SpD frequency across *APOE* carrier groups to rule out changes in laminar organization related to AD risk. While all SpDs were represented in all donors, we observed different SpD frequencies across tissue sections, likely due to dissection variation (**Fig S11A**). However, differential proportion analysis with *crumblr*^37^, which utilizes centered log ratios (CLR, a transformation of proportion values) to test for composition differences, did not show significant differences in SpD composition by *APOE* carrier status (**Fig S14B-D**, **Fig S3**, **Table S4**). These data represent the first molecular atlas of the human ERC and define transcriptionally discrete SpDs for downstream differential expression analyses.

### 2.3| Identification of fine resolution ERC single cell clusters and integration with SRT data

To define molecular profiles for ERC cell type populations, we generated snRNA-seq data for each donor (Chromium, 10x Genomics, **Figure 1A, Table S5**). After quality control and batch correction, we performed two rounds of clustering to 1) identify clusters driven by low quality and 2) identify cell type clusters resulting in the characterization of 122,004 high quality nuclei (**Fig S15**-**Fig S25**). Based on literature-derived and data-driven marker gene expression^19,38^, cells were categorized into 38 fine subclusters across 8 broad cell types: astrocytes (subclusters: Astro.1-5), macrophages (Macro), microglia (Micro.1-5), oligodendrocytes (Oligo.1-5), oligodendrocyte precursor cells (OPC.1-5), vascular cells (pericytes (Vasc.PC), endothelial (Vasc.Endo), and vascular leptomeningeal cells (Vasc.VLMC)), plus excitatory (Excit) and inhibitory (Inhib) neurons with 8 and 7 sub-clusters, respectively (**Figure 2A-B**, **Fig S26**, **Fig S27**, **Fig S28**). Inhibitory neuron subclusters were annotated based on expression of specific inhibitory marker genes (*SST*, *VIP*, *LAMP5*, **Fig S27**, **Fig S29A**)^30^ and cluster registration^26–28^ against previously published DLPFC and ERC datasets^29,39^ (**Fig S30**). We identified data-driven *MeanRatio*^40^ marker genes for the fine subclusters to generate molecular profiles for these ERC cell type subclusters (**Fig S28**, **Table S6**).

**Figure 2.**
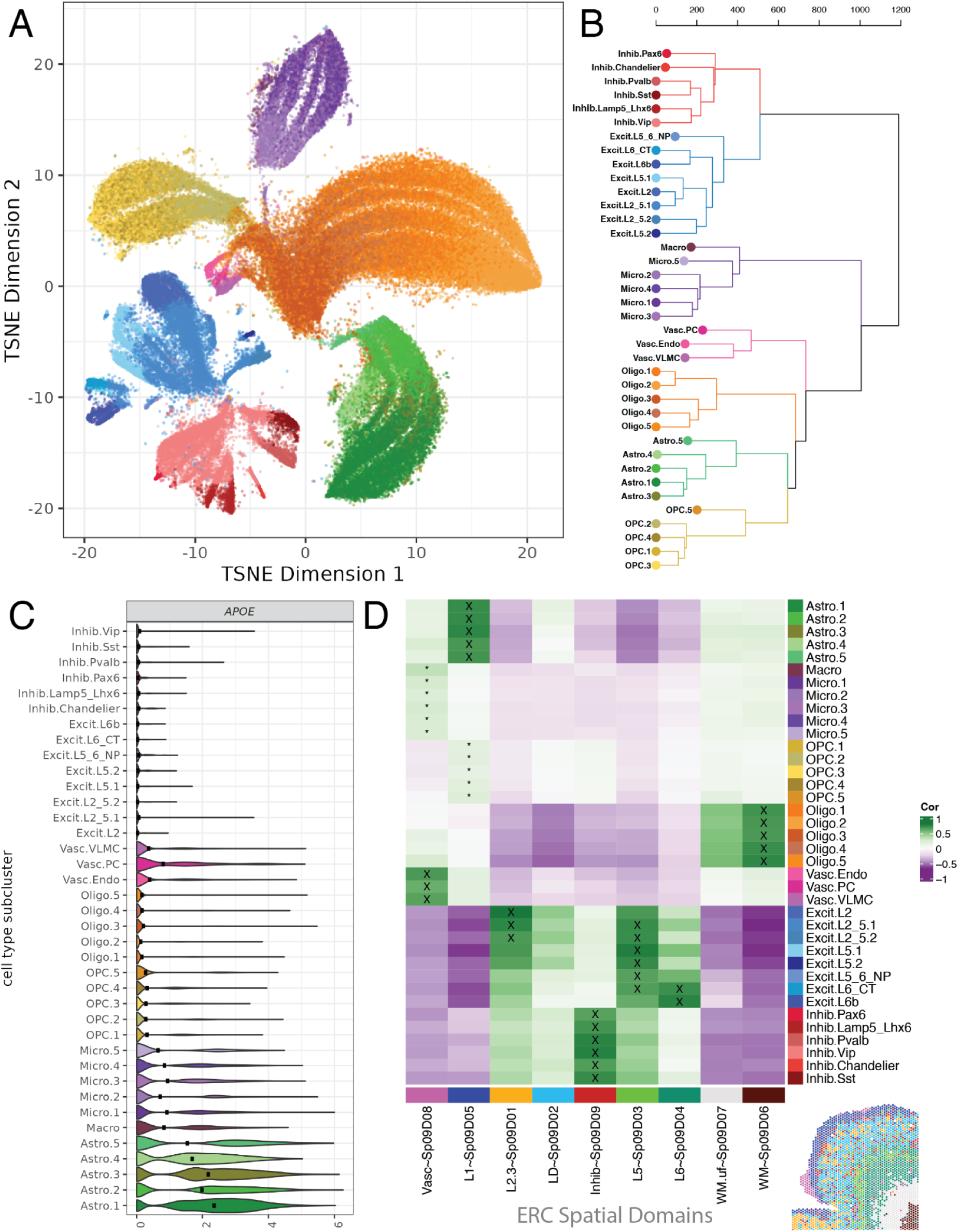
Identification and annotation of fine resolution ERC subclusters in the snRNA-seq data. **A.** *t*-distributed stochastic neighbor embedding (tSNE) plot of 122k nuclei across 38 fine resolution subclusters. **B.** Hierarchical clustering (`method = “ward.D2”`) of fine resolution subclusters. **C.** Violin plot of *APOE* log-normalized expression (`logcounts`) across fine subclusters. **D.** *spatialLIBD* spatial registration heatmap^26–28^ showing the correlation between enrichment *t*-statistics for the ERC SpDs and snRNA-seq subclusters. Inset shows SRT spotplot of SpDs for Br5517 (Figure 1B). High confidence matches (cor > 0.5, merge ratio = 0.1) are marked with an “X” low confidence matches are marked with “*”. Cluster colors in all panels match. Related to Figure 1, **Fig S15**-**Fig S29**, **Fig S43**, **Table S5**.

Spatial registration^26–28^ between ERC snRNA-seq fine subclusters and ERC SRT SpDs revealed cellular organization within ERC cytoarchitecture (**Figure 2D**). This allowed for refined annotation of excitatory neuron subclusters based on their laminar organization, including two populations (Excit.L2_5.1 and Excit.L2_5.2) that mapped to both L2.3∼Sp_9_D_1_ and L5∼Sp_9_D_3_ (**Figure 2D**). Compared to ERC SpDs, all snRNA-seq inhibitory neuron subclusters correlated strongly with Inhib∼Sp_9_D_9_ as expected. Vascular snRNA-seq subclusters mapped to the vascular SpD (Vasc∼Sp_9_D_8_), astrocyte fine subclusters mapped to L1∼Sp_9_D_5_, and all snRNA-seq oligodendrocyte subpopulations mapped to WM∼Sp_9_D_6_ (**Figure 2D**). Macrophage and microglia subclusters had weak matches to Vasc∼Sp_9_D_8_, and likewise for OPC subclusters to L1∼Sp_9_D_5_ (**Figure 2D**). Integration of SRT and snRNA-seq data through spot deconvolution with *RCTD*^41^ further confirmed SRT and cell type annotations, such as increased Astro proportions in L1∼Sp_9_D_5_ (**Fig S32**, **Table S7**).

Next, we examined *APOE* and other known AD risk genes across all subclusters to evaluate the cellular landscape of AD risk. As expected, *APOE* expression was highest in astrocytes, specifically Astro.1 (**Figure 2C**, **Fig S10**), although other cell types, including microglia, macrophage, and pericytes expressed *APOE* at lower levels. While AD-associated cell states have been identified in several glia cell types^20^, none of our Astro, Micro, and Oligo subclusters showed high expression of genes linked to disease-associated cell type states such as *SERPINA3*, *IL33*, and *IL1B* (**Fig S29B-D**). As donors in our cohort did not have AD-related pathology, this provided evidence that these previously reported disease-associated cell type states are not intrinsic to genetic risk for AD (**Fig S29B-D**).

To investigate whether *APOE* carrier status, sex, and age influenced snRNA-seq cell type proportions, we examined subcluster composition across these demographic groups. While Astro and Oligo were the most abundant ERC cell types (**Fig S26A**), all 38 subclusters were detected in at least 19 donors, and 26 (68%) were detected in all donors. Subcluster composition varied across individual donors (**Fig S26C**), but no clusters were specific to one *APOE* carrier group (**Fig S26B**). Next, we performed differential proportion analysis with *crumblr*^37^, over *APOE* carrier status, sex, and age (**Fig S33**, **Fig S34**, **Table S8**, **Methods: Differential proportion analysis**). *APOE* E4+ carriers showed a decrease in frequency in Oligo.1 and Oligo.2 compared to E2+ carriers (**Fig S33**). In female donors, Oligo.5 and Oligo.3 subclusters had significantly increased frequency compared to males (**Fig S34**). Age-associated proportion differences were identified in Oligo.5, with no significant interaction with *APOE* carrier groups (**Fig S34**). Overall, Oligo frequency was the most variable over *APOE* carrier status, sex, and age, suggesting changes in ERC cell composition could be a factor in AD risk with implications for early changes in AD.

### 2.4| *APOE* carrier status influences gene expression in spatially-defined white matter and vascular domains

To understand the impact of *APOE* carrier status on spatial gene expression in the ERC, we performed differential gene expression analyses of SRT data followed by Gene Ontology (GO) overrepresentation analysis in E4+ versus E2+ carriers adjusting for ancestry and sex (**Fig S35**, **Fig S36, Table S9**). The highest number of differentially expressed genes (DEGs, FDR < 0.05, 22 upregulated and 8 downregulated) was localized to Vasc∼Sp_9_D_8_, which contained Astro, Oligo, and Vasc cells based on spot deconvolution analyses (**Fig S32**). These DEGs included downregulation of *CNTN2* and *MAL*, genes implicated in oligodendrocyte differentiation (**Fig S36D**, **Figure 3G**). In line with findings that WM may be most susceptible to early AD-related pathology,^15,42^ the U-fiber WM SpD (WM.uf∼Sp_9_D_7_) had the second most DEGs (**Figure 3A-C**). Specific to WM.uf∼Sp_9_D_7_, we observed upregulation of the *MAPT* paralogue *MAP2* which has known functions in modulating tau pathology (**Figure 3D**)^43^. Notably, *MAPT* is an AD risk gene encoding the microtubule-associated protein tau^44^. Myelination and axon ensheathment GO terms were common in downregulated genes across WM.uf∼Sp_9_D_7_, Vasc∼Sp_9_D_8_, and L6∼Sp_9_D_4_ (**Figure 3G, Fig S36A-B, Table S10**).

**Figure 3.**
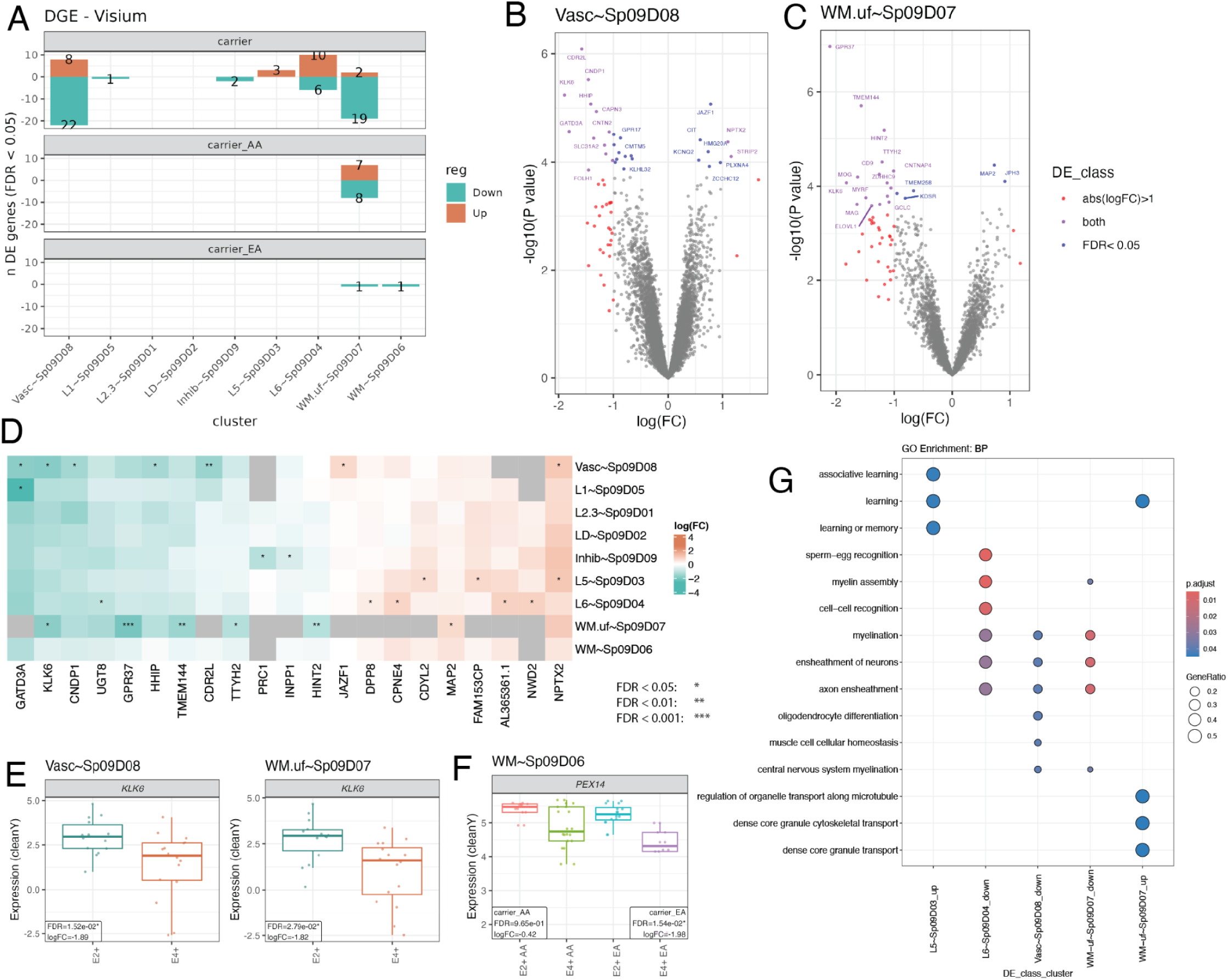
*APOE* carrier status interacts with ancestry to impact gene expression in distinct SpDs. **A.** Number of differentially expressed genes (DEGs, FDR<0.05) identified with differential gene expression (DGE) analysis, either up or downregulated in *APOE* E4+ carriers relative to E2+ carriers for the full dataset (carrier) as well as for ancestry-specific analyses (carrier_AA, carrier_EA). **B-C.** Volcano plots for carrier DE analysis for Vasc∼Sp_9_D_8_ (**B**) and WM.uf∼Sp_9_D_7_ (**C**) genes with FDR<0.05 or absolute log_2_ fold change (logFC) > 1 are indicated with point color. **D.** logFC heatmap for up to top 5 DEGs in each SpD. **E.** Log-normalized gene expression boxplots with model adjustment covariates removed (*cleanY*)^123^ for *KLK6,* comparing E2+ and E4+ carriers in Vasc∼Sp_9_D_8_ and WM.uf∼Sp_9_D_7_. **F.** *cleanY* expression boxplot of *PEX14* in WM.uf∼Sp_9_D_7_ comparing E2+ and E4+ carriers over ancestry groups. **E** and **F** show the FDR-adjusted p-value and logFC comparing E2+ and E4+. **G.** *clusterProfiler*^127^ dot plot of top biological process (BP) gene ontology (GO) terms for the E4+ versus E2+ DEGs. Related to **Fig S35**, **Fig S37**, **Fig S38**, **Table S9**, **Table S10**.

While most DEGs were unique to one SpD, we identified several shared DEGs across multiple SpDs, several of which were linked to AD (**Figure 3D**). For example, *NPTX2*, a gene involved in memory and learning, was significantly upregulated in both Vasc∼Sp_9_D_8_ and L5∼Sp_9_D_3_ (**Figure 3D**). *KLK6*, a gene linked to dipeptidase activity and oligodendrocyte differentiation (**Fig S36**), was downregulated in E4+ in WM.uf∼Sp_9_D_7_ and Vasc∼Sp_9_D_8,_ (**Figure 3D-E**). KLK6 is a serine protease expressed primarily by oligodendrocytes and is downregulated in the temporal cortex and hippocampus of AD patients^45,46^. Notably, KLK6 has been proposed as a biomarker for AD given that AD patients have higher KLK6 levels in cerebrospinal fluid^47^. Other genes related to dipeptidase activity (*CAPN3*, *FOLH1*, *CNDP1*) were also downregulated in Vasc∼Sp_9_D_8_ (**Figure 3D, Fig S36A**). Taken together, we observed distinct gene expression changes between E2+ and E4+ carriers across SpDs, implicating different biological processes across ERC layers.

To directly investigate how ancestry and sex modify *APOE* risk, we identified ancestry-specific and sex-specific DEGs across E2+ and E4+ carriers. For ancestry-specific analyses, we identified 15 DEGs in WM.uf∼Sp_9_D_7_ for AA, while EA donors only had 2 downregulated DEGs (1 in each WM SpD, **Fig S37A**). In WM∼Sp_9_D_6_, *PEX14* downregulation was observed in EA, but not AA donors (**Figure 3A+F, Fig S37D, Methods: Differential expression analysis**). *PEX14* encodes a protein marker for peroxisomes, organelles involved in lipid homeostasis. Downregulation of *PEX14* is consistent with peroxisome density decreases in the ERC as AD pathology increases^48^. Ancestry-specific E4+ downregulation of *PEX14* suggests reduction of peroxisomes may be more pronounced in E4+ EA individuals. We next examined risk difference between males and females, and in total identified 91 male DEGs in WM.uf∼Sp_9_D_7_, and 1 female DEG, *PAX6*, in WM∼Sp_9_D_6_ (**Fig S38A+D+G**). The larger number of DEGs in males was likely influenced by the uneven representation of sexes across the cohort (21/30 male; 70%). Similar to the overall DEG results (**Table S10**), genes downregulated in male WM∼Sp_9_D_6_, such as *MAL*, *KLK6*, and *CD9,* were associated with myelination. In summary, we found that *APOE* carrier status has the largest impact on gene expression in WM and vasculature SpDs with differences across ancestry and sex.

### 2.5| *APOE* E4+ is associated with altered transcription in oligodendrocytes

AD causes diverse changes across cell types^7,16^. Given the limited cell type resolution of SRT data, we next leveraged our paired snRNA-seq data to perform differential expression followed by GO overrepresentation analysis between *APOE* carrier groups. We compared E2+ vs. E4+ carriers at both broad cell type and fine subcluster resolution. At broad resolution, Oligo had the most DEGs, followed by Astro, Excit and Inhib neurons (**Figure 4A-B**, **Fig S39**, **Table S11**). We observed very few DEGs in other broad cell types, including microglia. DEGs were often cell type-specific, although some genes, such as *NPTXR,* were upregulated in multiple cell types (i.e. Astro, Oligo, and Excit (**Figure 4C**)) in E4+ compared to E2+ carriers. Genes upregulated in Astro in E4+ carriers, such as *NETO1* and *PLPPR4,* were associated with synaptic membrane GO terms (**Table S12**, **Figure 4C**). In Oligo, upregulated genes in E4+ carriers, such as *ANXA1,* were associated with extracellular matrix and external encapsulating structure (**Figure 4B**, **Fig S40A+C**). We also observed upregulation of calcium channel genes in E4+ Oligo (**Fig S40B+D**). Likewise, *CD69* had the largest fold change between E4+ and E2+ Oligos (**Figure 4F**).

**Figure 4.**
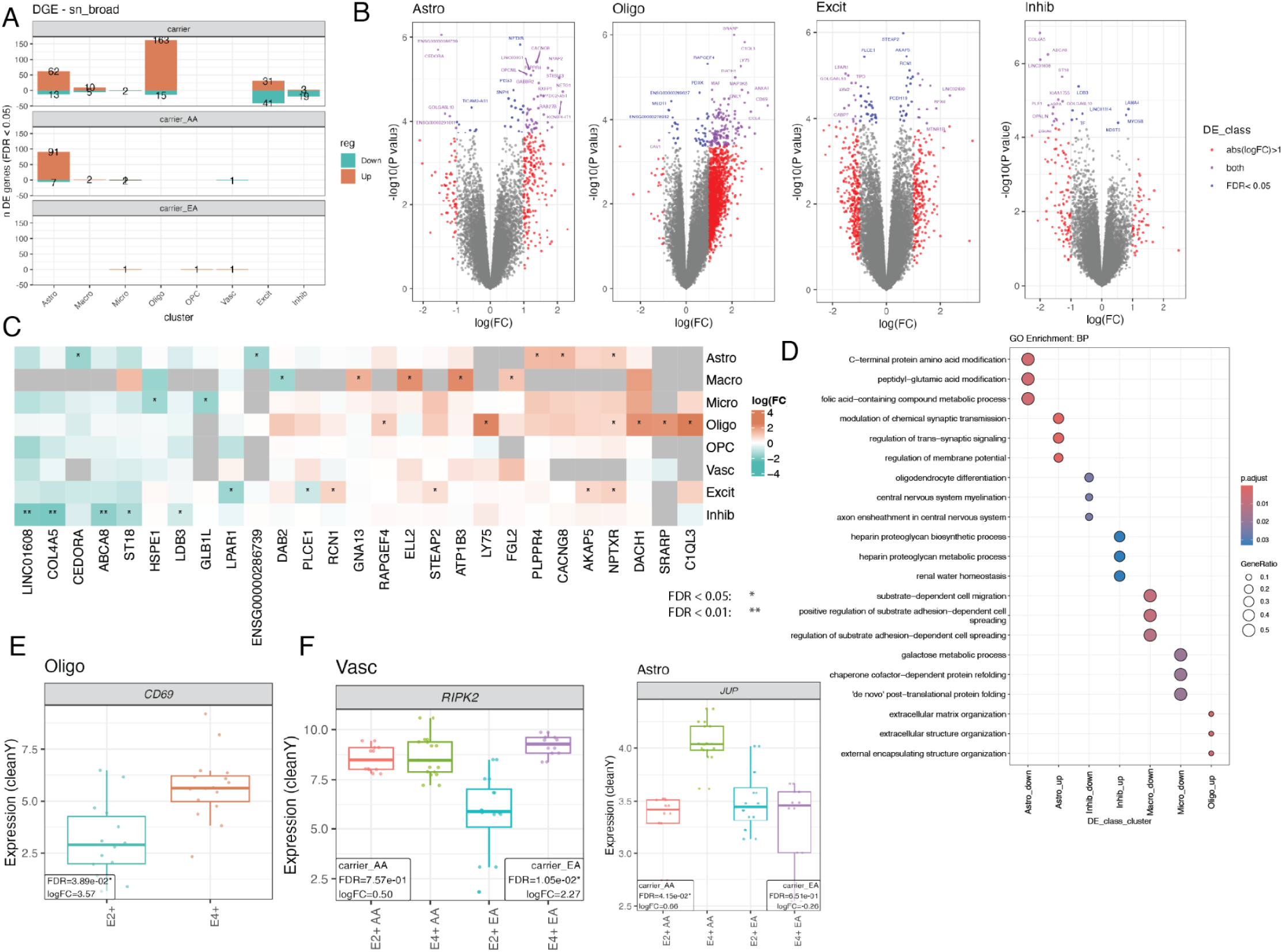
*APOE* carrier status impacts cell type gene expression predominantly in oligodendrocytes. **A.** Bar plot of number of FDR<0.05 DE genes, either up or downregulated in *APOE* E4+ carriers relative to E2+ carriers for the full dataset analysis (carrier) and ancestry-specific analyses (carrier_AA, carrier_EA). **B.** Volcano plot for carrier DE analysis for astrocytes (Astro), oligodendrocytes (Oligo), excitatory (Excit), and inhibitory (Inhib) neurons. Genes with FDR<0.05 or absolute logFC > 1 are indicated with point color. **C.** logFC heatmap for up to top 5 DEGs per broad cell type across all broad cell types. **D.** *clusterProfiler*^127^ dot plot of top biological process (BP) gene ontology (GO) terms for carrier model Visium DEGs. Gene ratio is the number of genes in each gene set annotated in a term (y-axis) over the total in the respective gene set. p.adjust is the FDR-adjusted p-value. **E-F.** *cleanY* gene expression boxplots for *C1QL3* in Oligo (**E**) and *RIPK2* in Vasc and *JUP* in Astro over carrier and ancestry (**F**). *E* and *F* show the FDR-adjusted p-value and logFC comparing the two *APOE* carrier groups. Related to **Fig S37**, **Fig S38**, **Fig S39**, **Table S11**, **Table S12**.

Similar to SRT data, we also examined ancestry- and sex-specific DEGs by cell type across *APOE* E2+ and E4+ carriers. Ancestry-specific analysis identified 98 DEGs in Astro in AA donors (**Figure 4A**, **Fig S37B**), with *JUP* upregulated in E4+ carriers with a large difference between EA and AA donors (**Figure 4, Fig S37H**). *RIPK2* was uniquely downregulated in Vasc of EA donors compared to AA donors (**Figure 4F).** Sex-specific DEGs were detected mostly in males (948 DEGs), with the majority observed in Oligo and Excit. Only 2 DEGs were detected in females (**Fig S38B+E+H**, **Methods: Differential expression analysis**), likely due to reduced sample size. In summary, DE analysis at the broad cell type level was consistent with DE analysis in SRT data, with most DEGs identified in glia-enriched SpDs (WM.uf∼Sp_9_D_7_ and Vasc∼Sp_9_D_8_) and Oligos.

### 2.6| Fine resolution snRNA-seq DEGs concentrated in an oligodendrocyte subcluster

Given that broad cell types contain a diversity of highly heterogeneous subtypes, we next investigated differential gene expression in our fine resolution snRNA-seq subclusters between *APOE* E2+ and E4+ carriers. The majority of DEGs were found in oligodendrocyte subcluster Oligo.3 (**Figure 5A, Fig S41**, **Table S13**, 679 upregulated and 343 downregulated genes, FDR<0.05). Downregulated genes in Oligo.3 were implicated in myelination and oligodendrocyte differentiation, such as *PLP1*, *MAG*, *MAL*, *MBP*, *SOX10*, and *OPALIN* (**Figure 5B**, **Fig S42A**, **Table S14**). The gene encoding tau, *MAPT*, is expressed in oligodendrocytes and has a significant developmental role in oligodendrocyte maturation and myelin synthesis^49^. *MAPT* was downregulated in Oligo.3 (**Table S13**), suggesting a link between AD risk and impaired oligodendrocyte differentiation. Upregulated Oligo.3 DEGs included *FOS*, *TLR2*, *STAT1*, and *STAT4*, which are all involved in response to calcium ion (**Figure 5B, Fig S42B, Table S13**). *TLR2*, *STAT1*, and *STAT4* are also associated with inflammation, and may suggest an interferon response^50^. Specifically, *TLR2* promotes a pro-inflammatory feed forward loop that causes secondary inflammation in AD pathogenesis^51^. This suggests that in E4+ carriers, Oligo.3 may exist in an immune-reactive state. Oligo.3 cells also had significant upregulation of *LRRK2*, a gene implicated in Parkinson’s disease (PD)^52^, which is of interest given commonalities between PD and AD ^53^. OPCs showed the highest *LRRK2* expression (viewable through our *iSEE*-powered app^54^), which is consistent with reports showing *LRRK2* knockdown affects OPC to oligodendrocyte maturation^55^.

**Figure 5.**
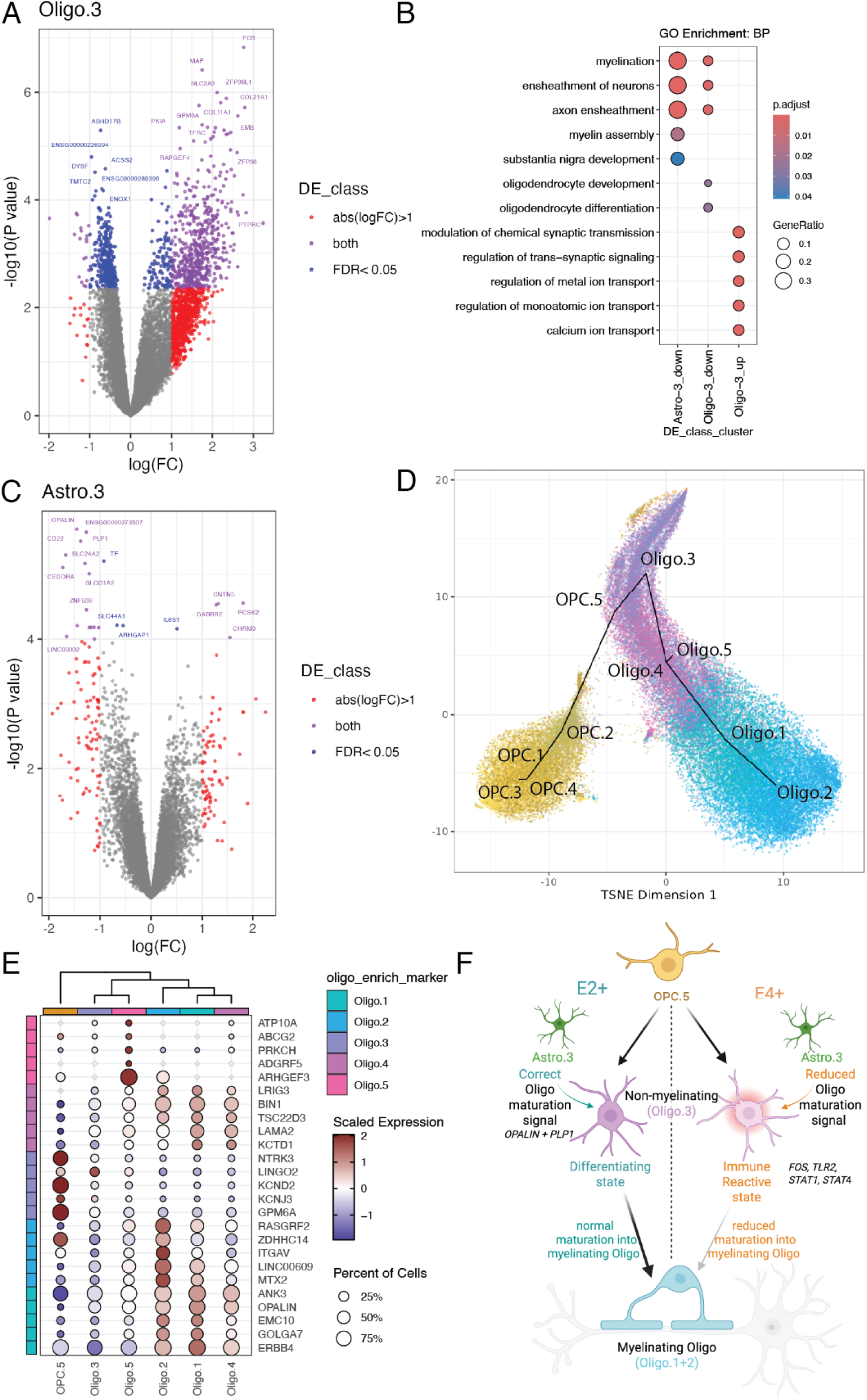
Fine subcluster differential expression shows interruption of Oligo maturation. **A.** Volcano plots of Oligo.3, genes with FDR<0.05 or absolute logFC > 1 are color highlighted. **B.** *clusterProfiler*^127^ dotplot of gene ontology (GO) term enrichment for Oligo.3 and Astro.3. C. Volcano plot of Astro.3 (similar to *A*). tSNE plot of OPC and Oligo subclusters with trajectory analysis (**Methods: Oligodendrocyte subcluster trajectory analysis**). **B.** Centered and scaled log-normalized gene expression *scDotPlot*^136^ of Oligo subcluster marker genes. **F.** Proposed model for oligodendrocyte subcluster classification and maturation trajectory in *APOE* E2+ and E4+ carriers. OPC.5 is the transcriptionally closest OPC subcluster to Oligo.3, which may receive oligo maturation signals from Astro.3 in the *APOE* E2+ carrier background. In E2+, Oligo.3 is in a differentiating state and can mature into myelinating oligodendrocytes (Oligo.1 and 2). In an *APOE* E4+ carrier background, Oligo.3 may have reduced maturation signals and appears in an immune-reactive state leading to altered maturation into myelinating oligodendrocytes. Created in BioRender https://BioRender.com/1gocr2n. Related to **Fig S29**, **Fig S37**, **Fig S38**, **Fig S41**, **Fig S43**, **Fig S46**, **Fig S47**, **Table S13**, **Table S14**.

Next, we examined DEGs in other subclusters, with both Astro.3 downregulated and Micro.4 upregulated in E4+ DEGs presenting common GO overrepresented terms with Oligo.3 DEGs down and upregulated in E4+, respectively (**Fig S44**). Astrocyte subcluster Astro.3 shared many downregulated DEGs with Oligo.3 (**Figure 5C**, **Table S13**), and also showed GO overrepresentation hits for myelination terms (**Figure 5D+E, Fig S42A**). Notably, *OPALIN* was the most significant downregulated gene in Astro.3 (**Figure 5E**). Generally, oligodendrocytes express very low levels of *APOE*, therefore, the *APOE* effect in the Oligo.3 subcluster may be due to effects in cell type in which *APOE* is more highly expressed, such as astrocytes (**Figure 2**)^7^. Communication between two or more cell types could lead to coordinated responses in gene expression, which could lead to common sets of DEGs^56–58^. For instance, DEGs downregulated in Inhib in E4+ carriers included *PLP1* and *OPALIN*, which were also downregulated DEGs in Oligo.3 (**Figure 4B+D**, **Fig S42A**).

To better characterize cellular interactions between Oligo.3 and other fine resolution ERC subclusters, we performed cell to cell communication (CCC) analysis with *LIANA+*^59^ to identify recurrently used ligand-receptor (LR) pairs between a specific pair of subclusters (LR pairs observed in at least 20 donors). Unsurprisingly, communication among neuronal subclusters accounted for the most LR pairs in commonly observed interactions (**Fig S43A, Table S16**). Oligo.3 was either the source (n=115) or the target (n=111) for many recurrent LR interactions. Oligo.3 cells frequently communicated with several neuronal populations including Excit.L2, while also communicating with itself (**Fig S43B**). Common LR interaction genes and subcluster DEGs only overlapped in Oligo.3, involving 11 LR interactions communicating between Oligo.3 and OPC, Oligo, or neuronal subclusters (**Fig S43C)**. DEG-involved L-R interactions were most spatially likely in gray matter domains (**Fig S43D**). Overall, altered gene expression in Oligo.3 may impact communication with linked cell types across ERC and within specific local microenvironments.

Identification of Oligo.3 as the primary subcluster implicated in *APOE* E4+ risk prompted further investigation into the molecular identity of all 5 Oligo subclusters. Trajectory analysis revealed that Oligo.3 was most closely related to oligodendrocyte precursor cells (OPC), suggesting Oligo.3 represented a more immature cell state compared with Oligo.1 and Oligo.2, the most mature Oligo subclusters (**Figure 5D**, **Fig S29, Methods: Oligodendrocyte subcluster trajectory analysis**). Analysis of marker genes within Oligo subclusters revealed that Oligo.1 and Oligo.2 highly express marker genes associated with mature myelinating Oligos, such as *OPALIN* and *MOG* (**Figure 5E**, **Fig S29D**). Top Oligo.3 enrichment marker genes, including *LINGO2, GPM6A*, and *KCNJ3*, were implicated in neuronal communication. Many Oligo.3 marker genes were shared with OPC.5, supporting the close relationship between these subclusters identified by trajectory analysis (**Figure 5D-E**). Following Oligo.3 in the trajectory analysis, Oligo.4 also showed marker genes involved in neuronal communication and had higher expression of myelination genes, although lower than that of Oligo.1 and Oligo.2 (**Figure 5E**, **Fig S29D-E**). In comparison to other Oligo subclusters, Oligo.5 had higher expression of vascular-associated genes, such as *ARHGEF3*, suggesting this population may represent perivascular oligodendrocytes^20^ (**Figure 5E)**. Overall, trajectory and marker gene analyses support the identification of Oligo.3 as a non-myelinating oligodendrocyte population. Taken together with differential expression analyses, our data support a model in which large differences in transcription in Oligo.3 between E2+ and E4+ carriers could distinguish different cell states associated with AD risk. In E2+ carriers, Oligo.3 could be primed to differentiate into mature myelinating oligodendrocyte populations, while in E4+ carriers, Oligo.3 may be in an inflammatory state with reduced maturation into myelinating oligodendrocytes (**Figure 5F**).

Next, we evaluated the impact of ancestry and sex on Oligo.3 gene expression across E2+ and E4+ carriers. Both AA and EA donors showed ancestry-specific DEGs associated with *APOE* carrier status in Oligo.3 (**Fig S37B**), with some genes regulated in opposite directions (**Fig S37E+H**). Oligo.3 upregulated genes in E4+ AA donors were associated with modulation of chemical synaptic transmission, neurotransmitter transport and secretion, and calcium ion transport. On the other hand, upregulated genes in E4+ EA donors were associated with muscle tissue development and extracellular matrix organization (**Fig S45**). In particular, Oligo.3 showed upregulation in E4+ EA donors of *SORBS1*, *ADRA1B*, and *ATP1A2,* genes implicated in the cellular component GO term caveola. Caveolae are bulb-shaped pits in the plasma membrane implicated in active intracellular signal transduction supporting cell survival signaling^60^, and they have been previously proposed as a target for AD treatment^61^. In terms of sex, Oligo.3 sex-specific DEGs were only detected in males (1,830 DEGs, **Fig S38C**). This may reflect oligodendrocyte transcriptional changes associated with AD observed only in males^18,62^, but interpretation is limited by the lower number of female donors in our study. Some Oligo.3 DEGs in males, such as *PRKCH* and *ITPA3* (**Fig S38F+I**), showed expression changes trending in an opposite direction from that of females, suggesting diverging sex-specific changes associated with *APOE* carrier status. In summary, *APOE* E4+ carriers show differential gene expression in a specific non-myelinating oligodendrocyte subtype suggesting a differentiating state in E2+ donors compared to an immune reactive state in E4+ donors, with individual DEGs and their functions showing differences across donors with EA and AA ancestries.

### 2.7| Detection of oligodendrocyte risk signatures in AD

Previous snRNA-seq studies in AD donors report differential expression in oligodendrocytes^17,19^. To determine whether AD risk signatures overlap with DEGs associated with AD diagnosis, we compared our data to two publicly available postmortem human snRNA-seq datasets including AD donors–one in the ERC (Grubman et al.)^19^ and one in the prefrontal cortex (Blanchard et al.)^17^. First, we performed gene set enrichment analysis between our fine resolution subcluster DEGs and cell type DEGs from Grubman et al., which included 6 AD donors and 6 controls, across a range of *APOE* genotypes^19^ (**Table S15**). We observed significant overlap (FDR < 0.05) between downregulated Oligo DEGs in AD donors and downregulated Oligo.3 DEGs in *APOE* E4+ carriers, including a shared decrease in *PLP1* and *OPALIN*. However, there was no enrichment between upregulated Oligo DEGs in AD donors and upregulated Oligo.3 DEGs in *APOE* E4+ carriers (**Fig S46A**, **Table S15**). There was significant enrichment between Grubman et al. downregulated genes in Oligo and AA downregulated genes in Oligo.3 (**Fig S47A**). Grubman et al. also examined DEGs between AD and control donors in six Oligo subclusters^19^. Our Oligo.3 subcluster had the highest correlation with their Oligo o4 subcluster, which was present in both AD and control donors (**Fig S46B**)^19^. Oligo o4 showed upregulation of genes enriched in synapse organization, calcium signaling, and metal ion transport^19^, similar to GO terms identified for upregulated Oligo.3 DEGs in E4+ carriers (**Figure 5E**).

Next we compared our snRNA-seq DEGs to cell type DEGs from Blanchard et al., which analyzed gene expression differences between E3/E3 against E3/E4 and E4/E4 Oligos in the prefrontal cortex (PFC) of 20 AD and 12 control donors^17^. We found significant enrichment between our Oligo.3 downregulated DEGs in E4+ carriers and their AD downregulated Oligo genes in E3/E4 vs. E3/E3 carriers, again including *OPALIN* and *PLP1* (**Fig S46C, Table S15**). We also observed opposite significant enrichment between Oligo.3 E4+ upregulated DEGs and several sets of DEGs identified by Blanchard et al.: downregulated Oligo DEGs identified in E3/E4 vs. E3/E3 in donors with no AD, DEGs from E4+ vs. E3/E3 in all donors, and induced pluripotent stem cells (iPSC)-derived E4/E4 vs. E3/E3 Oligos (**Fig S46C**). Ancestry-specific Oligo.3 AA DEGs also showed opposite significant enrichment (**Fig S47**). Congruently, Oligo.3 E4+ upregulated DEGs were enriched with upregulated DEGs in iPSC-derived Oligos^17^ (**Fig S46C**). Blanchard et al. highlighted that genes involved in cholesterol homeostasis were upregulated in Oligos between *APOE* genotypes, while myelination-related genes were downregulated^17^. In contrast, we did not see differential expression of cholesterol-related genes in Oligo.3, but we did replicate downregulation of myelination-related genes (*PLP1*, *MAG*, *PLLP*, and *OPALIN*, **Fig S46D**). While not significant, the same trend was observed in ancestry-specific DEGs (**Fig S47C**).

Together, our study in *APOE* E4+ carriers and two previously published studies in AD donors consistently show downregulation of oligodendrocyte differentiation and myelination genes, such as *PLP1* and *OPALIN*, and their corresponding GO terms^17,19^. While myelination and oligodendrocyte differentiation have been previously implicated in AD pathogenesis^17^, we link these processes to changes in a specific oligodendrocyte population in *APOE* E4+ carriers of middle age with no clinical signs of AD.

### 2.8| Association between *APOE* carriers and AD pathology linked to Oligo.3

To identify common gene expression signatures between SRT and snRNA-seq data, we used a Multi-Omics Factor Analysis (MOFA) framework to define coordinated gene expression patterns^63–65^, or MOFA factors, across the datasets. Each MOFA factor assigns contribution weights to expressed genes independently for each input gene expression view (either a SRT SpD or a snRNA-seq subcluster)^63–65^, thus summarizing coordinated expression across spatially-defined local cell communities and cell types. MOFA factors are then compressed into a latent space such that each donor has one score for each MOFA factor^63–65^. The donor-level MOFA factor scores can then be tested for statistical association with donor-level covariates^65^. The gene-level MOFA factor weights provide information about the contribution from each gene to the MOFA factor within each SRT SpD or snRNA-seq subcluster. The genes with the highest and lowest gene-level MOFA factor weights provide a signature pattern for the factor, and these signature genes provide a biological functional interpretation for the factor. Overall, MOFA factors link coordinated expression patterns to jointly explain differences between donors. We calculated 7 MOFA factors from both snRNA-seq subclusters and SRT SPDs, then tested for factor associations with *APOE* carrier status (E2+ or E4+), ancestry, age, sex. We also considered several measures of AD pathology, including Braak stage and CERAD score, since MOFA examines factor weights on a per donor basis (**Figure 6A**, **Table S17**, **Methods: Multicellular factor analysis**). We note that only eighteen donors had available Braak staging as middle age donors with no clinical AD symptoms are not routinely screened for AD pathology. To overcome this limitation, we performed immunostaining for pTau in adjacent ERC tissue sections from each donor (**Fig S1E**). Overall levels of pathology increased with age, but did not separate by *APOE* carrier status (**Fig S1F**).

**Figure 6.**
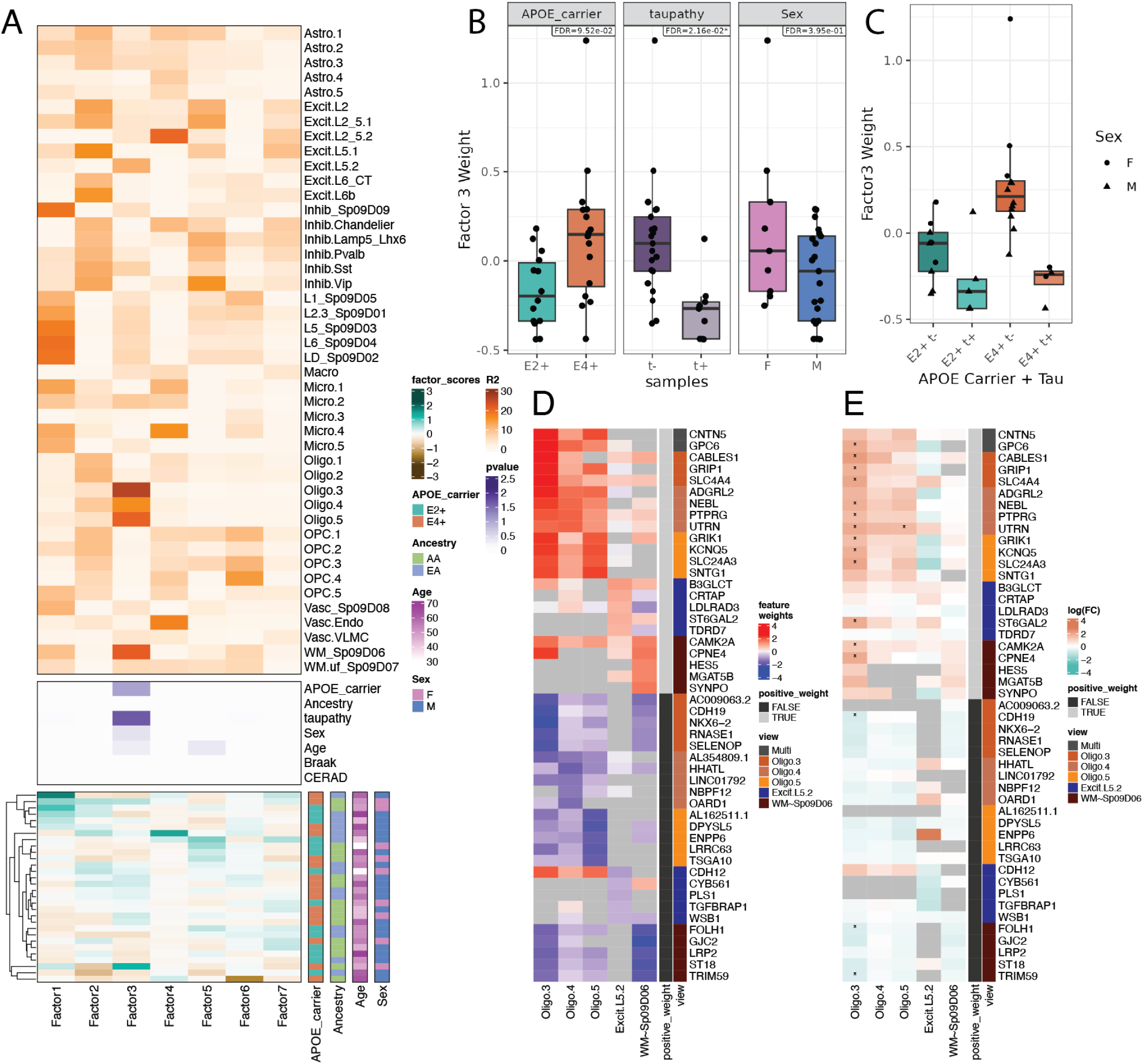
Common gene expression signature across oligodendrocytes and white matter associated with *APOE* carrier status and tau pathology. **A.** Multi-Omics Factor Analysis (MOFA) *MOFAcelullaR*^65^ heatmap. Input gene expression views (SRT SpDs and snRNA-seq subclusters) are shown in the top orange section rows, with colors representing the gene expression variance explained (R^2^) in each input gene expression view by the donor-level MOFA factor scores (columns). The purple heatmap in the middle shows the p-values for the association between donor-level MOFA factor scores and donor-level covariates (rows). The bottom heatmap shows all 30 donors (rows) and their MOFA factor scores annotated with donor-level covariates. **B.** Boxplot of MOFA factor 3 donor-level scores over *APOE* carrier status and presence of pTau pathology in ERC (**Methods: pTau immunostaining**), and sex. **C.** Boxplot of MOFA factor 3 donor-level scores over *APOE* carrier status combined with presence of pTau pathology (t-, tau negative; t+, tau positive). **D.** Heatmap of MOFA factor 3 gene-level weights for top 5 signature genes for MOFA factor 3 across select snRNA-seq clusters or SRT SpDs (gene expression `view`) where this MOFA factor explains a high percent of the gene expression variance. A negative factor gene-level weight indicates a negative association of the gene with the factor score. Annotation bars show views where the gene has a top gene-level weight for MOFA factor 3 (`view` annotation), and whether factor 3 has a positive gene-level weight in this view (`positive_weight` annotation). **E**. Heatmap of *APOE* carrier differential expression logFC for the top MOFA factor 3 signature genes shown in **D**. Annotation bars are the same as **D** with stars denoting DEGs between E2+ and E4+ carriers. Related to **Table S17**.

We identified that MOFA factor 3 donor-level scores were associated with *APOE* carrier status and pTau pathology (**Figure 6A**). Factor 3 explained a high percent of gene expression variance in WM∼Sp_9_D_6_ in SRT data and Oligo.3, Oligo.4, Oligo.5 (Oligo.3-5), and Excit.L5.2 in snRNA-seq data (**Figure 6A**). Higher factor 3 scores were nominally associated with E4+ vs. E2+ (p = 0.014, FDR = 0.095, **Figure 6B**) and female vs. male donors (p = 0.056, FDR = 0.395, **Figure 6B**). Surprisingly, MOFA factor 3 donor-level scores also had a significant negative association (FDR< 0.05) with pTau scores in adjacent ERC tissue sections (**Figure 6B**), indicating that genes implicated in this coordinated gene expression pattern were associated with both AD risk and pTau levels. A combined category of *APOE* carrier status and pTau detection showed highest factor 3 donor-level scores for E4+ and no detection of pTau (**Figure 6C**).

To biologically interpret MOFA factor 3, we next examined its signature genes in WM∼Sp_9_D_6_ and Oligo.3-5 (**Figure 6D-E**). The top five MOFA factor 3 signature genes that had high gene-level weights in WM∼Sp_9_D_6_ or Oligo.3-5 (**Figure 6D**) overlapped with upregulated E4+ DEGs in Oligo.3 (**Figure 6E**), although not all these genes were Oligo.3 upregulated E4+ DEGs, or WM∼Sp_9_D_6_ or Oligo.4-5 upregulated E4+ DEGs. Likewise, the top five MOFA factor 3 signature genes with the lowest gene-level weights had lower expression in E4+ in Oligo.3-5 and WM∼Sp_9_D_6_, with only three of these genes being E4+ downregulated DEGs in Oligo.3 (**Figure 6D-E**). MOFA factor 3 composed a coordinated expression pattern in Oligo.3-5 in the spatially defined WM∼Sp_9_D_6_ cell community, and this coordinated expression pattern explained differences between *APOE* carriers and observed pTau pathology in ERC. The MOFA framework identified signature genes that supported the independent DEG analysis results in Oligo.3. At the same time, MOFA identified additional genes contributing to *APOE* risk changes in other Oligo subclusters and WM∼Sp_9_D_6_, such as *SYNPO, HES5*, and *MGAT5B* (**Figure 6D**). These genes were not expressed in Oligo.3-5, but were expressed in WM∼Sp_9_D_6_ (**Figure 6D**), and have been previously implicated in AD and oligodendrocyte maturation^66–71^. The MOFA framework thus independently supported DGE results, while also recovering complementary smaller expression changes missed by DGE analyses. In summary, MOFA provided further evidence for non-myelinating oligodendrocytes (Oligo.3-5 subclusters, **Figure 5**) as vulnerable in *APOE* E4+ carriers and identified additional genes previously implicated in AD in these cell types and associated WM.

## 3| Discussion

Here we characterized the spatio-molecular landscape of epidemiological AD risk factors (*APOE* carrier status, ancestry, and sex) at cellular resolution in the neurotypical human ERC by generating paired SRT and snRNA-seq data from middle age donors. Our study focuses on the molecular biology of risk prior to pathology associated with clinical dementia. We annotated nine spatial domains (SpDs) and 38 cellular subclusters in the SRT and snRNA-seq data, respectively. The organization of SpDs reflected the structure of the periallocortex, highlighting reduced laminar organization compared to neocortical structures such as the DLPFC^35^. Unique ERC SpDs included LD∼Sp_9_D_2_, with decreased cellular density, and WM.uf∼Sp_9_D_7_, a fibertrack WM region. In the paired snRNA-seq data, we defined glial subclusters including five oligodendrocytes subclusters that map to different stages of oligodendrocyte maturation. This data provides a resource for understanding the spatio-molecular organization of the human ERC, and is explorable through interactive web application: SRT data via *spatialLIBD*^26^ and snRNA-seq data through *iSEE*^54^. Analysis of *APOE*-dependent risk revealed the most transcriptomic differences in glia-enriched SpDs (WM.uf∼Sp_9_D_7_ and Vasc∼Sp_9_D_8_) and a non-myelinating oligodendrocyte sub-cluster (Oligo.3), including ancestry-specific changes.

SRT facilitated localization of *APOE*-dependent changes across the spatial topography of the ERC. We found the highest number of *APOE*-dependent DEGs in vasculature and WM domains (Vasc∼Sp_9_D_8,_ and WM.uf∼Sp_9_D_7_). These data provide evidence that transcriptomic changes may occur early on in WM in individuals with high AD risk, which is consistent with changes in WM preceding accumulation of plaques and tangles in animal models^15^. Changes in Vasc∼Sp_9_D_8_ could reflect *APOE* E4+ dysfunction in the blood brain barrier, which has been noted in pre-clinical AD^72^. Downregulated DEGs in WM and vasculature in *APOE* E4+ carriers were also enriched for myelination-related GO terms, which may reflect early changes leading to reduced or altered myelination, which has been observed in the WM of AD patients, particularly in the ERC^15^. *NPTX2* upregulation in Vasc∼Sp_9_D_8_ and L5∼Sp_9_D_3_ is counter to findings of *NPTX2* downregulation in AD^73^, but this discrepancy could reflect evolving changes or compensation as a factor of risk, early pathology, and more advanced AD. SRT studies in human ERC tissue from AD donors could clarify how disease progression impacts these changes. Differential expression of both *NPTX2* and its receiver *NPTXR* in different SpDs and cell types could be evidence of dysregulation of ligand-receptor pairs.

While SRT data highlighted transcriptomic changes associated with vasculature in *APOE* E4+ carriers, we did not detect many DEGs associated with vasculature by snRNA-seq (**Fig S41**). This discrepancy is likely due to insufficient power (∼53 vascular cells/donor, **Fig S26A**)^74^. Previous imaging studies found that neurofibrillary tangles were concentrated in L2 during AD progression, and AD mouse models highlighted L2 neuron vulnerability^9,14^. We did not observe DEGs in L2.3∼Sp_9_D_1_ and only 7 total DEGs in excitatory L2 subclusters (**Fig S41**). Among other neuronal SpDs, a limited number of DEGs (21 total) were identified in Inhib∼Sp_9_D_9_, L5∼Sp_9_D_3_, and L6∼Sp_9_D_4_ (**Figure 3A**), suggesting that *APOE* risk has a limited effect in ERC GM layers. For neuronal cell types, we detected moderate signal at the broad cell type level (**Figure 4A**), which only resulted in a few DEGs at the fine subcluster resolution (**Fig S41**), likely due to the small number of cells per donor for these subclusters (**Fig S26A**)^74^ and the potentially limited *APOE* impact on ERC neurons. Differences between snRNA-seq and SRT results likely reflect the fact that snRNA-seq measures only nuclear transcripts, while SRT measures both cytosolic and neuropil transcripts. Our paired dataset provides a more comprehensive assessment of gene expression changes across spatial compartments and fine resolution subclusters.

Fine cell type subclusters provided a high resolution view of cellular diversity in the ERC, allowing for detection of *APOE* associated changes in Oligo.3. Multiple findings pointed to an association between *APOE* E4+ carriers and impaired oligodendrocyte differentiation, which supports a recent hypothesis that a vulnerable pre-myelinating oligodendrocyte contributes to AD^20^. Gene expression in Oligo.3 suggests that this sub-cluster is non-myelinating, combined with trajectory analysis this could suggest a pre-myelinating identity, although further work is needed to profile oligodendrocyte maturation process in the human ERC. Our DGE analyses support the hypothesis that maturation is stalled in E4+ Oligo.3, which could explain the reduced proportions of myelinating Oligo.1 and Oligo.2 observed in E4+ donors through cell type frequency analyses. Corresponding increases in OPC.5 could also be linked to this disruption, perhaps from increased OPC recruitment in response to inadequate numbers of myelinating oligocytes (**Fig S33B-E**). Reduction of myelinating Oligo populations could disrupt myelination, which may be related to the first signs of neurodegeneration and/or AD pathogenesis ^75^. Our cell type frequency changes in E4+ carriers are in line with observations of increased frequency of OPCs in an AD mouse model and decreased Oligos in postmortem AD patients ^76^, although AD-associated frequency changes in oligodendrocytes are inconsistent across the literature^15,77^. Although *crumblr* attempts to control for confounding factors^37^, cell type frequency analyses in snRNA-seq data can be biased, and validation with alternative assays would provide further support for these observations. Furthermore, while our data suggests alteration of oligodendrocyte maturation, we cannot exclude the possibility that Oligo.3 cells mediate risk via direct interactions with neurons and glia independent of a role in myelination.

Since expression of *APOE* in Oligo.3 is low, the mechanism by which *APOE* allelic variation impacts this cell type is likely indirect. High expression of *APOE* in Astros and observations that *APOE* E4+ disrupts lipid transport and oligodendrocyte differentiation via Astro-Oligo signaling suggests the involvement of astrocytes^78,79^. Supporting the potential of astrocytes mediating this mechanism, DEGs in Astro.3 pointed to disruption of similar biological processes as in Oligo.3. Follow up work in cellular models will be necessary to elucidate the causal origin of *APOE* genotype affecting Oligo.3 gene expression. For example, impaired signaling between Astro-Oligo could be directly investigated in human iPSC-derived co-cultures derived from E2+ and E4+ donors ^17^.

A unique aspect of our study design was assessing the impact of ancestry and sex on gene expression associated with *APOE* carrier status. We found large numbers of ancestry and sex-specific differences in Oligo.3 DEGs, suggesting that these factors could modulate the observed disruption in Oligo biology. Due to limitations in the number of female donors in our cohort, larger studies with more females will be needed to confirm sex-specific transcriptional changes. However, previous studies have reported male specific changes in Oligos from AD donors^18,62^, providing evidence for sex-specific differences associated with illness state that may also be related to risk. We also found profound effects of ancestry on Oligo.3 gene expression across *APOE* carrier status. Given the comparable number of AA and EA donors, we were well powered to detect these findings. GO overrepresentation analysis of ancestry-specific Oligo.3 DEGs highlighted different functional pathways. For instance, only EA donors had E4+ upregulated genes involved in extracellular matrix organization, which is likely linked to the association of *APOE* E4+ with blood-brain barrier (BBB) dysfunction that predicts cognitive decline^72^. Moreover, *APOE* E4+ and amyloid beta have been found to modify microstructural abnormalities associated with BBB breakdown in the ERC in a predominantly EA cohort^80^. Our study provides the first evidence that there are interactions between ancestry and *APOE* risk in cell types and SpDs of the ERC that may contribute to the reduced risk effect of *APOE* E4+ in AA compared with EA^4^. Future studies should aim to replicate these findings in the ERC or other brain regions in a larger donor cohort carefully controlled for sex, ancestry, and *APOE* carrier status.

When comparing gene expression changes associated with AD risk in ERC to previously published snRNA-seq datasets in other brain regions from patients with AD, we encountered both commonalities, such as E4+ downregulated oligodendrocyte DEGs, and contrasting findings, such as Oligo.3 E4+ upregulated genes enriched in downregulated DEG sets from other cell types^17,19^. Discrepancies could be due to differences in 1) statistical methods, 2) demographics between donors, 3) brain region (e.g. PFC vs. ERC), 4) sample size, and 5) presence of AD pathology. Specifically regarding statistical methods used, previous AD DGE results were computed using effective cell-level gene marker finding methods^17,81^, instead of pseudobulk DGE methods that allow adjusting for sample-level covariates^74^, batch effects^37^, cell type heteroscedasticity^82^, high frequency of zeros in expression data^83^, or identify differential detection^84^. An analysis of previously published snRNA-seq AD datasets^17–19,38^ combining a common oligodendrocyte subcluster annotation with the latest sc/snRNA-seq DGE statistical methods could be useful to validate differences in oligodendrocytes across brain regions, AD risk in additional neurotypical donors, and changes due to the presence of AD.

Using cell-to-cell communication (CCC) analysis, we identified ligand-receptor (LR) pairs involving DEGs associated with *APOE* carrier status. However, we only observed LR pairs involving DEGs in Oligo.3, potentially due to the larger number of DEGs in this subcluster (**Fig S43C**). Overall, Oligo.3 was implicated as both a source and target across different LR pairs (115 vs 111), and frequently communicated with Excit.L2 and other neuronal subclusters along with itself (**Fig S43B**). Some Oligo.3 LR pairs involving DEGs could be involved in communication cascades leading to further changes in Oligo.3 between *APOE* carrier groups. While we had hypothesized that CCC results might link *APOE-*expressing astrocytes to Oligos and show overlap with Oligo.3 DEGs, no such genes were identified. Further research will be needed to determine whether *APOE* allele changes originate in Oligo.3, or if Oligo.3 is the most susceptible to alterations in cell to cell communication from other *APOE*-expressing cell types.

Multi-Omics Factor Analysis (MOFA)^63–65^ identified factor 3, which represented a coordinated expression pattern between non-myelinating Oligos (Oligo.3-5) and WM∼Sp_9_D_6_. Interestingly, pTau detection in the ERC had a strong negative association with MOFA factor 3. On the other hand, MOFA factor 3 supported a positive association between *APOE* risk and E4+ upregulated gene expression changes in Oligo.3-5 (snRNA-seq) and WM (SRT), while additionally implicating genes not found through DGE. MOFA identified genes associated with AD risk only expressed in WM∼Sp_9_D_6_ that were not detected in Oligo.3-5, including: *SYNPO,* which was recently linked to AD genome wide association study using UK Biobank data^66^*; HES5*, which is important for neural stem cell homeostasis^67–69^ and was identified as a DEG in 1-month-old mice from a triple transgenic mouse model of AD; and *MGAT5B* whose deficiency decreases astrocyte activation and increases oligodendrocyte maturation in demyelinating mouse brain model^70,71^. MOFA also detected *CPNE4*, an upregulated E4+ DEG expressed in both WM∼Sp_9_D_6_ and Oligo.3. *CPNE4* has a copy number variant associated with AD and is a hippocampus CA3 AD DEG^85,86^. These findings suggest that transcriptomic differences in Oligo.3 and other non-myelinating oligodendrocytes may precede the onset of AD pathology in high risk individuals, and further transcriptional differences occur at the onset of pTau deposition. Supporting this interpretation, Blanchard et al.^17^ showed greater downregulation of myelination genes in donors without AD pathology relative to positive AD pathology. Our work provides important justification for future studies to investigate this relationship using iPSC-derived co-cultures of cortical neurons and oligodendrocytes^17^, as well as assembloid models to interrogate oligodendrocyte function^87^ in the context of AD risk and resilience.

While this study was primarily focused on understanding *APOE* E4+ risk and E2+ resilience in the context of AD, it is important to note that *APOE* E4+ has also been associated with increased risk of cognitive decline in Parkinson’s disease (PD)^88,89^. Interestingly, we found upregulation of the PD-associated gene *LRRK2* in Oligo.3^52^, highlighting this oligodendrocyte subcluster as potentially also relevant for PD. *LRRK2* has varied roles across tissues and cell types, including cytoskeletal function, membrane trafficking, subcellular localization and ability to phosphorylate members of the family of Rab-GTPases^90^. In the brain, *LRRK2* is most highly expressed in OPCs^91^, although the literature exploring *LRRK2* expression in the human brain is scarce. *LRRK2* expression has been associated with oligodendrocyte differentiation, with evidence that *LRRK2* knockdown prevents differentiation and subsequent oligodendrocyte maturation^55,92^. Pre-myelinating oligodendrocytes have also been recently linked to PD genetic risk^93^, supporting a potential broader role for Oligo.3 in neurodegenerative diseases. Our results suggest that oligodendrocyte differentiation could be a key process by which *APOE* E4+ and *LRRK2* interact with potential impact on both PD and AD.

In the current study, we examined molecular changes in the ERC, a brain region involved in the first stages of AD^9,10^. However, AD pathogenesis involves multiple brain regions^94^. For example, norepinephrine (NE) neurons in the locus coeruleus (LC) send neuronal projections to the ERC forming a unidirectional LC-ERC circuit that is an initial site of degeneration in AD^95^. The molecular changes that contribute to AD progression and disrupted functional connectivity in the LC-ERC circuit are not understood. The MOFA framework allows integrating data from different assays performed on the same set of donors, even if not all genes are expressed in the different input gene expression views^65^. This makes the MOFA framework ideal for integrating gene expression data from different brain regions that almost by definition, do not express the same sets of genes.

In conclusion, we used integrated SRT and snRNA-seq approaches to define the cellular and molecular anatomical organization of the human ERC in a unique cohort of donors stratified by *APOE* carrier status, sex, and ancestry. We found that transcriptional changes associated with increased AD risk converged on a non-myelinating oligodendrocyte population, with differences in dysregulation between African and European ancestry. Our findings suggest that AD risk in the ERC may be associated with disrupted oligodendrocyte function and changes in myelination, potentially contributing to future neurodegeneration.

## Supporting information

Supplementary Tables 1-17 (small ones)

## Acknowledgements

We thank Amy Deep-Soboslay and James Tooke for their work in sample curation and clinical characterization at LIBD. We thank Geo Pertea for maintaining and imputing the LIBD DNA genotype data. We gratefully acknowledge use of the facilities at the Joint High Performance Computing Exchange (JHPCE) in the Department of Biostatistics, Johns Hopkins Bloomberg School of Public Health that have contributed to the research results reported within this paper. We thank Alison M. Goate (Mount Sinai), Brady J. Maher (LIBD), and Alejandra I. Romero-Morales (LIBD) for constructive feedback on the manuscript preparation. We thank the physicians and staff at the brain donation sites and the generosity of donor families for supporting our research efforts. We thank Hao Zhang and Joseph Margolick from the Flow Cytometry Cell Sorting Core Facility at Bloomberg School of Public Health, Johns Hopkins University (JHU) for doing FACS sorting. The FACS sorting facility was supported by 1S10OD016315-01,1S10RR13777001, and in part by CFAR: 5P30AI094189-04. We thank the Johns Hopkins University Single Cell and Transcriptomics Sequencing Core for single nucleus RNA and SRT sequencing. Finally, we thank the families of Connie and Stephen Lieber and Milton and Tamar Maltz for their generous support of this work.

## Author Contributions Statement

Conceptualization: LAHM, DRW, KM, MR, KRM, LCT

Data curation: LAHM, HRD, RAM, SH, LCT

Formal analysis: LAHM, HRD, NJE

Funding acquisition: DRW

Investigation: HRD, SVB, MRV, RAB, RZ, SCP, KRM

Methodology: SCP, KRM

Project administration: KM, KRM, LCT

Resources: JEK, TMH, DRW

Software: LAHM, NJE, BM

Supervision: DRW, KM, MR, KRM, LCT

Validation: HRD, RZ

Visualization: LAHM, HRD

Writing – original draft: LAHM, HRD, SVB, MRV, NJE, RZ, JRE, MGP, SH, SCP, KRM, LCT

Writing – review & editing: LAHM, JRE, MGP, SCP, DRW, KM, MR, KRM, LCT

## Data Availability

Raw data generated as a part of this study have been deposited on the Gene Expression Omnibus (GEO) with accession numbers GSE307990 and GSE308007. We also created two web applications to help explore this data and can be found on https://research.libd.org/LFF_spatial_ERC/#interactive-websites. Processed data files (R objects) used to make the interactive apps (they only have `logcounts`) can be found on https://research.libd.org/globus/ under the heading `jhpce#LFF_ERC`. R objects with both the `counts` and `logcounts` can be downloaded through the `fetch_data()` function from *spatialLIBD* v1.23.1 or newer^26^.

## Code Availability

All source code for this project is available at https://github.com/LieberInstitute/LFF_spatial_ERC^96^ under the MIT License and permanently archived on Zenodo^96^. The GitHub repository includes log files with specific software versions used for each analysis^96^.

## Materials and Methods

### Tissue

#### Post-mortem human tissue samples

Postmortem human brain tissue from control donors of European or African ancestry with allelic variation at the *APOE* locus (E2/E2, E2/E3, E3/E4, and E4/E4) were obtained at the time of autopsy following informed consent from legal next-of-kin, through the Maryland Department of Health IRB protocol #12–24 and the County of Santa Clara Medical Examiner-Coroner Office in San Jose, CA, under the WCG protocol #20111080. Additional samples were consented through the National Institute of Mental Health Intramural Research Program under NIH protocol #90-M-0142, and were acquired by LIBD via material transfer agreement. Using a standardized strategy, all donors were subjected to clinical characterization and diagnosis. Macroscopic and microscopic neuropathological examinations were performed, and subjects with evidence of significant neuropathology were excluded. Additional details regarding tissue acquisition, processing, dissection, clinical characterization, diagnoses, neuropathological examination, RNA extraction and quality control (QC) measures have been previously published^97^. Demographic information for all 31 donors initially dissected is listed in **Table S1**, **Fig S1**. Sex expression quality control later led us to drop 1 sample recorded as an *APOE* E3/E4 AA male, which was actually an E3/E4 AA female whose age at time of death was 17 years, and thus the sample could not be included in other analyses downstream of clustering (**Fig S25, Methods: Donor demographics check**). African Ancestry donors in this study are of sub-Saharan African Ancestry. Each tissue block was dissected from frozen coronal slabs at the level of the amygdala or the hippocampus. Using a hand-held dental drill, tissue blocks of approximately 1 X 1.5 mm were dissected across the collateral sulcus. Tissue blocks were stored in sealed cryogenic bags at −80°C until cryosectioning.

#### Tissue processing and anatomical validation

To anatomically validate the inclusion of ERC in tissue blocks, we performed multiplexed single-molecule fluorescence in situ hybridization (smFISH) on cryosections. Briefly, 10 μm thick sections were cut on a Leica Biosystems cryostat and mounted onto glass slides. Molecular signatures for distinct ERC cortical layers were probed to distinguish ERC from adjacent cortical regions. Specifically, we targeted three layer-selective markers — *RELN* (layer 1), *FREM3* (layer 3), and *TRABD2A* (layer 5) as well as *MBP* to delineate white matter. smFISH was performed using the RNAScope platform^98^, following the manufacturer’s instructions for the Fluorescent Multiplex Kit v2 and the 4-plex Ancillary Kit (catalog nos. 323100, 323120, Advanced Cell Diagnostics). Tissue sections were fixed in 10% neutral buffered formalin (catalog no. HT501128, Sigma-Aldrich) for 30 min at room temperature, followed by sequential dehydration in 50%, 70%, and twice in 100% ethanol. Sections were then pretreated with hydrogen peroxide for 10 min and incubated with Protease IV for 30 min at room temperature. RNAScope probes were applied as follows: *RELN* (Cat No. 413051-C2), *FREM3* (Cat No. 829021-C4), *TRABD2A* (Cat No. 532881-C1), and *MBP* (Cat No. 411051-C3), all from Advanced Cell Diagnostics. Probe hybridization was performed according to manufacturer protocols. After hybridization, tissue sections were stored overnight in a 4X saline-sodium citrate buffer (catalog no. 10128-690, VWR). Signal amplification was carried out (AMP 1-3), followed by labeling with Opal Dyes 570, 620, 690 and 520 (catalog nos. FP1488001KT, FP1495001KT, FP1497001KT, FP1487001KT, Akoya Biosciences; all at 1:500 dilution). Sections were counterstained with DAPI (4′,6-diamidino-2-phenylindole) for 20 seconds to visualize nuclei, and cover slipped using Fluoromount-G mounting medium (catalog no. 0100-01, Southern Biotechnology). Slides were stored and protected from light for at least 24 hours prior to imaging. Imaging was performed on a Nikon AXR confocal microscope with NIS-Elements software, using a Nikon APO Lambda D 20x/0.80 or APO LAmbda S 40X/1.25 objective for high-resolution analysis. For anatomical quality control, whole-slide imaging was performed in a single plane using a Nikon Plan APO Lambda D 2X/0.1 objective.

#### pTau immunostaining

Additional 10 μm sections were cryosectioned to detect the presence of phosphorylated tau (pTau) accumulation in the ERC. RNAScope in situ hybridization combined with immunofluorescence (RNAScope-IF) staining was performed targeting pTau (n=1 tissue section per donor). Prior to fixation, 10% Neutral Buffered Formalin solution (NBF, Cat no. HT501128, Sigma-Aldrich) was chilled for 1 hour at 4°C. Slides were incubated in the prechilled 10% NBF for 15 min at 4°C. After fixation, tissue sections were processed according to the manufacturer’s protocol using the RNAScope Fluorescent Multiplex Kit V2 (Cat no. 323110, ACD). Samples underwent a graded ethanol dehydration series (50%, 70% and twice in 100% ethanol) at room temperature (RT). Following the dehydration step, a hydrophobic barrier was applied using a barrier pen (Cat. no. H-4000, Vector Laboratories) to ensure that all subsequent regents fully covered the tissue sections. The samples were then pretreated with hydrogen peroxide for 10 min at RT. Next, samples were incubated overnight at 4°C with primary antibody mouse anti-p-Tau (Ser202/Thr205)-AT8 (Cat no. MN1020B, Invitrogen) diluted 1:20 in antibody diluent. After incubation, slides were washed and post-fixed in 10% NBF for 30 mins at RT, followed by protease treatment for 30 min at RT to optimize probe accessibility. Sections were then incubated with a mixture of two RNAScope probes: *MBP* (Cat no. 411051, Advanced Cell Diagnostics, Hayward, California) and *SNAP25* (Cat no. 518851-C3, Advanced Cell Diagnostics) for 2 hours at 40°C in a humidified chamber. Following hybridization, samples were washed five times with RNAScope wash buffer at RT. Signal amplification was performed using the RNAScope AMP 1-3 reagents, followed by fluorescence probe labeling with Opal Dye 520 (Cat. no. FP1487001K, Akoya Biosciences) and Opal dye 690 (Cat. no. FP1498001K, Akoya Biosciences). After probe labeling, samples were incubated with secondary antibody Donkey anti-mouse IgG-Alexa 555 (Cat no. A31570, Invitrogen) at a 1:500 dilution for 40 min at RT in the dark.

Following secondary antibody incubation, samples were counterstained with DAPI and mounted using Flouromount-G (Cat no. 01001-01, Southern Biotech), then stored in the dark overnight to dry before imaging. Slides were imaged using a Nikon AXR confocal microscope system with NIS-Elements imaging software vAR6.10.01, utilizing a Nikon APO lambda D 20x/0.8 objective and/or a Nikon APO lambda S 40x/1.25 objective. All lambda stacks were acquired using identical imaging settings and laser power. For the 20x objective, tile images were captured to encompass cortical layers L1 through white matter (WM). After image acquisition, all 31 samples were accessed for the pTau signal. Samples were considered pTau-positive if three or more discrete pTau signals were identified per section by visual inspection by two independent experimenters blinded to sample ID. Of the total 31 samples, 9 were classified as pTau-positive, and 22 as pTau-negative according to this criterion **Table S1**.

#### Visium H&E data generation

Tissue sections (10 μm thick) were prepared in a cryostat and mounted directly onto Visium Gene Expression Slides (10x Genomics). One section from each donor was placed within a single Visium capture area with experimenter blinded to sample ID, resulting in a total of 31 samples. Sample composition within an individual slide was organized across demographic variables (*APOE* genotype, sex, ancestry) to mitigate bias to the extent possible (**Table S2**). Sections were fixed in methanol and stained with Hematoxylin and Eosin (H&E) following the manufacturer’s protocol (User Guide CG000160 Rev C). H&E stained slides were imaged on a Leica CS2 equipped with a 20x objective and 2x doubler, prior to downstream molecular processing to facilitate subsequent tissue alignment. After imaging, tissue sections were processed using the Visium Spatial Gene Expression protocol (User Guide CG000239 Rev D). Briefly, tissue was permeabilized (optimized at 18 mins based on empirical testing with a representative sample), enabling release of mRNA for subsequent reverse transcription. Synthesized cDNA was then enzymatically removed from the slide and used to construct spatial transcriptomic libraries according to the manufacturer’s instructions. Libraries underwent quality control assessment and were sequenced on the Illumina Novaseq 6000 platform at the Johns Hopkins Single Cell Transcriptomics Core, following the manufacturer’s specifications, with a minimum sequencing depth of 60,000 reads per spot.

#### Ancestry inference

Genotype calling was performed using several Illumina BeadChip DNA microarrays. Using *Plink* v1.9^99^ we excluded variants with a minor allele frequency (MAF) less than 0.5%, a missingness rate of 5% or more, and a Hardy-Weinberg equilibrium (HWE) p<1×10-5. Genotype phasing and imputation were performed using the NHLBI NIH TOPMed imputation service with the GRCh38 TOPMed reference panel^100^. We finally filtered the imputed genotype data to keep only variants having MAF > 0.01.

We inferred local and global ancestry in our study samples using *FLARE* (Fast Local Ancestry Estimation)^101^, which extends the Li and Stephens haplotype copying model within a hidden Markov framework^102^. Study sample genotypes were first phased and missing genotypes were imputed with *Beagle*^103^. We used three reference populations from the 1000 Genomes Project^104^: YRI (African), CHB (East Asian), and CEU (European) (**Fig S1D)**. *FLARE* was run separately for each chromosome using the phased study genotypes, phased reference haplotypes, a GRCh38 genetic map, a minimum minor allele frequency threshold of 0.01 in the reference VCF (`min-maf=0.01`), and other default parameters.

#### Donor demographics check

After clustering and annotation of the snRNA-seq and Visium data types, it was discovered that one of the included donors (Br1289) was incompatible with the study design (**Fig S25**). This identified expression of female sex-specific genes on donor Br1289 who is a male donor (**Fig S25**). Upon further verification, a typo in the case donor selection table was discovered: it is actually Br1285. As this donor (Br1285) had an age of 17 years at time of death which is outside the age range used for this study (**Table S1**, **Fig S1**), it was dropped from further downstream analyses. The data from Br1285 is still valid and could be used for other purposes outside the *APOE* carrier analyses described in this manuscript. For example, Br1285’s data (labeled as Br1289 in tables, figures, and files) is still useful for SpD or snRNA-seq cell type identification.

### Visium

#### Visium raw data processing

Sample slide images were first processed using *VistoSeg*^105^. *VistoSeg* was used to divide the Visium sample slides into individual capture areas using the `VistoSeg::splitSlide()` function. This function takes one large image from the entire slide and separates them into individual capture areas, each capture area on the slide, labeled as A1, B1, C1 and D1. These individual capture area images were used as one of the inputs for *SpaceRanger*^106^.

Individual capture area images from *VistoSeg* were spatially aligned to Visium Slide capture areas using the *Loupe Browser* (version 7.0.1, 10x Genomics). This alignment enabled precise correspondence between histological features and individual capture spots on the slide.

The Visium FASTQ and image data were pre-processed with the 10x *SpaceRanger* pipeline v2.1.0^106^, using human reference genome GRCh38 and 2020-A annotation. Out of 154,752 total spots, 122,742 were in-tissue, with a median 3,989 spots a sample (min=2,396, max=4,880). Counts were detected for 32,006 genes. In-tissue spots have a median of 3,327 UMIs, and a median of 1,573 detected genes (**Table S2**).

#### Visium data processing and quality control

Quality control metrics were calculated with *spatialLIBD* v1.17.0^26^ `add_qc_metrics()`. For quality control manual annotation was used to identify 143 spots on tissue pieces separate from the main tissue section, and small tails of tissue on the edges of the sections. In tandem, `localOutliers()` from *SpotSweeper* v0.99.5^107^ was applied to identify local outliers in three quality control metrics: low library size, low detected gene count, and high rate of mitochondrial gene expression. The 403 spots (0.3%) that were local outliers in one or more metrics were discarded. In total 544 spots were dropped during quality control, leaving 122,202 total in-tissue spots with a median of 3,964 per sample (min=2,367, max = 4,869). H&E images, total number of UMIs, and known marker genes were visualized with spotplots to verify the data quality (**Fig S3A-C**).

#### Visium spatial clustering

### Feature selection

Spatially variable genes were identified for each sample with *nnSVG* v1.8.0^108^. A set of 1,454 genes that passed `filter_genes()` in 5 or more samples, and had an average rank of < 1,500, was selected as the spatially variable genes (SVGs) for this dataset.

A set of layer marker genes from spatial domains, associated with cortical layers was accessed with spatialLIBD `fetch_data(type = “spatialDLPFC_Visium_modeling_results”),` selecting the top 100 enrichment genes for each spatial domain^28^. 441 of these genes had already been detected as SVGs. The union of these two gene sets, n = 1,765 genes, was selected for dimension reduction and clustering.

### Dimension reduction

Principal component analysis was run using the union set of SVG and layer markers, with *scater* v1.30.1^109^. Batch correction was performed with *harmony* v1.2.0^24^, using `RunHarmony(group.by.vars = sample_id)` on the PCs. Visual inspection of the *harmony-*corrected PCs showed a reduction of batch effects by Visium slide or sample effects (**Fig S4**, **Fig S5**).

### Clustering

Visium spots were clustered into spatial domains (SpD) using *BayesSpace* v1.11.0, a spatially aware clustering method^25^, using the function `spatialCluster(nrep = 20000)`on a range of *k* (equivalent to *q* for *BayesSpace* notation) values from 2 to 28 in increments of 1. The optimal value *k*=9 was determined from the maximum negative log likelihood values predicted by `qTune()` from *BayesSpace* (**Figure 1B, Fig S6**). The total number of spots for each Sp_9_D and their proportion across each donor was checked to ensure that all Sp_9_Ds were present in all samples (**Fig S11**).

#### Visium spatial domain annotation & analysis

*MeanRatio* marker genes were found for each SpD using get_mean_ratio() from *DeconvoBuddies* v0.99.10^40^ (**Fig S8**, **Table S3**).

Gene expression modeling was run on the Sp_9_Ds using registration_wrapper() from *spatialLIBD* v1.21.3, which pseudobulks the data by sample and SpD then runs anova, pairwise, and enrichment modeling^26,27^.

To compare the ERC SpDs, the modeling results were compared to other Visium datasets from the human brain with cortical layer annotations^28^. Specifically from *k*=9 *BayesSpace*^25^ spatial clustering that most closely reflected the classic cortical layers (Sp_9_D in DLPFC, **Figure 1D**). SpDs were also compared to Franjic et al. fine subcluster populations subset to ERC (**Fig S9**)^29^.

#### AD risk gene set

We visualized expression of 23 *ClinVar* genes in the snRNA-seq and Visium data (**Fig S10**)^110^. https://platform.opentargets.org/disease/MONDO_0004975/associations Alzheimer’s disease-associated gene list was downloaded on May 21, 2025.

#### Visium nuclei segmentation

To quantify the number of nuclei per spot, first `spaceranger segment` from *SpaceRanger* v4.0.1^106^ was run on each Visium sample’s H&E image to produce nuclear segmentations in `.geojson` format. Reading in spot coordinates from `spaceranger count` and nuclear segmentations for each Visium sample, a custom *Python* workflow we developed was used to quantify nuclear counts within each Visium spot^96^. In particular, spots and nuclear segmentations were imported as `GeoDataFrame`s using *GeoPandas* v1.0.1^111^, and a spatial join using `geopandas.sjoin(predicate = ‘within’)` was performed to ultimately calculate the number of nuclei fully within each Visium spot (**Fig S7D**).

### Single-nucleus RNA-seq

#### snRNA-seq data collection and sequencing

Using 100μm cryosections collected from each donor, we conducted single-nucleus RNA-sequencing (snRNA-seq) using 10x Genomics Chromium Single Cell Gene Expression v3.1 technology. Approximately 70-100 mg of tissue was collected from each donor, placed in a pre-chilled 2 mL microcentrifuge tube (Eppendorf Protein LoBind Tube, Cat #22431102), and stored at −80°C until the time of experiment. To isolate nuclei, cryosections from each donor were combined with chilled Nuclei EZ Lysis Buffer (MilliporeSigma #NUC101) into a glass dounce. Sections were homogenized using 10-20 strokes with both loose and tight-fit pre-chilled pestles. Homogenates were filtered through 70 μm mesh strainers and centrifuged at 500g for 5 minutes at 4°C using a benchtop centrifuge. Nuclei pellets were resuspended in fresh EZ lysis buffer, centrifuged again, and resuspended in wash/resuspension buffer (1x PBS, 1% BSA, 0.2U/μL RNase Inhibitor). Final nuclei were washed in wash/resuspension buffer and centrifuged a total of 3 times. All nuclei were labeled with propidium iodide (PI) at 1:500 in wash/resuspension buffer and subsequently filtered through a 35 μm cell strainer. We performed fluorescent activated nuclear sorting (FANS) using a MoFlo Legacy Cell Sorter at the Johns Hopkins School of Public Health Cell Sorting Core Facility. Gating criteria were selected for whole, singlet nuclei (by forward/side scatter) and G0/G1 nuclei (by PI fluorescence). We sorted 14,000 nuclei based on PI+ fluorescence to include both neuronal and non-neuronal nuclei from each donor. This resulted in a final *n*=31 for snRNA-seq (1 PI+ sample for all 31 donors) with a total of 14,000 sorted nuclei per donor. We expected to retain approximately 2 out of 3 nuclei, resulting in a target of about 9,000 nuclei per donor. Samples were collected over multiple rounds, each containing 1-4 donors per round. All samples were sorted into reverse transcription reagents from the 10x Genomics Chromium Next GEM Single Cell 3’ v3.1 (dual-index) kit (without enzyme). Reverse transcription enzyme and water were added to bring the reaction to full volume. cDNA synthesis and subsequent library generation was performed according to the manufacturer’s instructions (CG000315, revision E, 10x Genomics). Samples were sequenced on an Illumina Novaseq 6000 at the Johns Hopkins University Single Cell and Transcriptomics Sequencing Core to a minimum depth of 50,000 reads per nucleus.

#### snRNA-seq data processing and droplet quality control

Sequenced chromium data was processed with *CellRanger* v7.2.0^112^, using human reference genome GRCh38 and the 2020-A annotation. Samples were sequenced to a median depth of 525,073,680 reads, with a median of 5,369 median unique molecular indices (UMIs) per nucleus, and a median of 2,452 median genes per nucleus. A total of 38,480,710 droplets (min: 581,462, max: 1,732,806) were sequenced.

Empty droplets were identified with *DropletUtils* v1.22.0 `emptyDrops(niters = 30000)`lower bound on UMI determined for each sample by the “knee point” calculated by *DropletUtils* `barcodeRanks()`plus 100, value ranging from 206 to 984 UMI^113^. A manual knee of 200 was set for 6 samples based on the UMI rank curve (**Table S5**). A total of 160,743 non-empty droplets were identified, with a median 5,222 nuclei per sample (min=2,169, max=7,847, **Fig S15A**).

Doublets were identified with *scDblFinder* v1.16.0 `scDblFinder()`^114^. A total of 10,820 doublets were detected and dropped (6.7% of non-empty droplets), leaving 149,923 single-nuclei droplets (**Fig S15A**). To further quality control the remaining nuclei, we evaluated three quality metrics: high percent mitochondrial reads, low total counts per nuclei (`sum_umi`), and low number of detected genes. An adaptive 3 median absolute deviation (MAD) cutoff for each sample was applied with `isOutlier()` from *scuttle* v1.12.0^109^, any nuclei that are outliers in one or more metrics were excluded (**Fig S16**). A total of 140,119 nuclei passed droplet quality control checks (93.4% of single-nuclei droplets), median 4,589 nuclei per donor; min=1,811, max = 6,946 (**Fig S15**).

#### snRNA-seq preliminary clustering and annotation

Reduced dimensions were calculated with generalized linear models principal component analysis (GLM-PCA)^115^. The top 2,000 deviant genes were selected with `devianceFeatureSelection()` from *scry* v1.14.0^116^. Principal components (PCs) were calculated with *scry* `nullResiduals()` and *scater* v1.30.1 `runPCA(ncomponents = 100)`^109^.

Batch correction was performed with *harmony* v1.2.1^24^, using `RunHarmony(group.by.vars = sample_id)` on the GLM PCs. Visual inspection of the *harmony-*corrected PCs showed a reduction of batch and donor effects (**Fig S17**).

The nuclei were then clustered using *scran* v1.30.2 `buildSNNGraph(k=10)`and *igraph* v2.1.4 `clusterwalktrap()`yielding 44 preliminary clusters^117,118^. Cell type annotations were automatically identified with *ScType*’s `sctype_score()`utilizing the provided brain marker gene database^119^ (**Fig S18**).

Clusters were assessed for quality by evaluating if the median value was over the cell class-specific (neuron or non-neuron) cutoff defined by the median + 3 * MAD cutoffs for sum UMI, detected genes, and percent mitochondria expression, or median `scDblFinder.score` > 0.5. The resulting cluster quality metric cutoffs were:

- Sum UMI: non-neuron > 1,340.94, neuron > 5,592.43
- Detected genes: non-neuron > 1,217.95, neuron > 4,174.27
- Percent mitochondria: non-neuron < 0.4, neuron < 0.53

Out of 44 clusters, 29 passed all quality metric cutoffs, retaining 125,771 nuclei (**Fig S19**).

#### snRNA-seq secondary optimal clustering and annotation

After dropping low quality clusters, normalized counts were recomputed with *scran* v1.34.0 quickCluster()and computeSumFactors(), then *scuttle* v1.34.0 logNormCounts()^117^ for computing log-normalized expression values (logcounts).

Reduced dimensions were recalculated using the GLM-PCA process described for the preliminary clusters^115^. Similarly, batch correction was repeated on the updated PCs using *harmony*^24^.

An optimal clustering was determined by testing values of *k* nearest neighbors from 10 to 30 in *scran* `buildSNNGraph()` and then clustered with *igraph* `clusterwalktrap()`^117,118^ (**Fig S20**). Clustering was evaluated by an elbow point in the number of clusters and the maximum mean silhouette score *bluster* v1.16.0 `approxSilhouette()` on *harmony* corrected PCs^120^. *k*=13 was selected as the optimal optimal clustering with max mean silhouette score 0.049 resulting in 23 clusters (**Fig S20**). Clusters were annotated by *ScType*^119^, and clusters were again evaluated for quality as described for the preliminary clusters, dropping one Oligo cluster, retaining 125,683 nuclei (**Fig S21**).

#### snRNA-seq non-neuronal subclustering and annotation

Nuclei identified as non-neuronal cell types (98,669 nuclei from Astro, Micro, Oligo, OPC, and Endo cell types) were subclustered to identify more fine grained subclusters. The nuclei from each cell type were subset, then clustered using scran `buildSNNGraph(k=10)`and `clusterwalktrap()`similarly to the preliminary clustering^117,118^. Again clusters failing the quality cutoffs were dropped, resulting in 94,990 non-neuronal and a total of 122,004 nuclei with a median of 3,949 nuclei per sample (min=1,517, max=6,288, **Fig S23**, **Fig S24**, **Fig S26**).

Marker gene expression expression confirmed the clusters’s broad cell type identities (**Fig S27**)^19^. Gene expression modeling was run on subclusters using `registration_wrapper()` from *spatialLIBD* v1.21.3^26,27^. Single-cell registration with *spatialLIBD*^26–28^ to external datasets Franjic et al. fine subcluster populations subset to ERC, and PsyENCODE DLPFC cell types^29,39^ supported both the broad cell type and fine subcluster annotations (**Fig S30**). *MeanRatio* marker genes were found for cell subclusters using `get_mean_ratio()` from *DeconvoBuddies* v0.99.10^40^ (**Fig S28**, **Table S6**).

#### Differential proportion analysis

Differential proportion analysis was performed for SpDs in Visium data, and fine subclusters in snRNA-seq data.

The proportions of each cluster were compared using the differential proportion analysis tools from *crumblr* v1.0.0^37^. Count frequencies were transformed to centered log ratio values (CLR) by *crumblr*. Variance partition was completed with `fitExtractVarPartModel()` from *variancePartition* v1.38.1^121^.

For Visium differential SpD proportions were tested with `dream(∼ APOE_carrier + Sex + Anc_Afr + Age + Visium_slide)`^122^ implemented in *variancePartition*^121^, fitting `coef = “APOE_carrierE4+”` (**Fig S14**, **Table S4**). Covariate effects (`Sex`, `Anc_Afr`, `Age`, `Visium_slide`) were removed using `cleaningY()` from *jaffelab* v0.99.34^123^ (**Fig S14**).

For snRNA-seq differential cell cluster proportions were tested with `dream(∼ APOE_carrier + Sex + Anc_Afr + Age + exp_round`)^121,122^, fitting `coef = “APOE_carrierE4+”` (**Fig S33**, **Table S8**). From the previous model formula, covariate effects were removed using `cleaningY()` from *jaffelab* v0.99.34^123^ protecting `APOE_carrier` (**Fig S33**), `APOE_carrier` and `Sex` (**Fig S34C-E**), or `APOE_carrier` and `Age` (**Fig S34H**).

#### Differential expression analysis

Differential expression (DE) on carrier status (E2+ vs. E4+) was run by “cluster” on pseudobulked data on Visium SpDs, and snRNA-seq broad cell types and fine resolution subclusters. Data was pseudobulked with *spatialLIBD* v1.21.5 `registration_pseudobulk(filter_expr = FALSE)`^26^, which has a `min_ncells = 10` default filter that that removes pseudobulked samples with less than 10 spots for SpDs and cells for snRNA-seq. Clusters with less than two pseudobulked samples for either carrier status were excluded from DE analyses.

By cluster, genes were normalized and filtered for low expression by *edgeR* v4.6.2 calcNormFactors() and filterByExpr.DGEList(), respectively^124,125^.

The *APOE* “carrier” model formula “∼0 + APOE_syn + Sex + Age + Anc_Afr + pseudo_expr_chrM_ratio” was fit with voomLmFit(block, adaptive.span = TRUE, sample.weights = TRUE) from *edgeR*^125^, using exp_round as a blocking variable in snRNA-seq and Visium_slide in Visium data. Contrast matrices were created and fit with makeContrast(), eBayes(), and topTable() from *limma* v3.64.1^126^ (**Figure 3**, **Fig S35**, **Figure 4**, **Fig S39**, **Fig S41**, **Figure 5C-D, Table S9**, **Table S11**, **Table S13**).

Ancestry-specific analysis were fit with model formula “∼0 + carrier_Anc + Sex + Age + pseudo_expr_chrM_ratio”, where carrier_Anc is a variable combining *APOE* carrier status and ancestry reported by legal next-of-kin and confirmed by death records (**Fig S37**).

Sex-specific analysis excluded genes from the X and Y chromosome, also did not consider ancestry as we had a low number of donors for females. Here we fit model formula “`∼0 + carrier_Sex + Sex + Age + pseudo_expr_chrM_ratio`”, where `carrier_Sex` is a variable combining *APOE* carrier status and sex (**Fig S38**).

#### Gene ontology overrepresentation analysis

Gene ontology overrepresentation analysis was run with `compareCluster()` *clusterProfiler* v4.16.0^127^. Dot plots were created with `dotplot()` (**Figure 3G**, **Fig S36A-B**, **Fig S40A-B**, **Fig S41**, **Figure 4D**, **Figure 5E**). Parent GO terms were found with `calculateSimMatrix()` and `reduceSimMatrix()` *rrvgo* v1.20.0^128^ (**Fig S36C-D**, **Fig S42**, **Table S10**, **Table S12**, **Table S14**).

#### Oligodendrocyte subcluster trajectory analysis

Oligo and OPC subcluster trajectory was predicted from PCs calculated on the snRNA-seq dataset, subset to just OPC and Oligo nuclei. Centroids were found with `aggregateAcrossCells()` from *scuttle* v1.18.0^109^. Minimum spanning tree was calculated with `createClusterMST()`, coordinates supplied with `reportEdges()`, and pseudotime calculated with `mapCellsToEdges()` from *TSCAN* v1.46.0 ^129^ (Figure 5A, Fig S29E).

#### DEG enrichment analysis with external datasets

DEG enrichment analyses were performed using `fisher.test(alt = “greater”)` for the 2 by 2 tables comparing a reference and a target gene set (**Fig S46**, **Fig S47**, **Table S15**), similar to how `gene_set_enrichment()` and `gene_set_enrichment_plot()` *spatialLIBD*^26^ are used for comparing enriched SpD marker gene sets against user-specified gene sets. However, this case was more complex as each comparison involved cluster-specific differential expression results.

### Integrative analyses

#### Spot deconvolution

The spot deconvolution method *RCTD*^41^ was used to integrate the Visium and snRNA-seq by identifying the proportions of cells in each Visium spot. For each sample we ran *RCTD* as implemented in *spacexr* v1.0.0 with `createRctd(UMI_min = 2)` and `runRctd(rctd_mode = “multi”, max_multi_types = 5)` in order to match the median number of nuclei intersecting with a spot (**Methods: Visium nuclei segmentation**), with either the broad or the fine resolution snRNA-seq data as the input. The broad resolution results confirmed the spatial registration results (**Figure 2**, **Table S7**). The fine resolution results were unreliable as only one subcluster per non-neuronal cell type was commonly detected, despite *RCTD* currently being frequently benchmarked as the most accurate spot deconvolution method^130,131^ (**Figure 2B**). This is likely due to the difficulty in detecting differences between highly similar subclusters (**Fig S29**).

#### Multicellular factor analysis

Multicellular factor analysis (MOFA) was performed with *MOFAcellulaR* v0.0.0.9^65^, as described in the vignette https://saezlab.github.io/MOFAcellulaR/articles/get-started.html#fitting-a-mofa-model. The pseudobulked Visium and snRNA-seq data was combined into a “mofa” object with `create_init_exp()`. A MOFA model was fit with `prepare_mofa()` and `run_mofa()` using default options except for `spikeslab_weights = FALSE` and `num_factors = 7`. Association with sample covariates was computed with `get_associations()`. The MOFA heatmap plot was created with `plot_MOFA_hmap()` (**Figure 6A, Table S17**). To assess significance of factor-demographic variable associations, `get_associations()` uses a linear model for continuous demographic variables and performs ANOVA for categorical variables. The associated p-value is Benjamini-Hochberg-corrected (FDR), with each factor constituting a test.

#### Cell-cell communication

To quantify ligand-receptor (LR) interactions by sample and cell type pair, *LIANA+* v1.5.1^59^ was first run using the snRNA-seq data as input. Specifically, `mt.rank_aggregate.by_sample()` was used with default arguments except `use_raw=True`, grouping by fine subcluster. To identify dataset-wide interactions of interest, ligand-receptor pairs for a given source and target subcluster were retained when their `magnitude_rank` was commonly observed, defined as significant (`magnitude_rank`< 0.05) in at least 20 of 30 samples (**Fig S43A-C, Table S16**). We performed a similar analysis at the broad cell type level, although no DEGs overlapped commonly observed LR interactions at broad cell type resolution.

To spatially validate ligand-receptor interactions, spatial connectivities were next computed in the Visium data for each sample using `ut.spatial_neighbors()`, with the exception of `set_diag=True` and using a sample-specific bandwidth precisely encompassing immediate-neighboring spots. Then, `mt.bivariate(global_name=’morans’, n_perms=1000, add_categories=True, use_raw=False)` was invoked for each sample. This produced a spatially aware score (bivariate score) at the spot level of the tendency of ligand and receptor to be locally co-expressed. The mean of this bivariate score was calculated for each LR interaction by SpD to assess domain-level co-expression (**Fig S43D, Table S16).**

### Software

Analyses were performed using R versions 4.3 to 4.5^132^ with Bioconductor versions 3.18 to 3.21^133^. Heatmaps were created with *ComplexHeatmap* v2.22.0-2.24.1^134^. Other visualizations were made using *ggplot2* v3.5.1 and v3.5.2^135^. Interactive web applications were made using *spatialLIBD* v1.21.5^26^ and *iSEE* v2.20.0^54^, and are both hosted on LIBD’s Posit Connect server with links listed at https://research.libd.org/LFF_spatial_ERC/.

## Supplementary Tables

**Table S1 Donor demographics.** Demographics of donors included in the study, including pathology information. Related to **Figure 1**, **Fig S1**.

**Table S2 SRT sample information**. Sample information for SRT dataset including data generation metrics. Related to **Figure 1**.

**Table S3 SpD marker genes.** SRT SpD marker gene statistics from enrichment and *MeanRatio* analysis. The statistics were computed with `get_mean_ratio()` from *DeconvoBuddies* and `registration_wrapper()` from *spatialLIBD*; see package documentation for the definition of the variable names. Related to **Figure 1**.

**Table S4 SRT SpD differential proportions.** Differential proportion statistics for SpDs from *crumblr* analysis computed with `treeTest()`. Related to **Fig S14**.

**Table S5 snRNA-seq sample information.** Sample information for snRNA-seq dataset including data generation metrics. Related to **Figure 2**.

**Table S6 Cell type subcluster marker genes.** snRNA-seq cell type marker genes from *MeanRatio* and enrichment analysis. The statistics were computed with `get_mean_ratio()`from *DeconvoBuddies* and `registration_wrapper()` from *spatialLIBD*; see package documentation for the definition of the variable names. Related to **Figure 2**, **Fig S28**.

**Table S7 Spot deconvolution results.** Spot deconvolution estimates from `runRctd()` by *RCTD* at the broad cell type resolution. Related to **Fig S29**.

**Table S8 snRNA-seq cell type differential proportions.** Differential proportion statistics for cell types from *crumblr* analysis computed with `treeTest()`. Related to **Fig S33**, **Fig S34**.

**Table S9 SRT differential expression results.** Differential expression statistics for SpDs for overall *APOE* carrier (E2+ vs. E4+), ancestry, and sex specific *APOE* carrier DGE analysis. The statistics were computed with `voomLmFit()` from *edgeR*; see its documentation for the definition of the variable names. Related to **Figure 3**.

**Table S10 SRT GO overrepresentation analysis.** SRT SpD GO results for overall *APOE* carrier (E2+ vs. E4+) DEGs, and ancestry and sex specific *APOE* carrier DGE analysis. The statistics were computed with `compareCluster()` from *clusterProfiler*; see its documentation for the definition of the variable names. Related to **Figure 3**.

**Table S11 snRNA-seq broad cell type differential expression results.** Differential expression statistics for broad cell types for overall *APOE* carrier (E2+ vs. E4+), ancestry, and sex specific *APOE* carrier DGE analysis. The statistics were computed with `voomLmFit()` from *edgeR*; see its documentation for the definition of the variable names. Related to **Figure 4**.

**Table S12 snRNA-seq broad cell type GO overrepresentation analysis.** snRNA-seq broad cell type GO results for overall DEGs *APOE* carrier (E2+ vs. E4+), and ancestry and sex specific *APOE* carrier DGE analysis. The statistics were computed with `compareCluster()` from *clusterProfiler*; see its documentation for the definition of the variable names. Related to **Figure 4**.

**Table S13 snRNA-seq subcluster differential expression results.** Differential expression statistics for fine subclusters for overall *APOE* carrier (E2+ vs. E4+), ancestry, and sex specific *APOE* carrier DGE analysis. The statistics were computed with `voomLmFit()` from *edgeR*; see its documentation for the definition of the variable names. Related to **Figure 5**.

**Table S14 snRNA-seq subcluster GO overrepresentation analysis.** snRNA-seq cell subcluster GO results for overall *APOE* carrier (E2+ vs. E4+) DEGs, and ancestry and sex specific *APOE* carrier DGE analysis. The statistics were computed with `compareCluster()` from *clusterProfiler*; see its documentation for the definition of the variable names. Related to **Figure 5**.

**Table S15 External dataset DEG enrichment results.** Gene set enrichment analysis statistics of DE gene sets against Grubman and Blanchard differential expression datasets^17,19^. These statistics were computed with `fisher.test()` from base R. Related to **Fig S46**, **Fig S47**.

**Table S16 Cell to cell communication results summary.** CCC summary data, including counts of commonly observed LR pairs by source and target, statistics for LR source-target interactions involving a DEG including mean bivariate score. See *Methods: Cell-cell communication* for summary methods. Related to **Fig S43**.

**Table S17 MOFA results.** MOFA statistics including gene expression variance explained (R^2^) for SpD and subcluster views, covariate p-values, and donor factor scores. The statistics were computed with `run_mofa()` from *MOFA2*; see its documentation for the definition of the variable names. Related to **Figure 6**.

