## Supplementary Figures for "*APOE* E4 Alzheimer’s Risk Converges on an Oligodendrocyte Subtype in the Human Entorhinal Cortex"

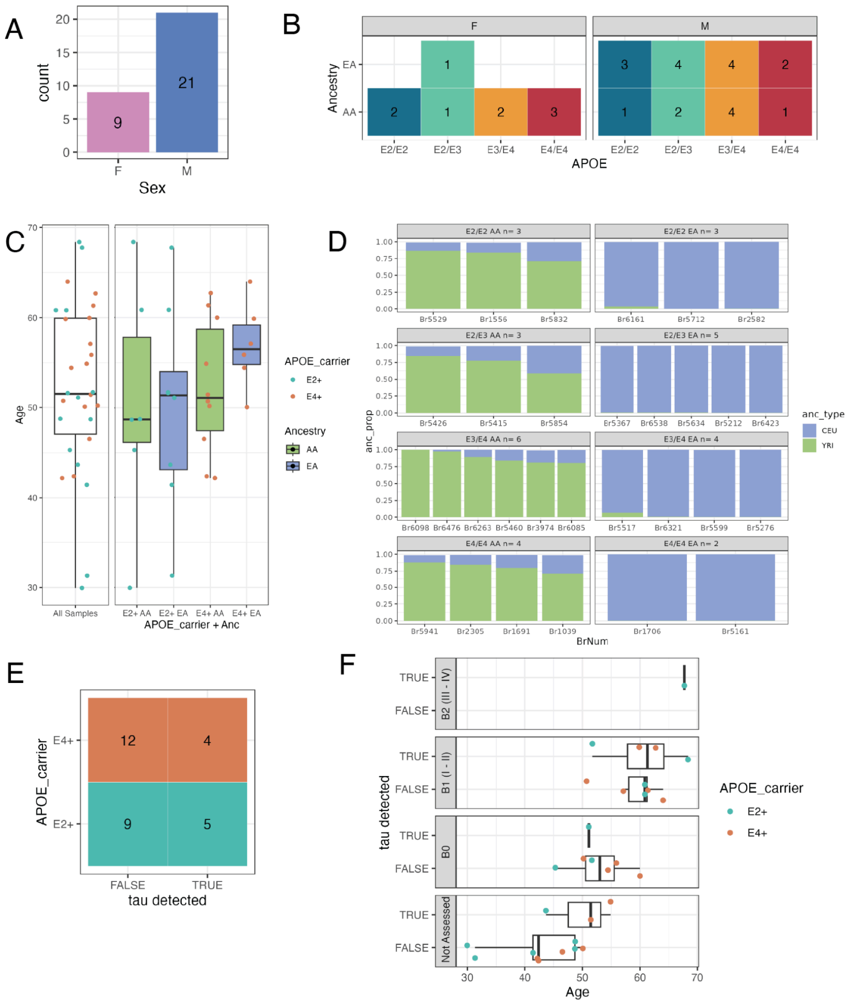


**Fig S1 Breakdown of sex, *APOE* genotype, age, and ancestry for 30 donors. A**. Barplot of donor sex. **B**. Tile plot of number of donors grouped by *APOE* genotype and ancestry. **C**. Boxplot of donor age, over all samples and divided by *APOE* genotype. **D**. Barplot of African (YRI) and European (CEU) ancestry fractions estimated from DNA genotype data for each donor (**Methods: Ancestry inference**), organized by *APOE* genotype and reported ancestry (African [AA] and European [EA]) . **E.** Tile plot of *APOE* carrier status and detected pTau in tissue samples measured by immunostaining. As expected, pTau signal was low, and thus quantified qualitatively as either present or absent. pTau was detected in 9 donors with more (5/14, 35.7%) pTau positive donors in E2+ carriers than in E4+ carriers (4/16, 25%). **F.** Boxplots of age by *APOE* genotype, pTau detection, and Braak stage (available for 18/30 donors, **Methods: Post-mortem human tissue samples**). Related to **Figure 1**, **Table S1**.

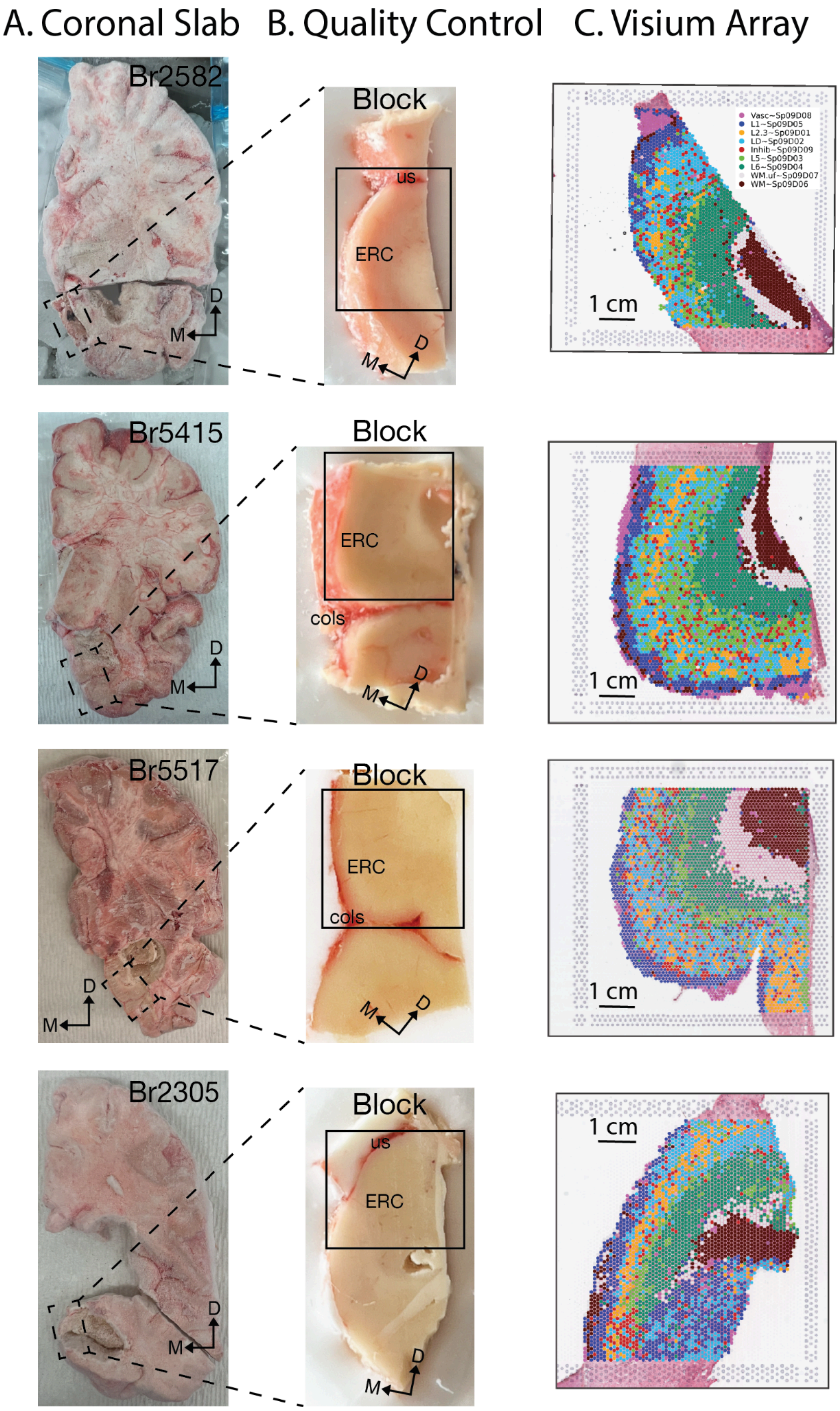


**Fig S2 Representative postmortem human ERC dissections and laminar organization. A**. Fresh-frozen coronal postmortem human brain slabs from control donors containing the ERC. Dashed boxes indicate the location of the dissected blocks in *B*. Neuroanatomical orientation is indicated by arrows: D, dorsal; M, medial. **B.** Dissected scored tissue blocks from donors in *A*. Black boxes indicate the approximate area placed on Visium capture areas in *C*. Neuroanatomical orientation is indicated by arrows: D, dorsal; M, medial. ERC - entorhinal cortex, cols - collateral sulcus, us - uncal sulcus. **C.** SRT tissue sections generated from brain blocks in *B*., showing *BayesSpace* clusters at *k*=9 taken from **Figure 1** and rotated to match the orientation in B. Related to **Figure 1**.

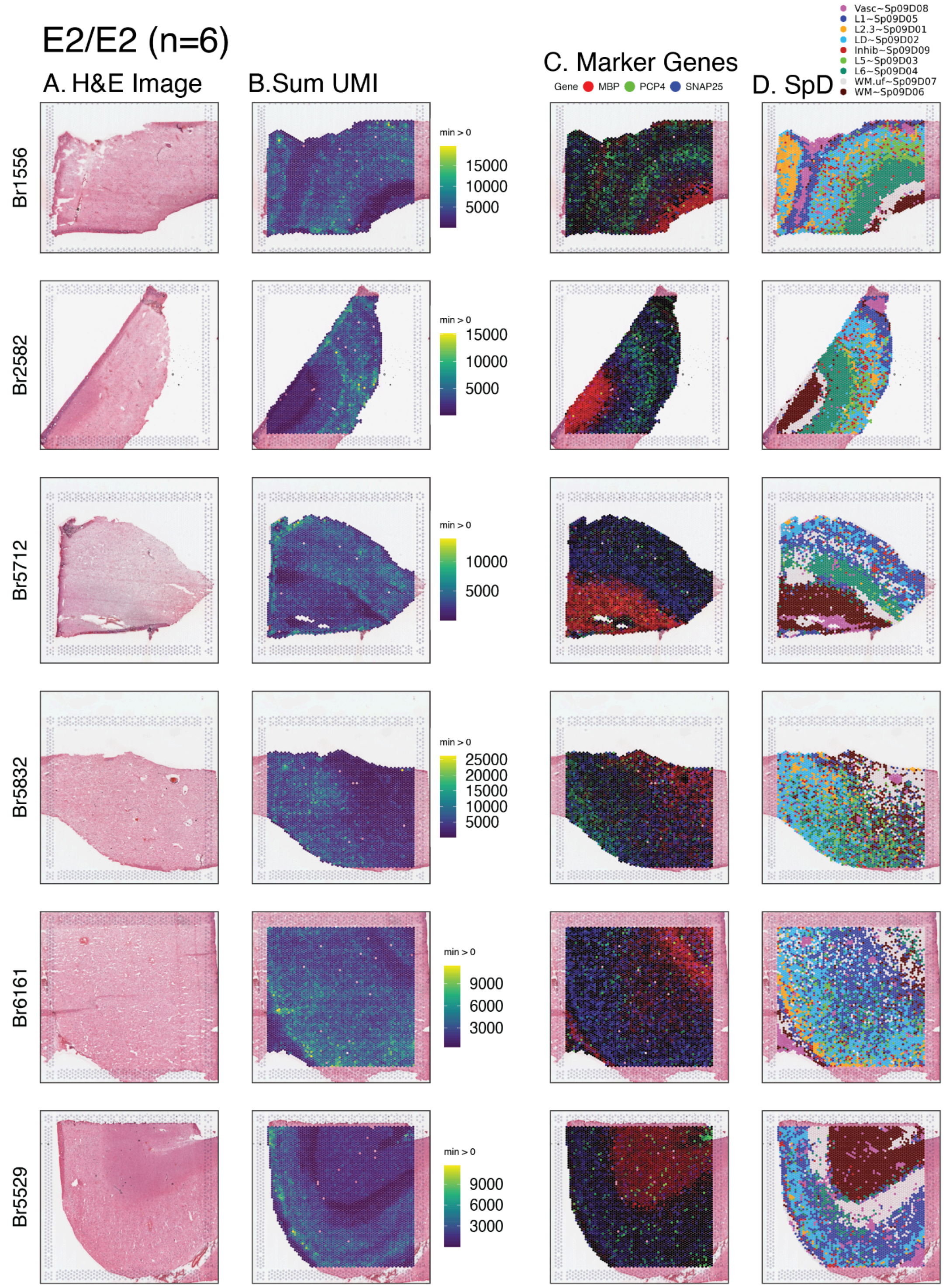


**Fig S3 SRT samples from E2/E2 donors. A.** Hematoxylin and Eosin (H&E) staining confirms orientation and morphology of tissue sections. **B**. Spotplot of sum unique molecular identifiers (UMI); **C.** Relative expression of spatial marker genes *MBP* (white matter), *PCP4* (Layer 5), and *SNAP25* (gray matter/neurons); and **D.** Spatial domains (SpDs, **Methods: Visium data processing and quality control**). Related to **Figure 1**, **Table S2**. **(cont.)**

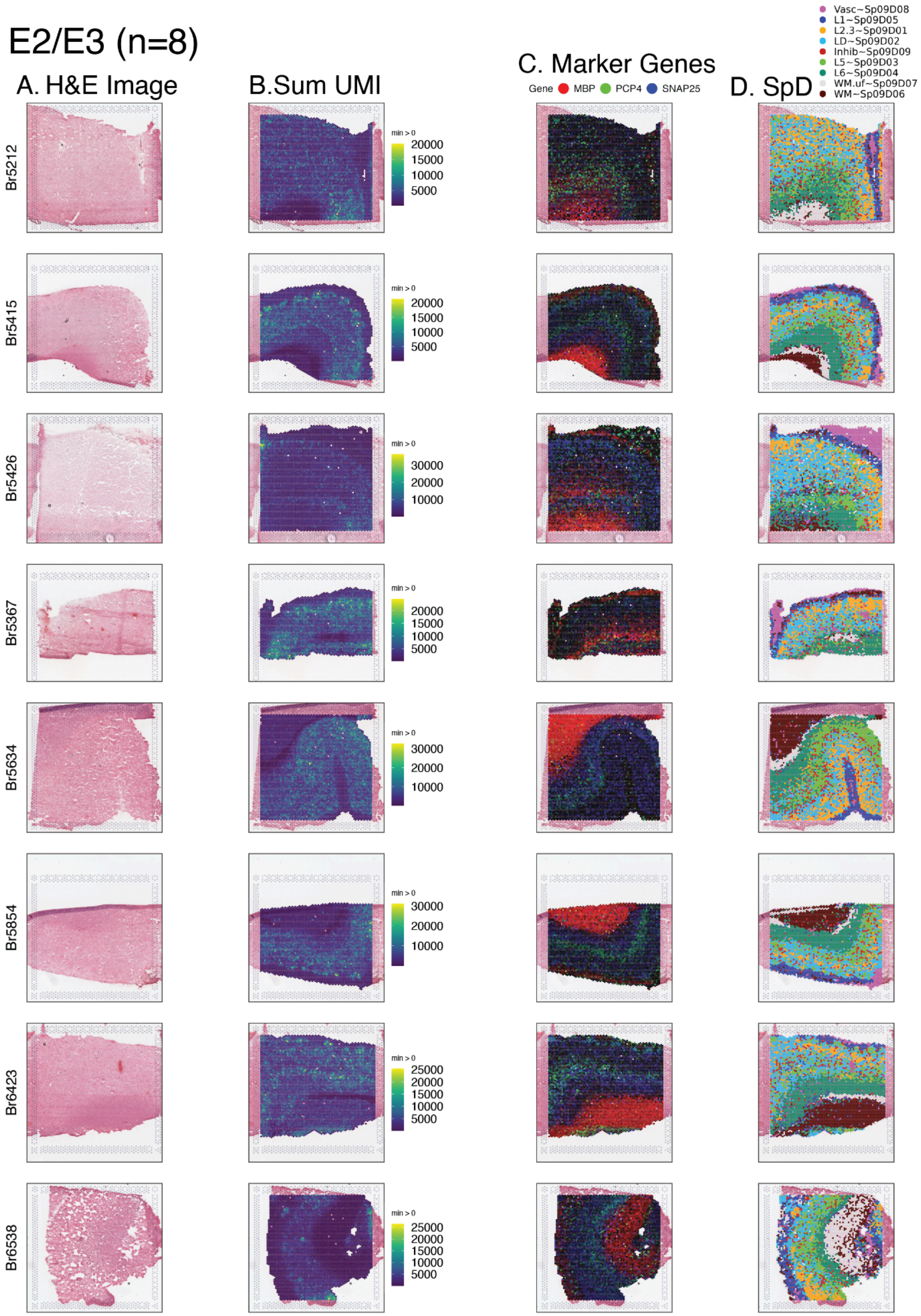


**Fig S3: SRT samples from E2/E3 donors. (cont.)**

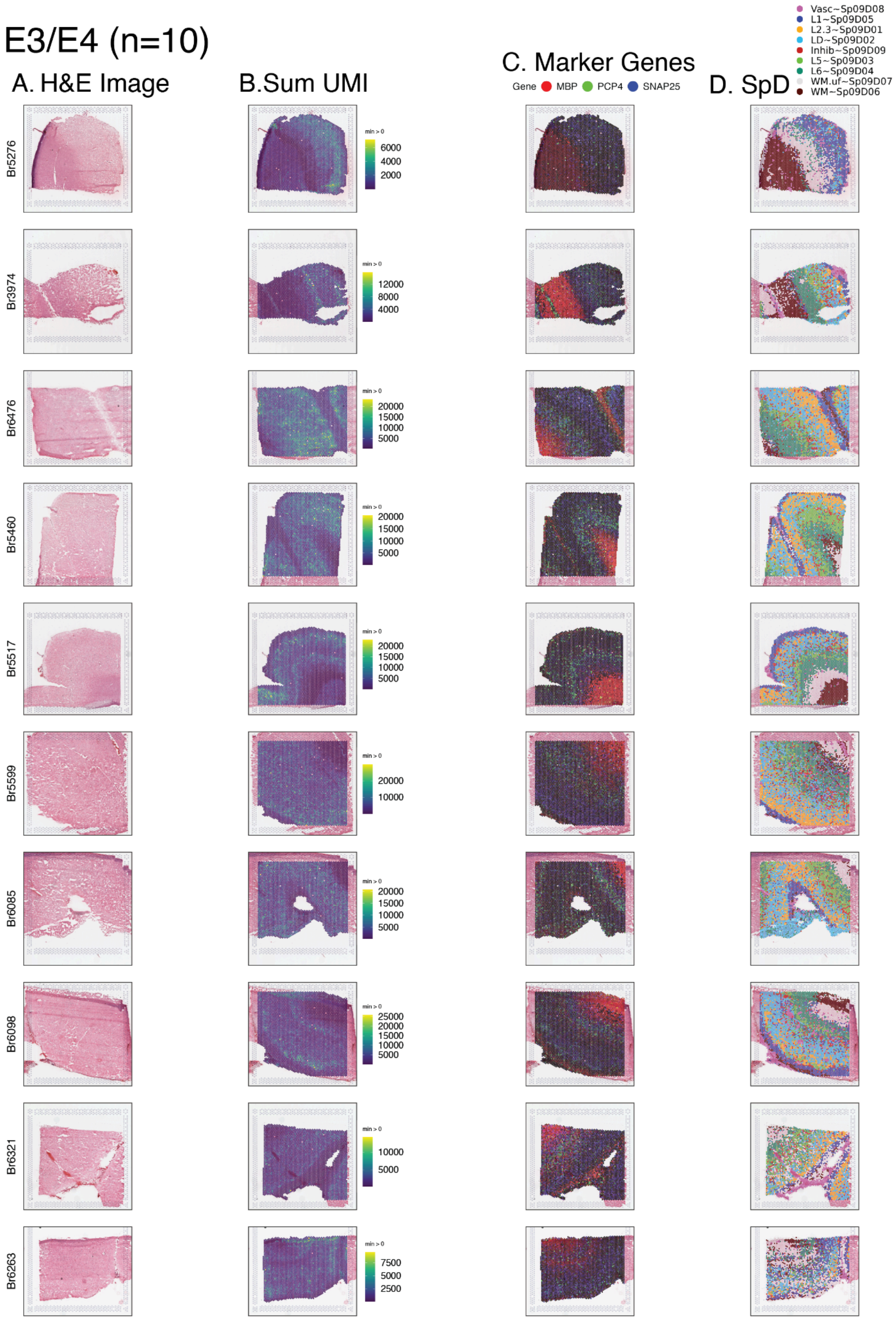


**Fig S3: SRT samples from E3/E4 donors. (cont.)**

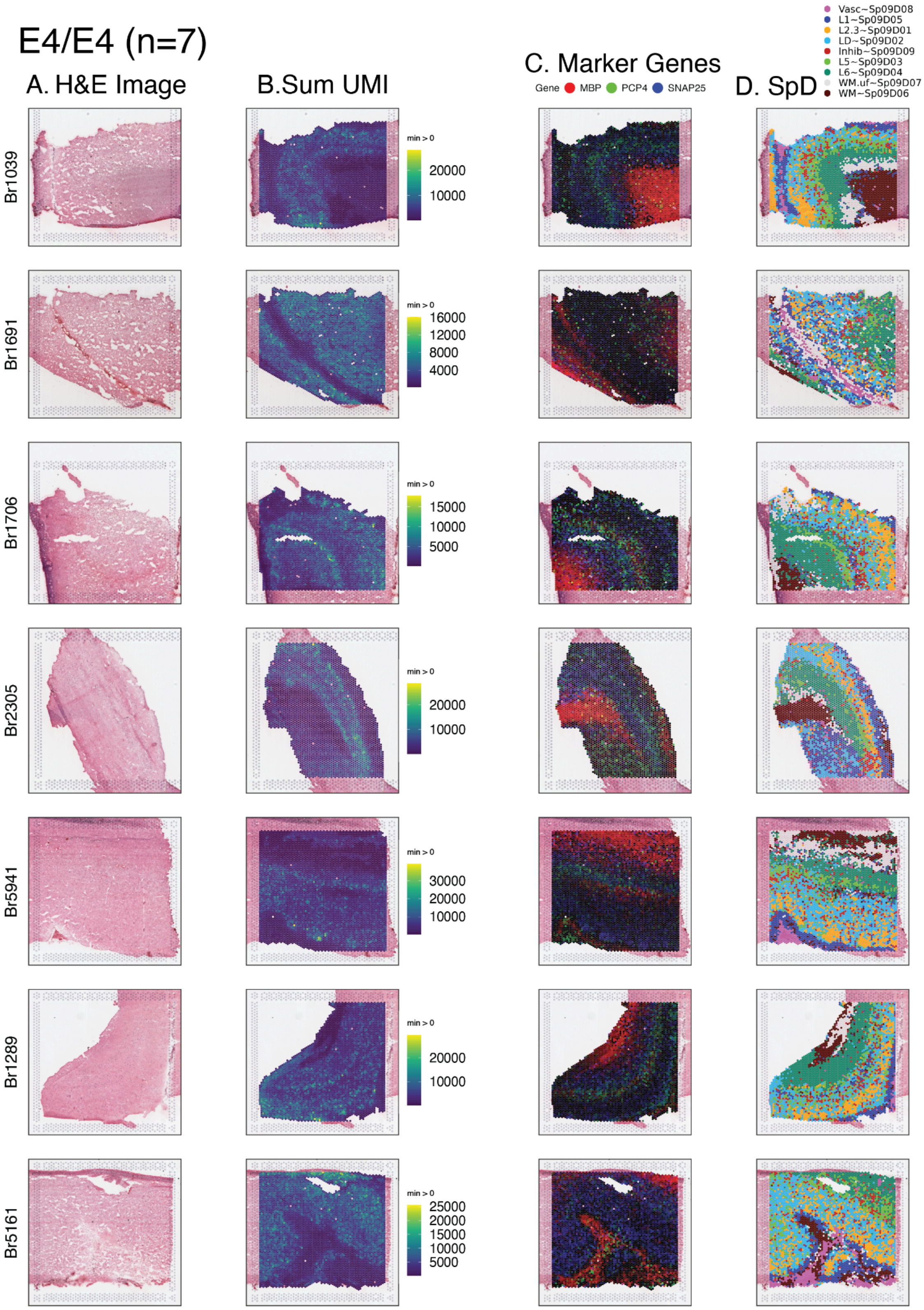


**Fig S3: SRT samples from E4/E4 donors. (cont.)**

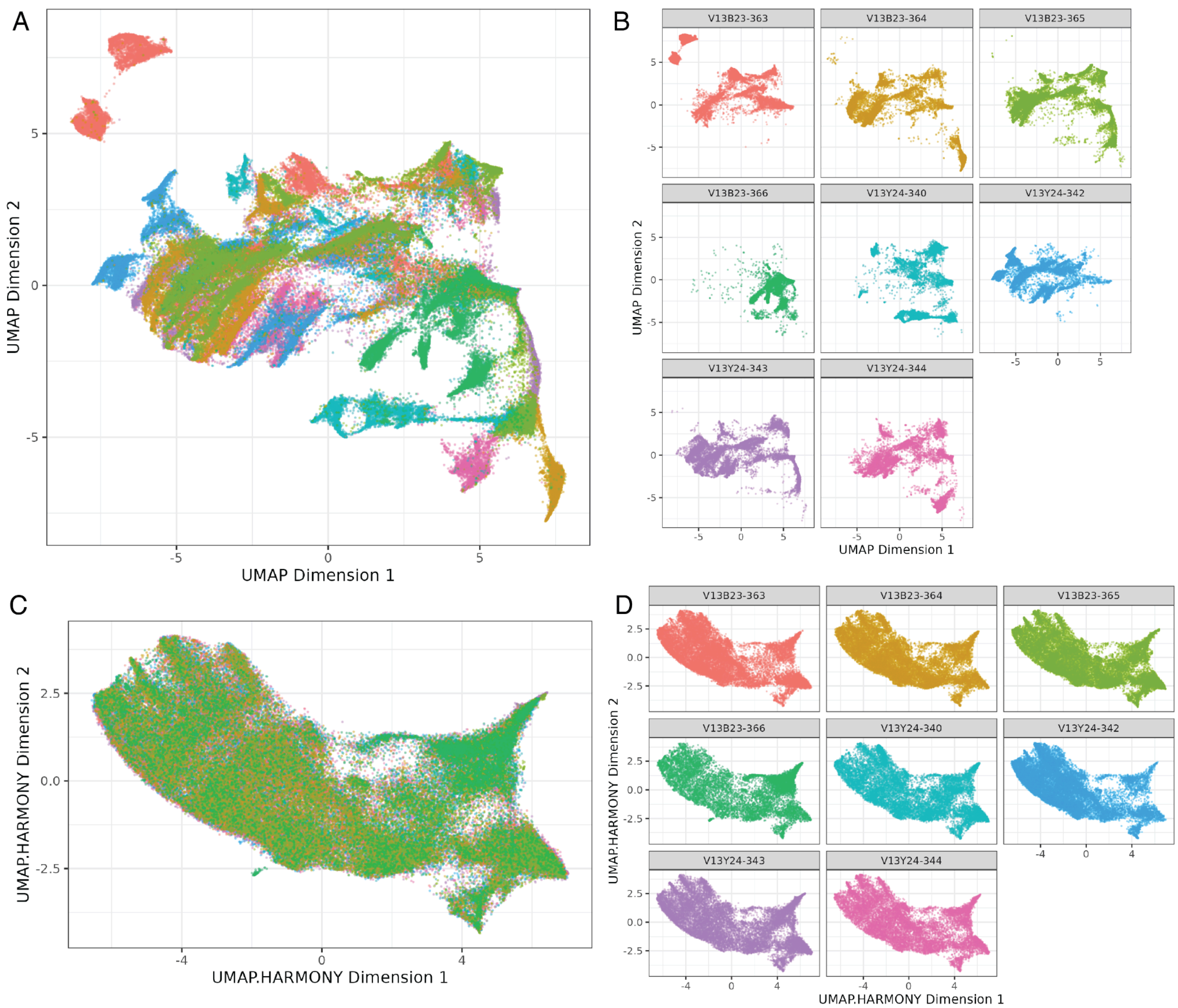


**Fig S4 UMAP of SRT data colored by Visium slide. A.** Pre-batch correction with *harmony* [24] **B.** faceted by Visium slide, and **C.** post-*harmony* batch correction by Visium capture area, **D.** faceted by Visium slide. Each Visium slide has four capture areas. See **Table S2** for details on which donors were captured in each Visium slide (**Methods: Visium spatial clustering**). Related to **Figure 1**.

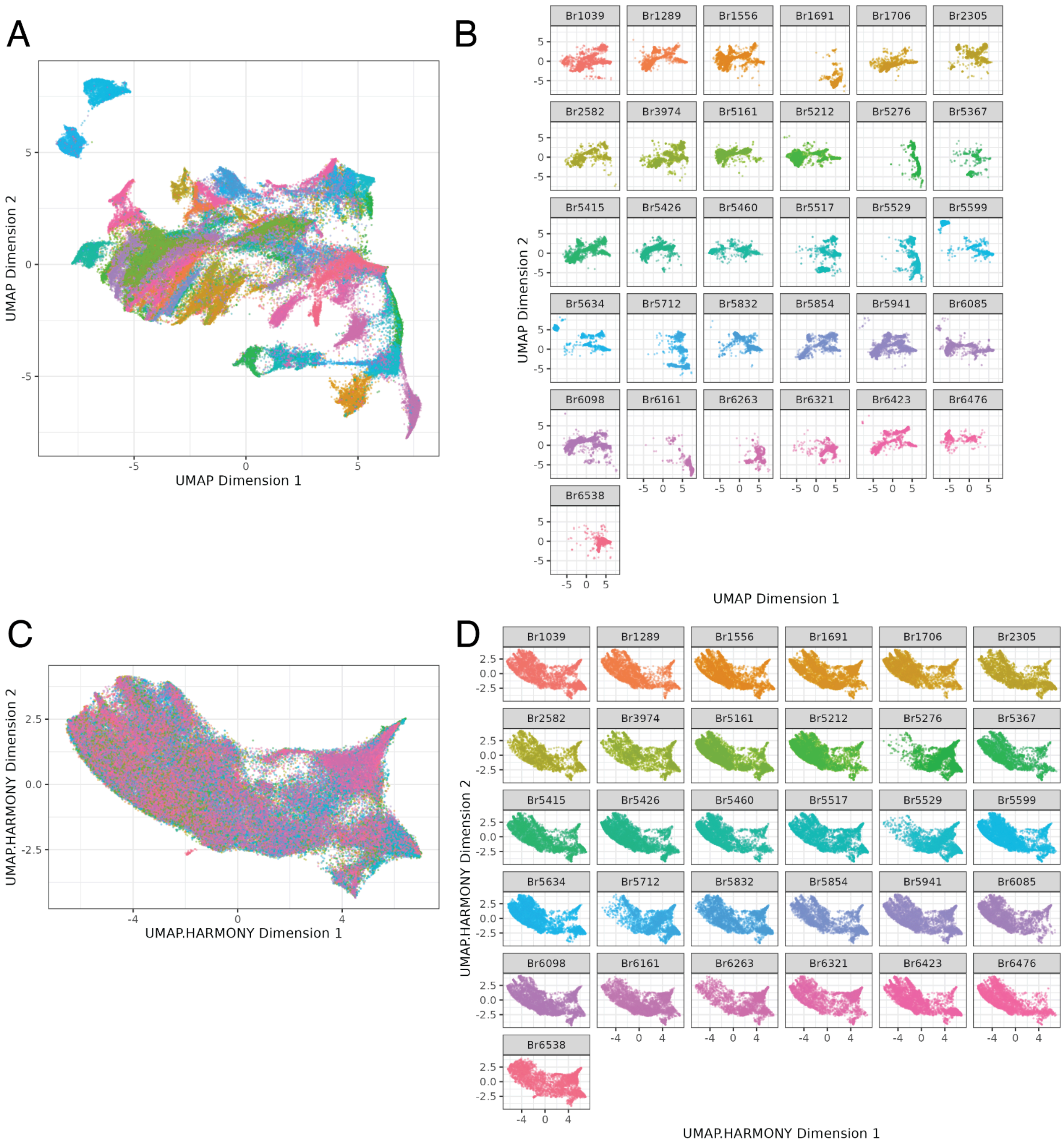


**Fig S5 UMAP of SRT data colored by donor. A.** Pre-batch correction **B.** faceted by donor, and **C.** post-*harmony* batch correction by Visium capture area, **D.** faceted by donor. Each donor was measured in a single Visium capture area (**Methods: Visium spatial clustering**). Related to **Figure 1**.

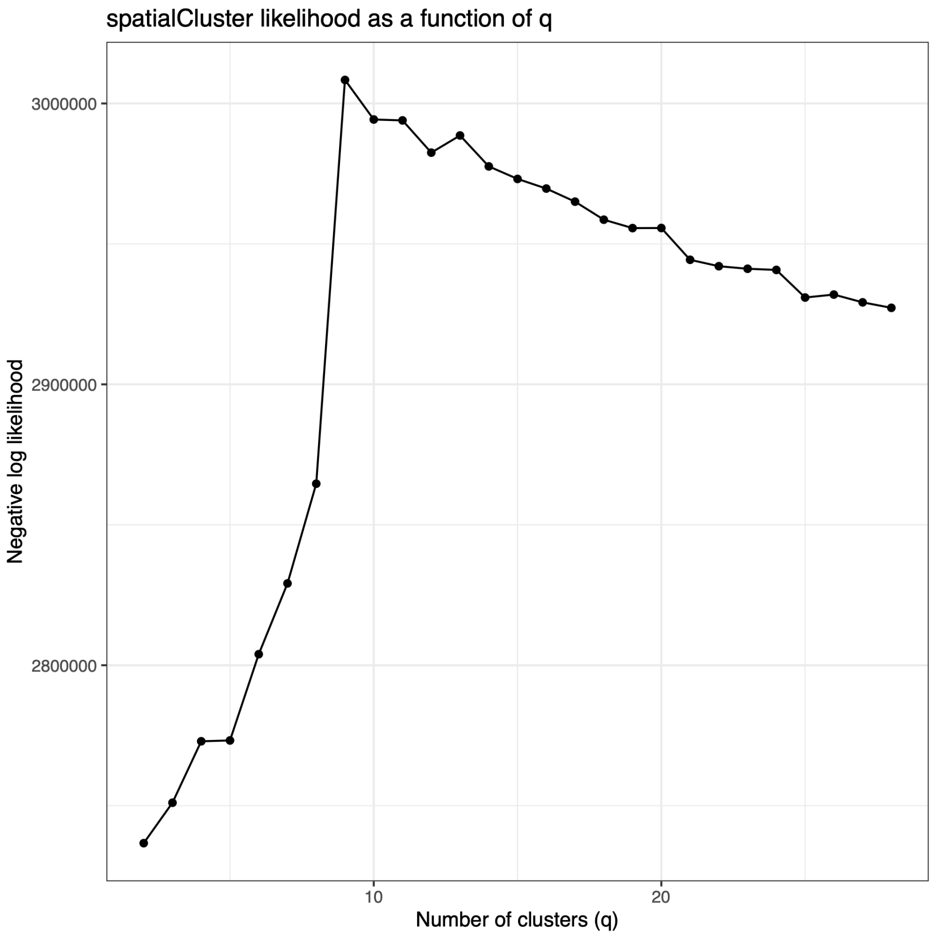


**Fig S6 Likelihood estimates predicted for number of *BayesSpace* clusters in Visium data.** Negative log likelihood estimates from `BayesSpace::qTune()` for *q* values 2 to 28 in increments of 1[25]. The optimal *q* for this data (*q* = 9) was selected by identifying the maximum negative log likelihood value (**Methods: Visium spatial clustering**). The number of clusters *q* is referenced as *k* elsewhere in this manuscript. Related to **Figure 1**.

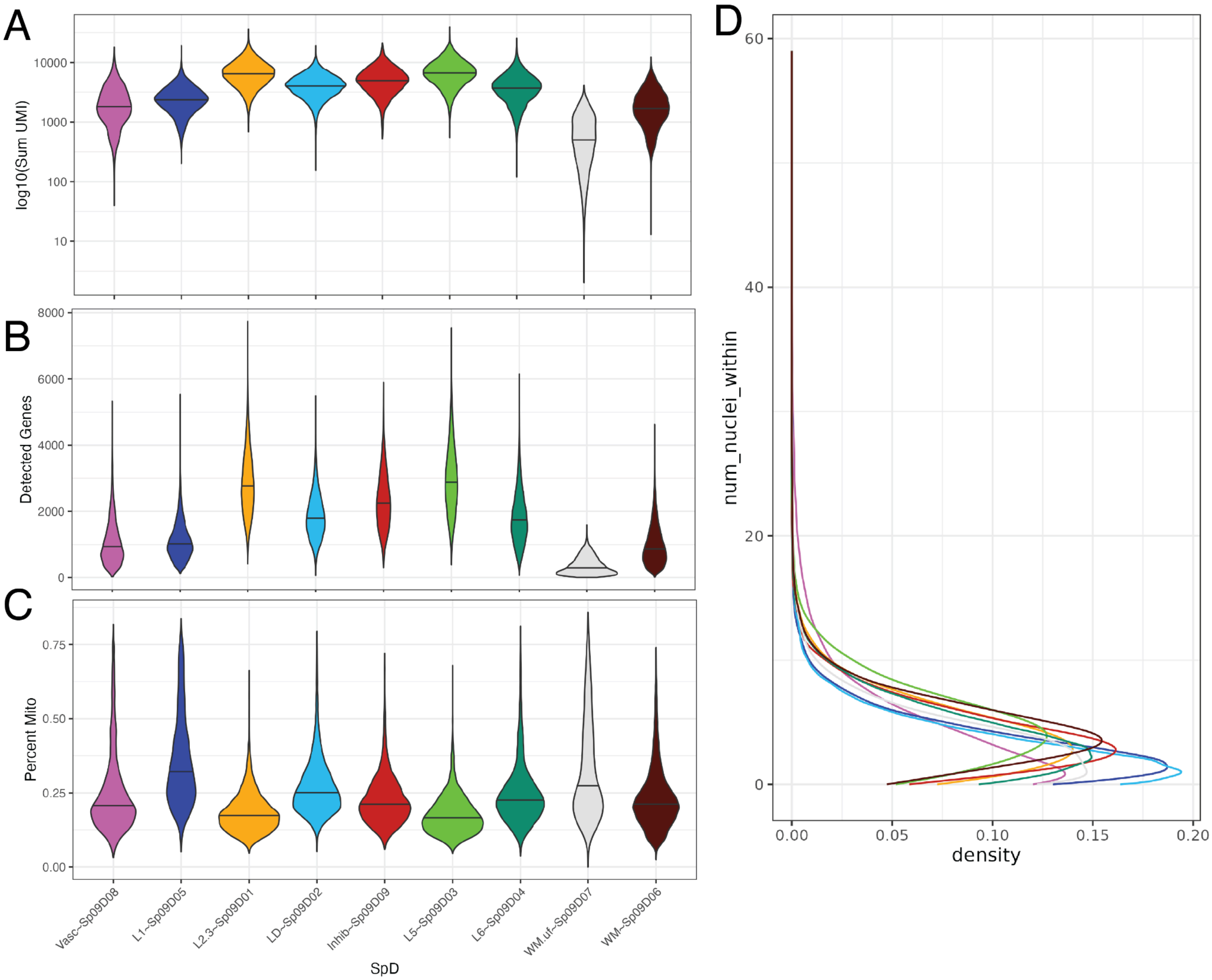


**Fig S7 Quality metrics over SRT spatial domains.** Violin plots of **A.** $\log_{10}$ of the sum of unique molecular identifiers (UMIs), **B.** number of detected genes, and **C.** mitochondrial expression for each spatial domain (SpD). **D.** Density of number of nuclei segmented within Visium spots by SpD (**Methods: Visium nuclei segmentation**). Related to **Figure 1**.

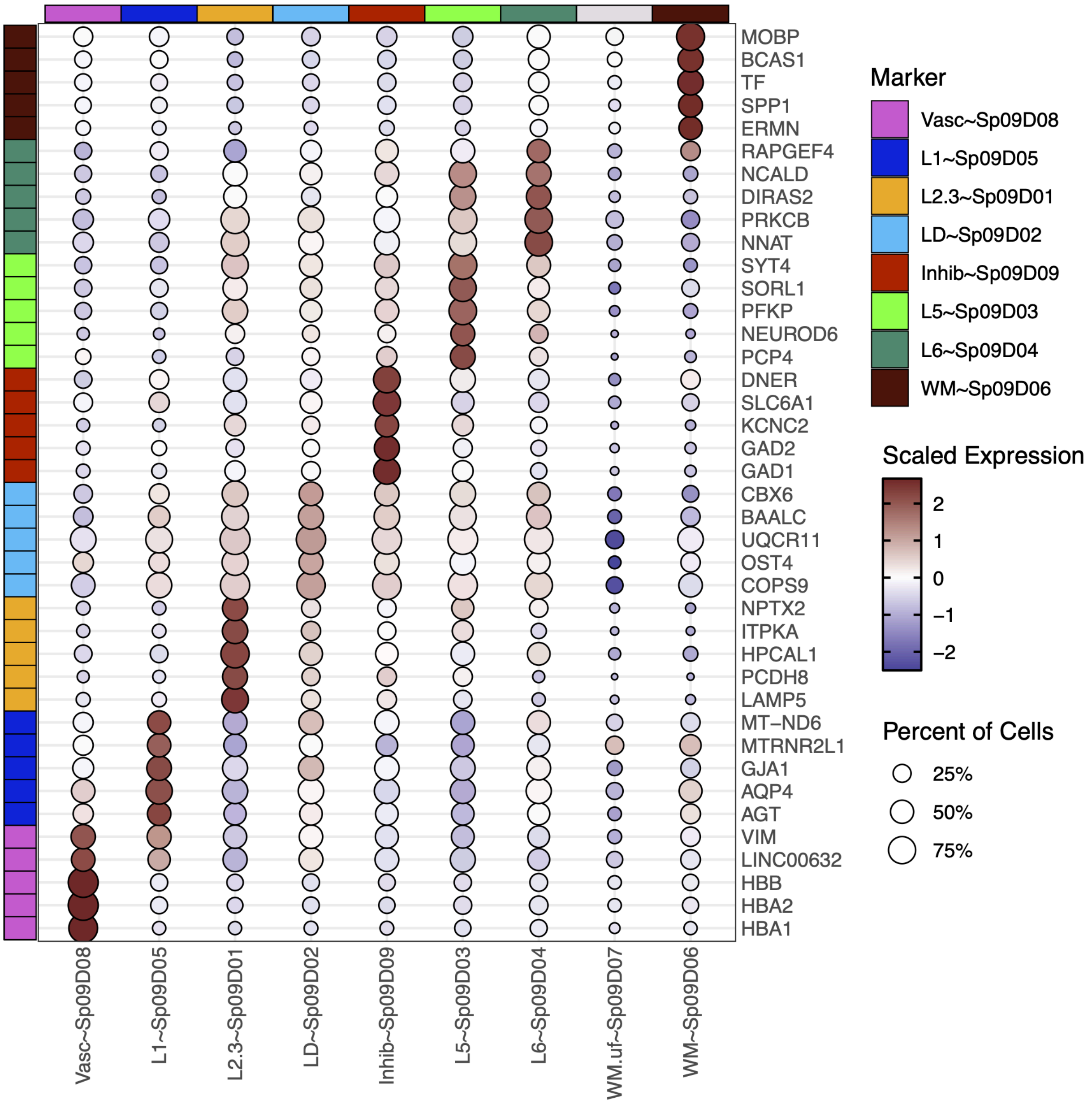


**Fig S8 Expression dotplot of *MeanRatio* marker genes for SRT Spatial Domains.** Rows pertain to the top five *MeanRatio* marker genes[40] for each spatial domain (SpD, columns). Log-normalized gene expression is scaled and centered. WM.uf~$Sp_9D_7$ showed low expression and did not have any *MeanRatio* marker genes with `MeanRatio > 1` (**Methods: Visium spatial domain annotation & analysis**). Related to **Figure 1**, **Table S3**.

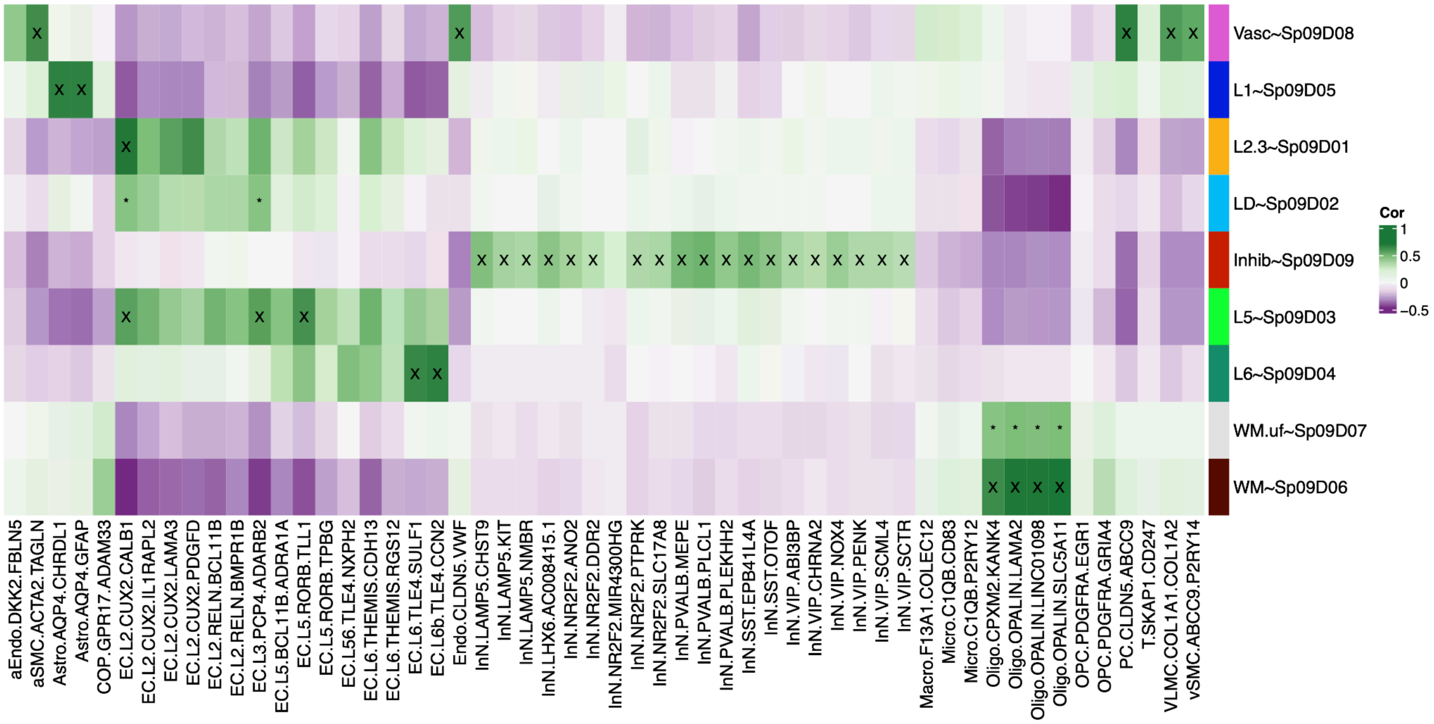


**Fig S9 Spatial domain registration to external human cortex datasets.** *spatialLIBD* spatial registration heatmap[26–28] of SpDs to the Franjic et al. ERC fine subcluster populations[29] (**Methods: Visium spatial domain annotation & analysis**). High confidence matches (cor > 0.5, merge ratio = 0.1) are marked with an "X" low confidence matches (max correlation but <0.5) are marked with "*". Related to **Figure 1**.

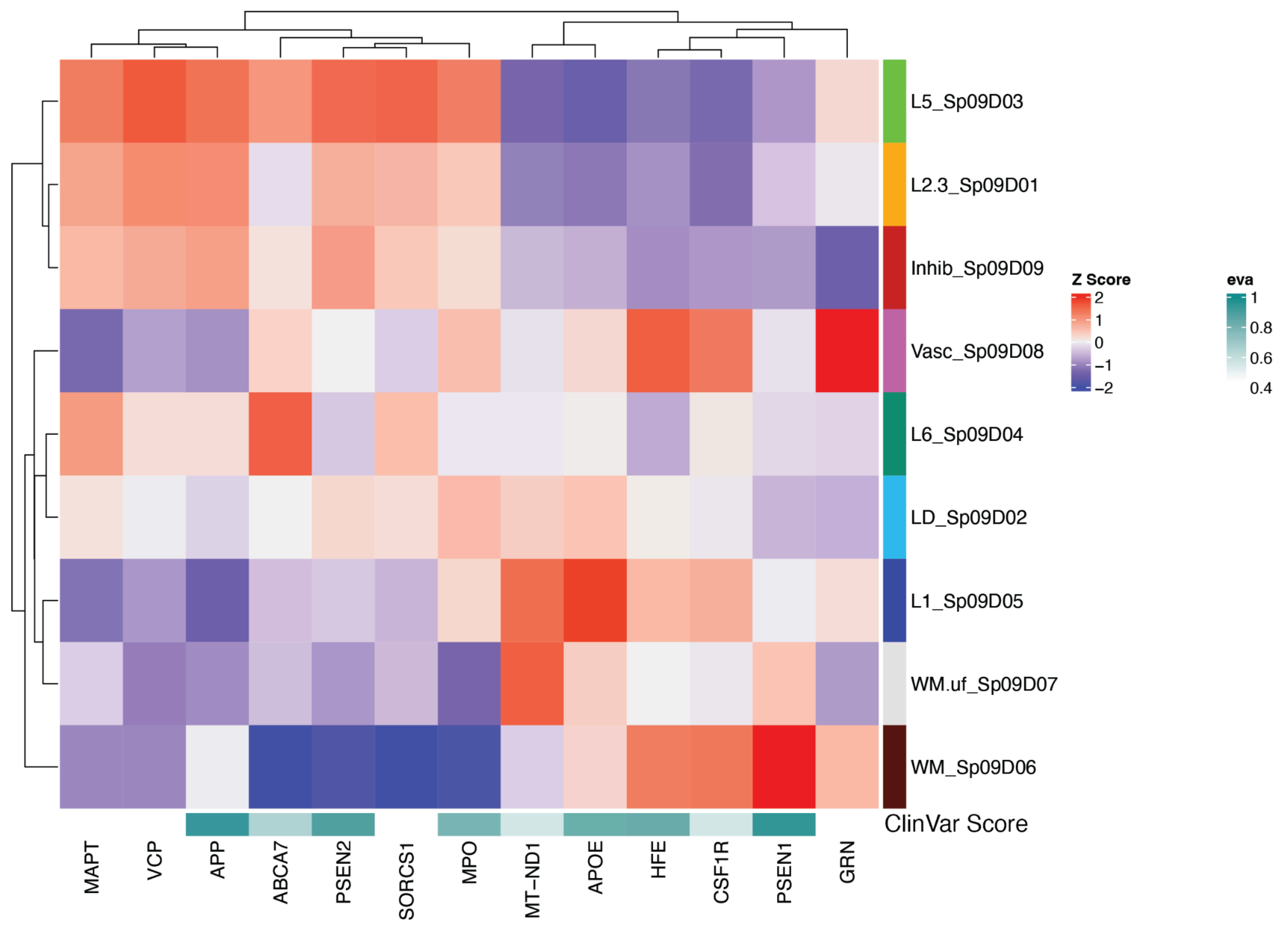


**Fig S10 Expression of AD risk genes across spatial domains.** Heatmaps of scaled and centered mean pseudobulked logcounts expression of *ClinVar* AD risk genes across SRT SpDs. *ClinVar* genes sourced from *OpenTargets*, where `eva` stands for European Variation Archive and in this case is the association score for SpD from EVA (available in *OpenTargets* through *ClinVar*)[110] (**Methods: AD risk gene set**). Related to **Figure 1.**

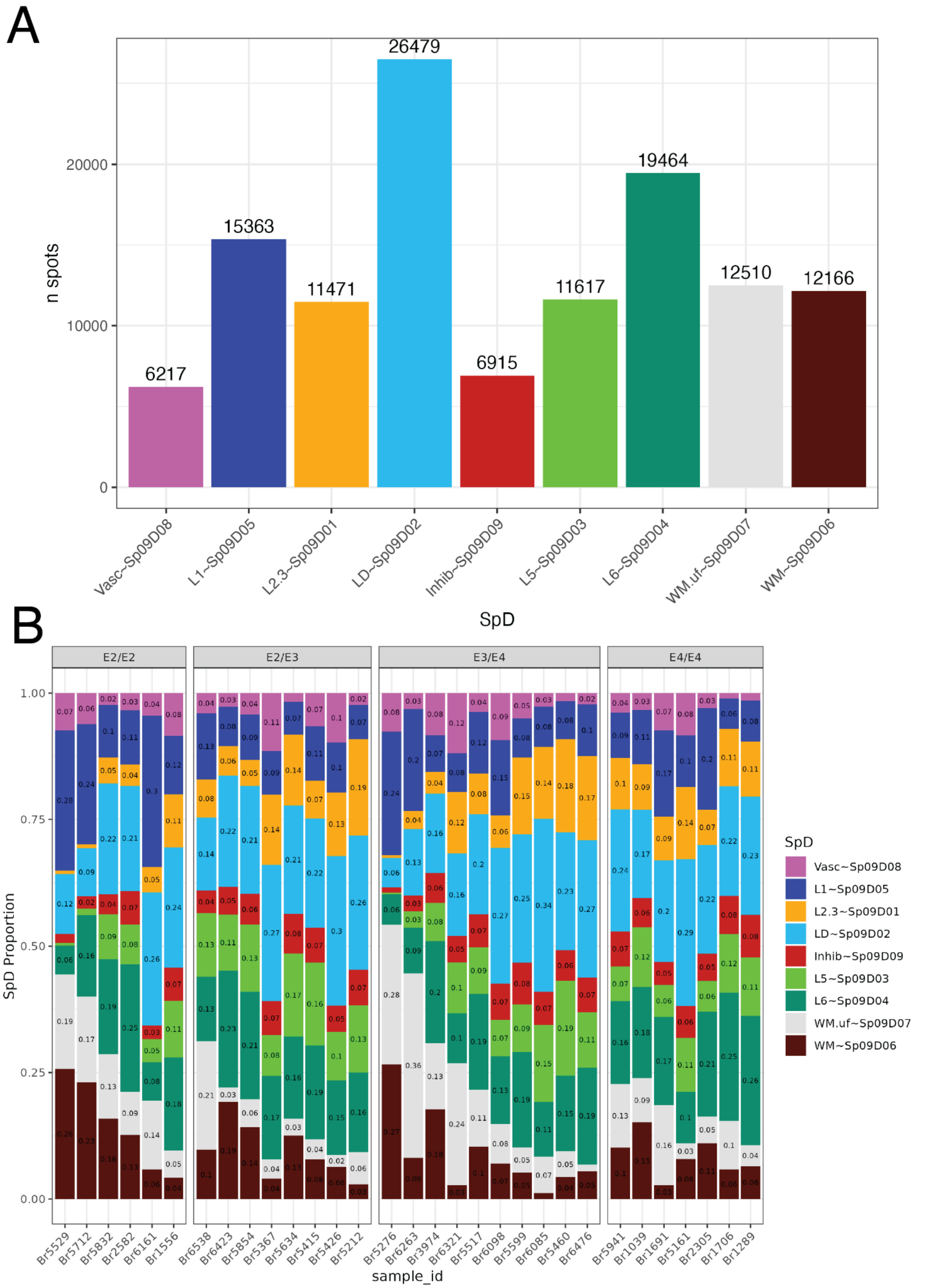


**Fig S11 Spatial domain composition across SRT samples. A.** Bar plot of the number of spots assigned to each spatial domain (SpD). **B.** Composition bar plots of the proportions of SpD for each donor, arranged from most to least white matter and grouped by *APOE* genotype (**Methods: Visium spatial clustering**). Related to **Figure 1**.

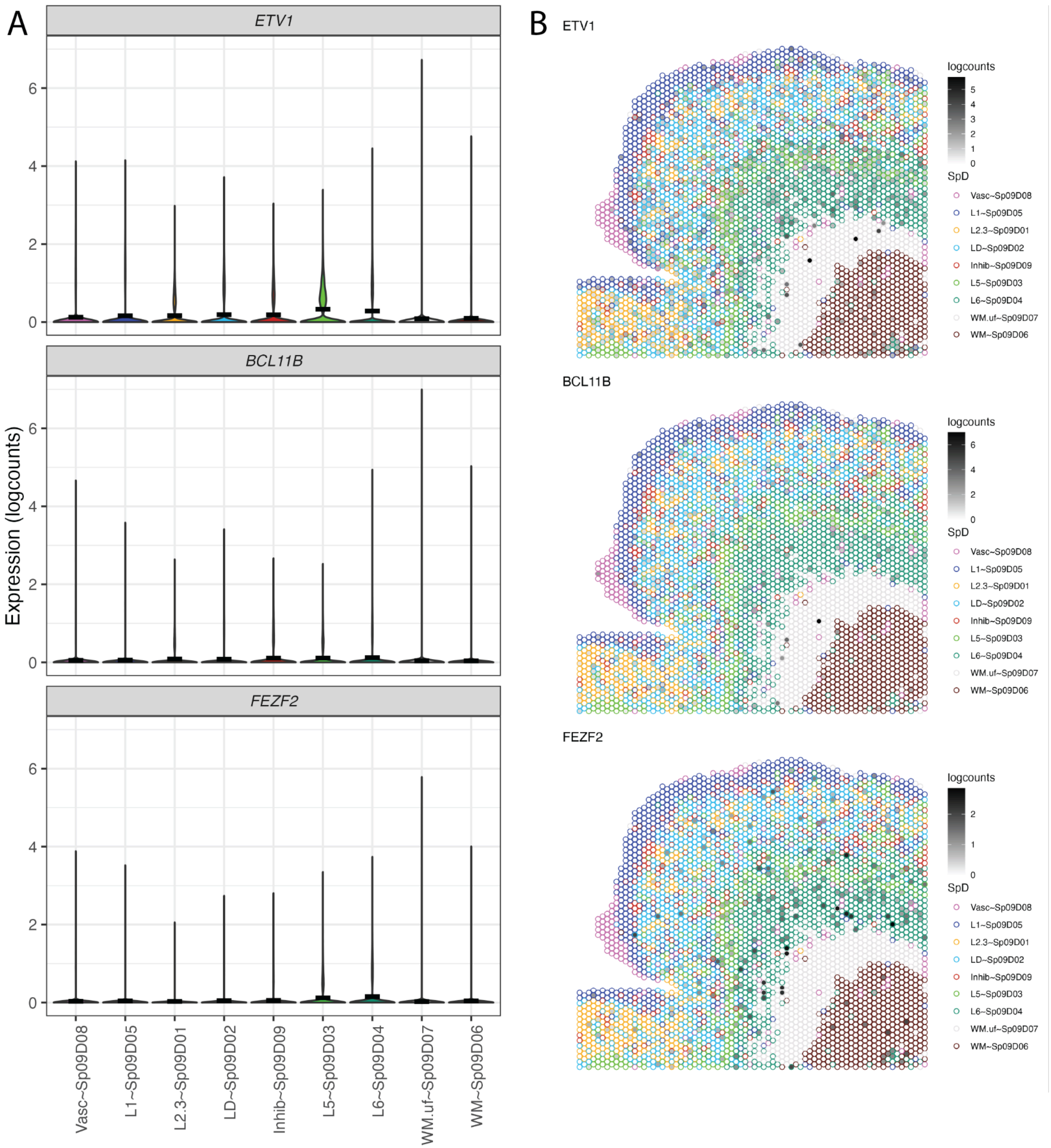


**Fig S12 Characterization of ERC layer 5.** Layer 5 sublayers had been previously reported in mouse ERC, with diverging expression of *ETV1* in L5a and *BCL11B* (Ctip2*)* and *FEZF2* in L5b[32,33]. **A.** Violin plots of gene expression for L5 sublayer markers. **B.** *escheR*[137] plots of representative sample Br5517, with gene `logcounts` as point fill layered over SpD identities (point outlines in different colors). Related to **Figure 1**.

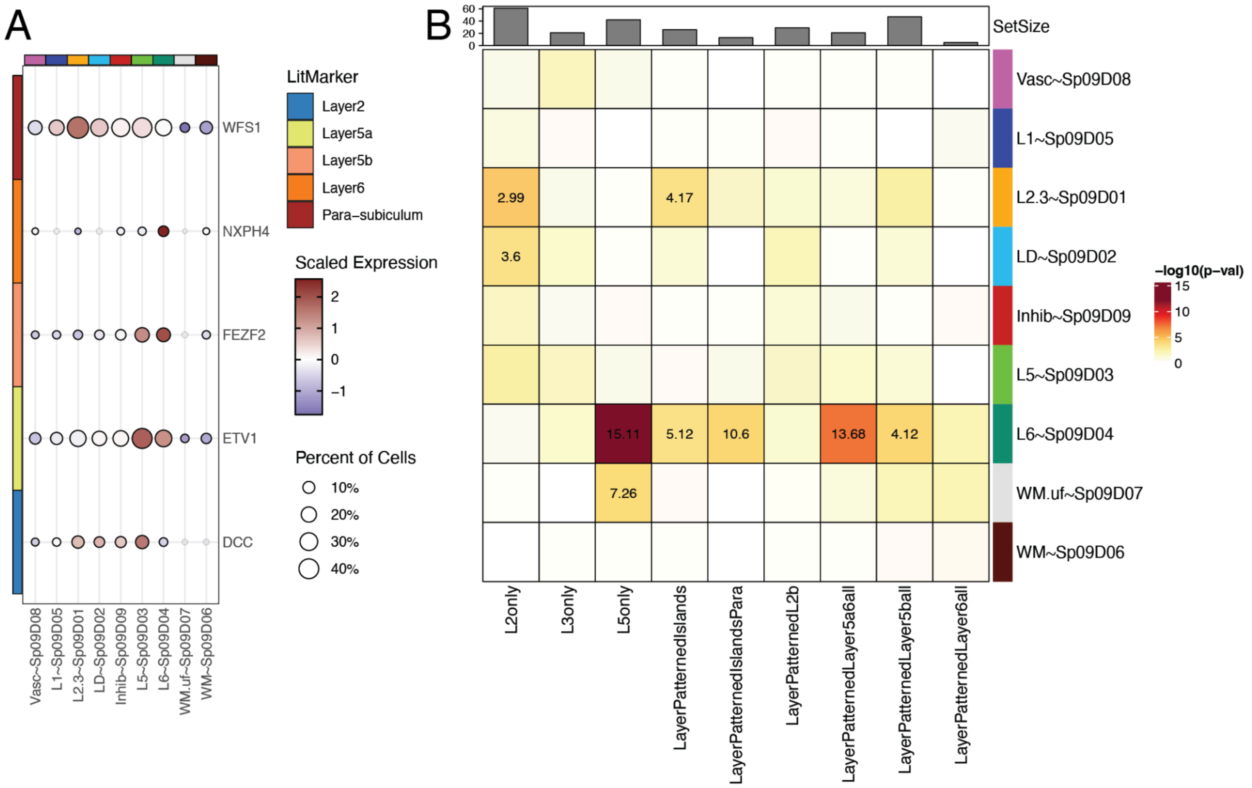


**Fig S13 Characterization of mouse ERC layer genes in human ERC spatial domains. A.** Dot plot of mouse ERC layer marker genes from Ramsden et al.[33]. **B.** Gene set enrichment heatmap of p-values for Fisher's exact test of SpD enrichment genes (FDR < 0.05) vs. mouse ERC layer marker genes from Ramsden et al.[33]. Values are odds ratios for statistically significant enrichments (p-val < 0.001). L2only genes were enriched in L2.3~$Sp_9D_1$ and LD~$Sp_9D_2$. L3only genes were not enriched in any SpDs. L5only genes were enriched in L6~$Sp_9D_4$ and WM.uf~$Sp_9D_7$. LayerPatternIslands gene sets were enriched in L2.3~$Sp_9D_1$ and L6~$Sp_9D_4$. Related to **Figure 1**.

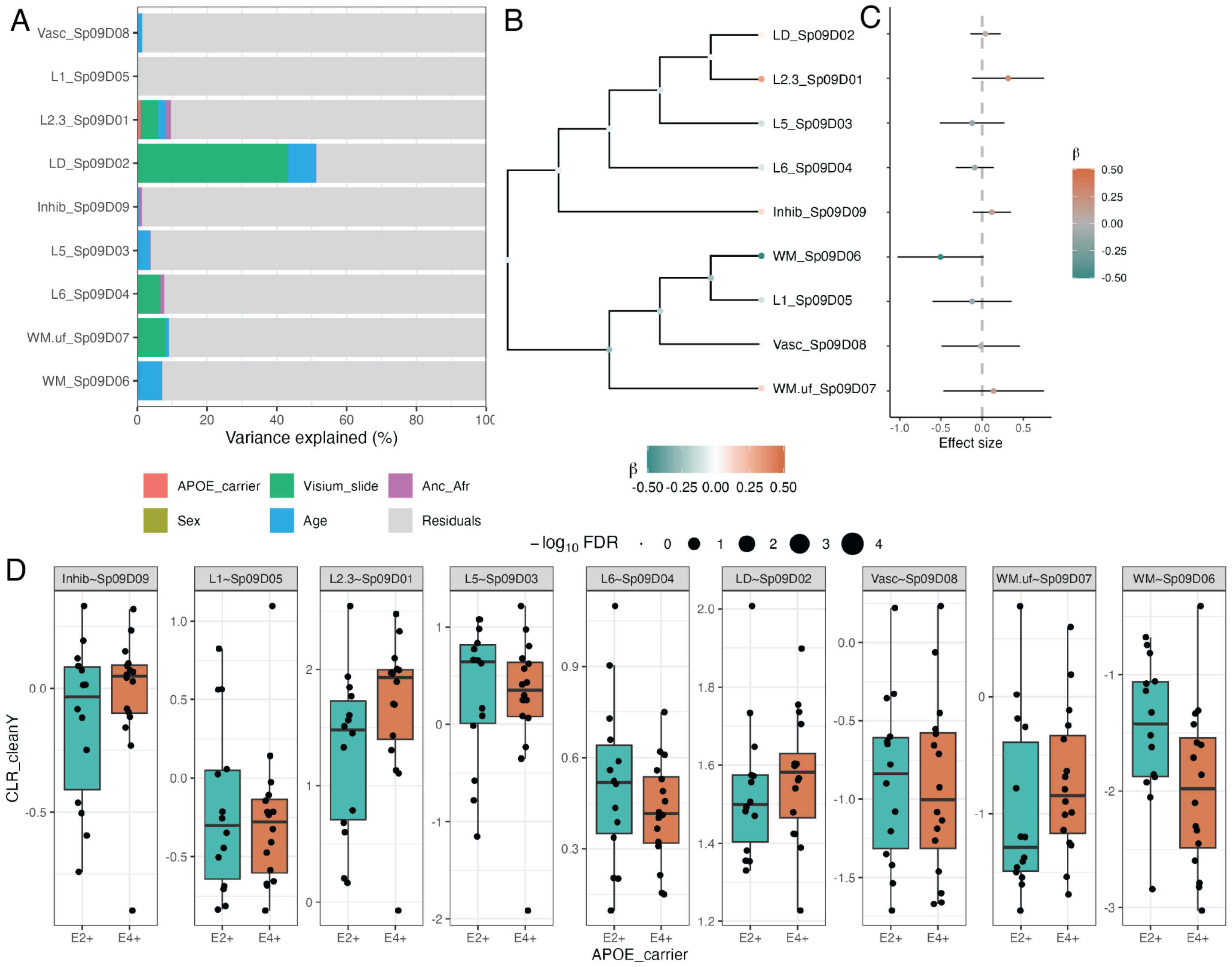


**Fig S14 Differential proportion analysis of spatial domains. A.** Barplot showing the percent of proportion variance explained by different variables calculated using *crumblr*[37]. **B.** Hierarchical tree of SpDs, node color indicates the estimated effect size from regression analysis of *APOE* carrier status (E2+ vs. E4+) at leaves (SpDs) and multivariate testing at internal nodes. Node point size represents FDR for each test. **C.** Forest plot of effect size from *APOE* carrier test. Effect size > 0 represents higher frequency of the cell type in E4+ donors, while effect size < 0 represents higher frequency in E2+ donors. The color corresponds to the effect size (β) and bars indicate the 95% confidence interval. **D.** Box plots of the centered log ratio (CLR) of SpD frequency over *APOE* carrier status, confounding variables regressed out with `cleaningY`, protecting `APOE_carrier`. (**Methods: Differential proportion analysis**). Related to **Figure 1**, **Table S4**.

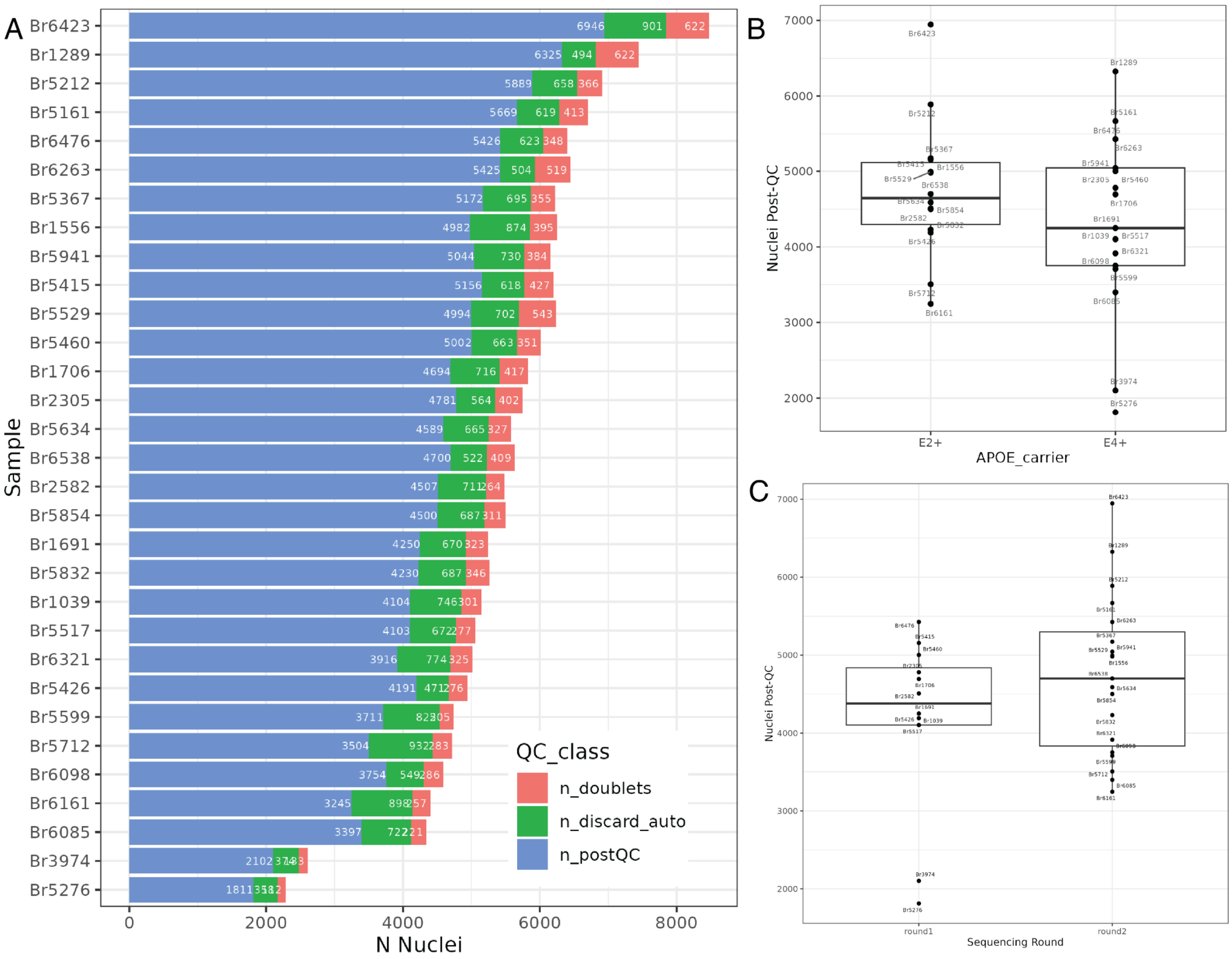


**Fig S15 Number of nuclei identified with snRNA-seq by donor and groups across quality control metrics. A**. Bar plot of number of nuclei per donor identified as doublets (red), or automatically discarded (`discard_auto`, green) for one or more poor quality metric (high percent mitochondrial reads, low sum umi, and low number of detected features), or passed quality control (post QC, blue). In total 140,199 nuclei passed droplet quality control. **B**. Boxplot of the number of post QC nuclei by *APOE* carrier status, with no significant difference across *APOE* carrier groups. **C**. Boxplot of the number of post QC nuclei by sequencing round, showing that data generation was overall consistent across the two rounds (**Methods: snRNA-seq data processing and droplet quality control**). Related to **Figure 2**, **Table S5**.

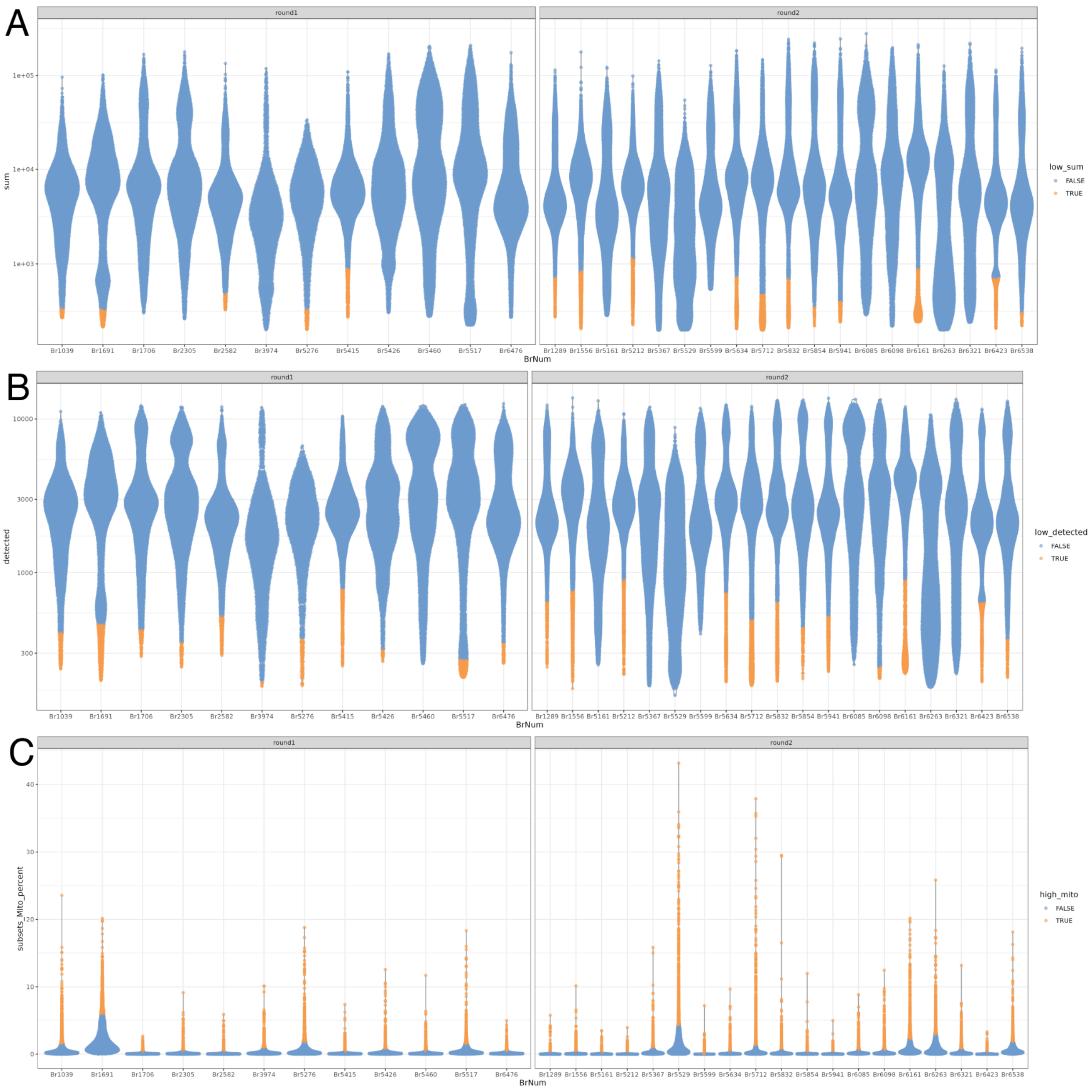


**Fig S16 snRNA-seq quality control metrics by donor.** Donor-specific cutoffs were selected with 3 median absolute deviations (MAD) away from the median using `scuttle::isOutlier()` applied at the cell level[109]. **A**. Total number of unique molecular identifiers (`sum_umi`) in a $\log_{10}$ scale. **B**. Number of genes detected in a cell (`detected`). **C**. Percent of mitochondrial gene expression (`subsets_Mito_percent`, **Methods: snRNA-seq data processing and droplet quality control**). Related to **Figure 2**.

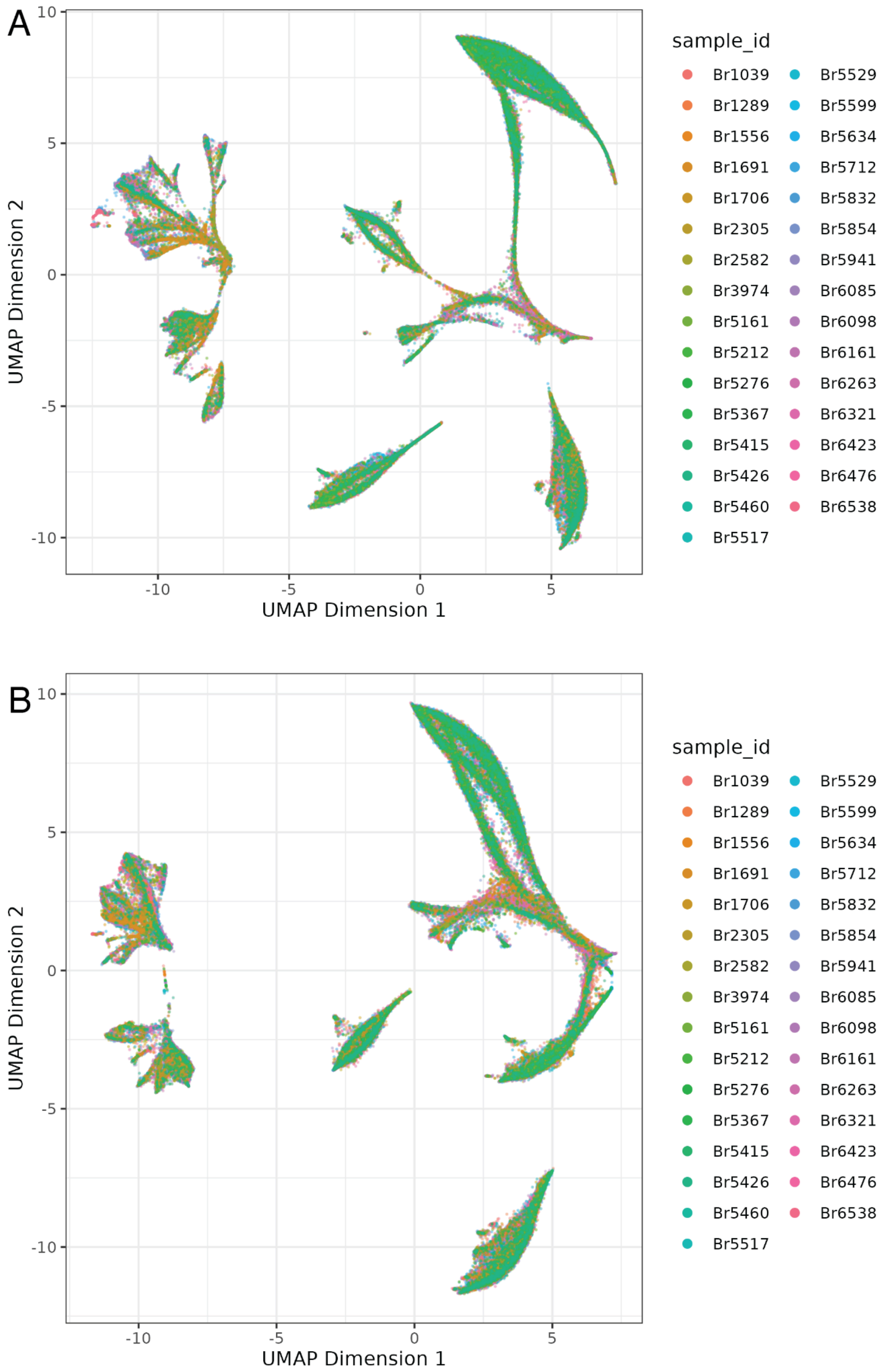


**Fig S17 Batch correction of snRNA-seq data visualized by UMAP. A**. UMAP of principal components (PCs) pre-batch correction with *harmony*[24] colored by donor. **B**. UMAP of PCs post-batch correction with *harmony*, colored by donor. Batch correction with *harmony* was configured to remove donor-specific effects (**Methods: snRNA-seq preliminary clustering and annotation**). Related to **Figure 2**.

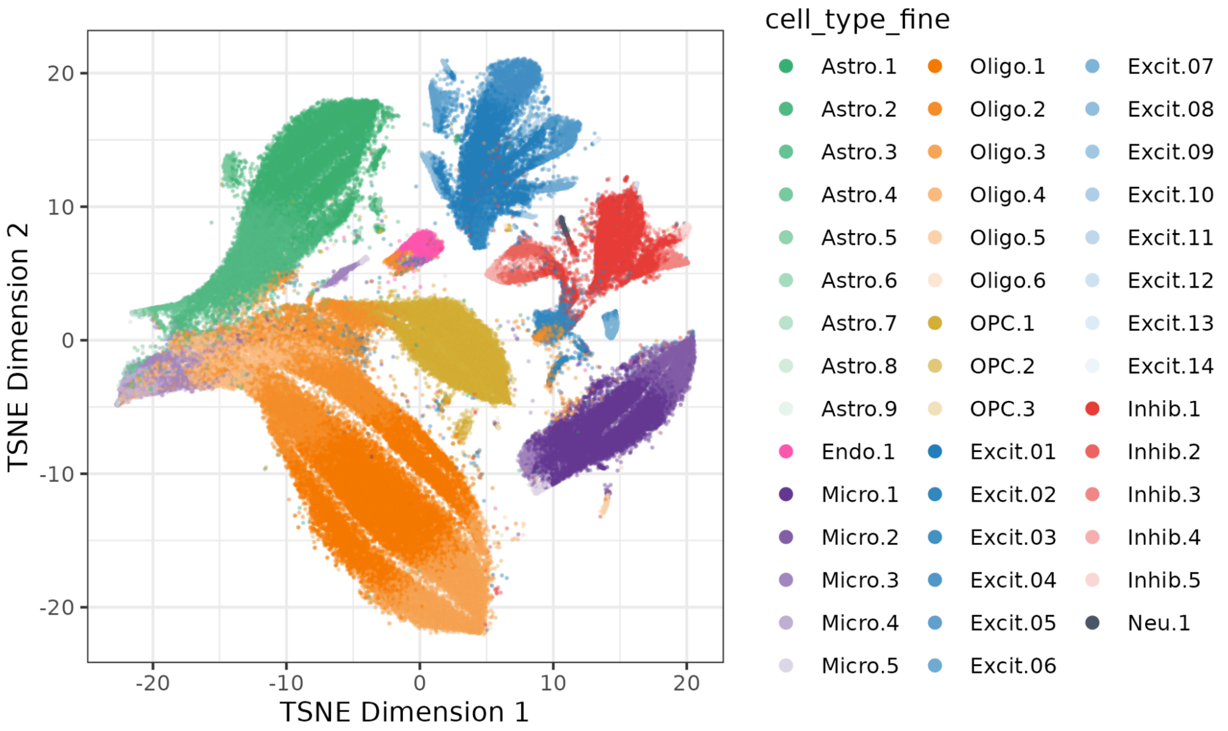


**Fig S18 Preliminary clustering and subcluster annotations of snRNA-seq data visualized by tSNE.** After batch correction with *harmony*, 44 preliminary clusters were identified using a shared nearest neighbor graph (*k* = 10) and `clusterwalktrap()`. They were annotated using *ScType*[119] (**Methods: snRNA-seq preliminary clustering and annotation**). Note that the cluster numbers used in this figure are preliminary cluster names and are different from the final 38 cell type subclusters described after quality control steps. Related to **Figure 2**.

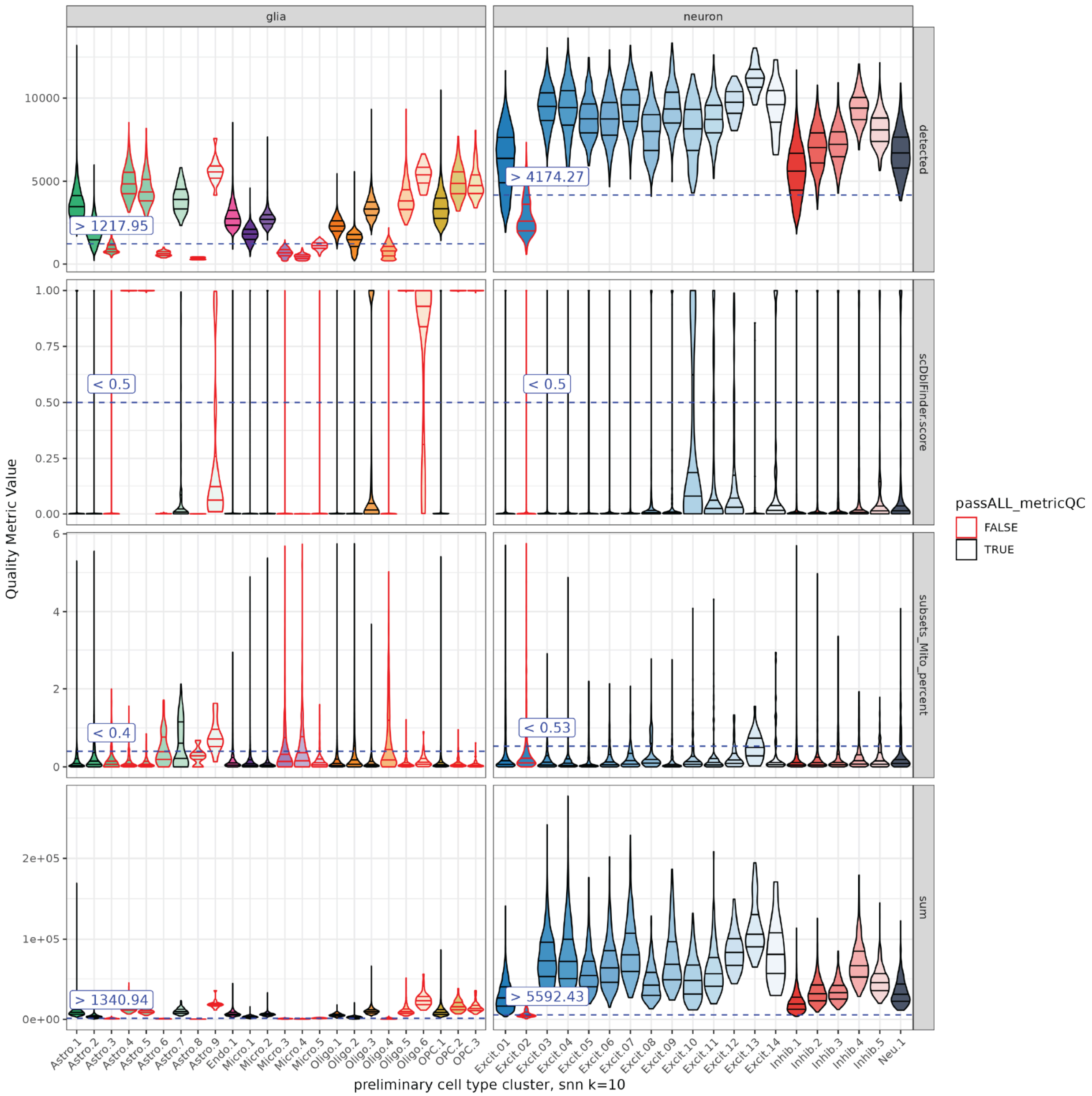


**Fig S19 Quality Control of preliminary clustering of snRNA-seq data.** Clusters were evaluated for low total UMIs (`sum`), low number of detected genes (`detected`), high percent of mitochondria expression (`subsets_Mito_percent`), and high doublet scores (`scDblFinder.score`). Cutoffs were determined by cell type class (neuron or non-neuron "glia") annotated as text box and dotted line. Clusters that failed one cutoff were outlined in red and were dropped due to their low quality in a cell type class context-aware manner (**Methods: snRNA-seq preliminary clustering and annotation**). In this step 125,771/140,119 (89.8%) nuclei from 29/44 clusters pass preliminary clustering quality control. Note that the cluster numbers used in this figure are preliminary cluster names and are different from the final 38 cell type subclusters described after quality control steps. Related to **Figure 2**.

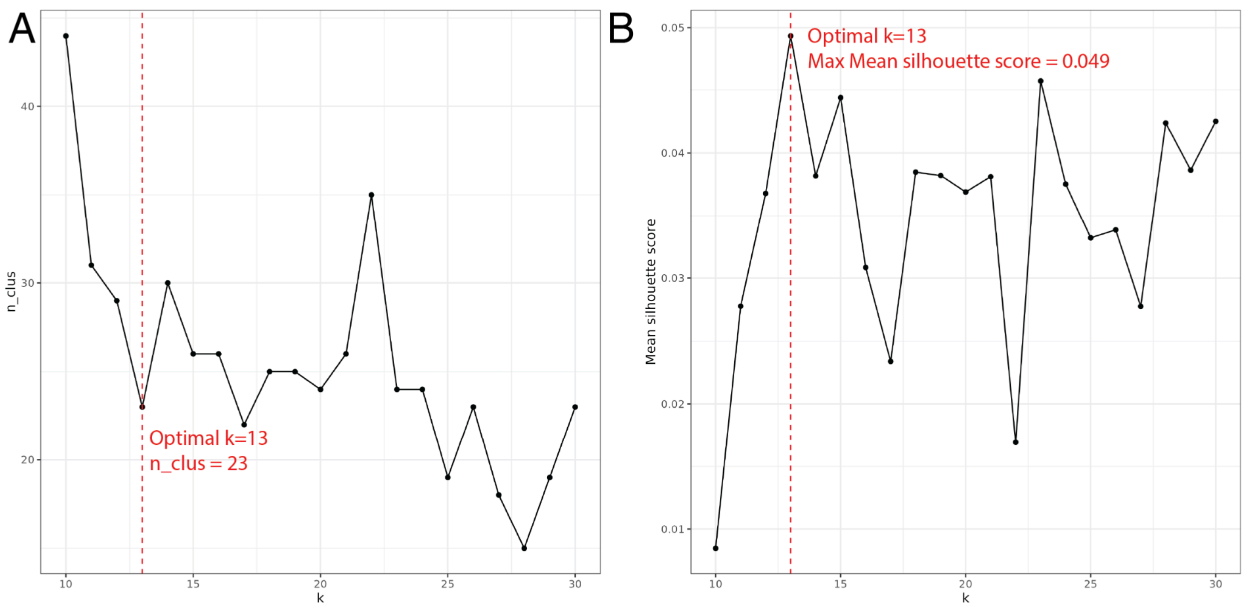


**Fig S20 Selection of optimal k for single-nucleus walktrap secondary clustering. A**. Number of clusters (`n_clus`) and **B**. mean silhouette score over values of *k*=10 to 30 (**Methods: snRNA-seq secondary optimal clustering and annotation**). Related to **Figure 2**.

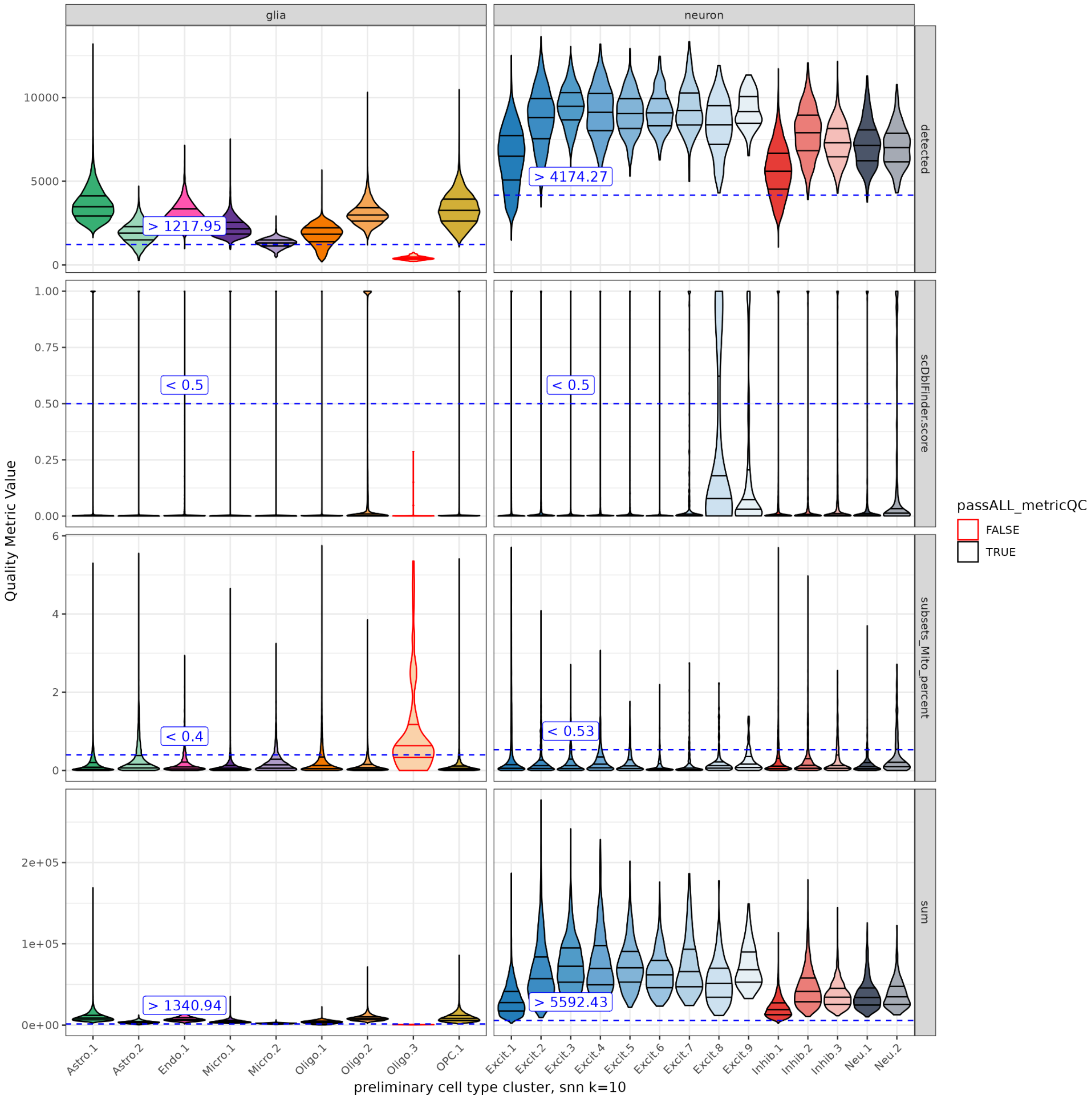


**Fig S21 Quality Control of secondary clustering of snRNA-seq data.** Clusters were evaluated for low total UMIs (`sum`), low number of detected genes (`detected`), high percent of mitochondria expression (`subsets_Mito_percent`), and high doublet scores (`scDblFinder.score`). Cutoffs were determined by cell type class (neuron or non-neuron "glia"), and annotated text box and dotted line. Clusters that failed one cutoff were outlined in red and were dropped due to their low quality. In this step 125,683/125,771 (99.9%) nuclei from 22/23 clusters pass secondary clustering quality control. (**Methods: snRNA-seq secondary optimal clustering and annotation**). Note that the cluster numbers used in this figure are preliminary cluster names and are different from the final 38 cell type subclusters described after quality control steps. Related to **Figure 2**.

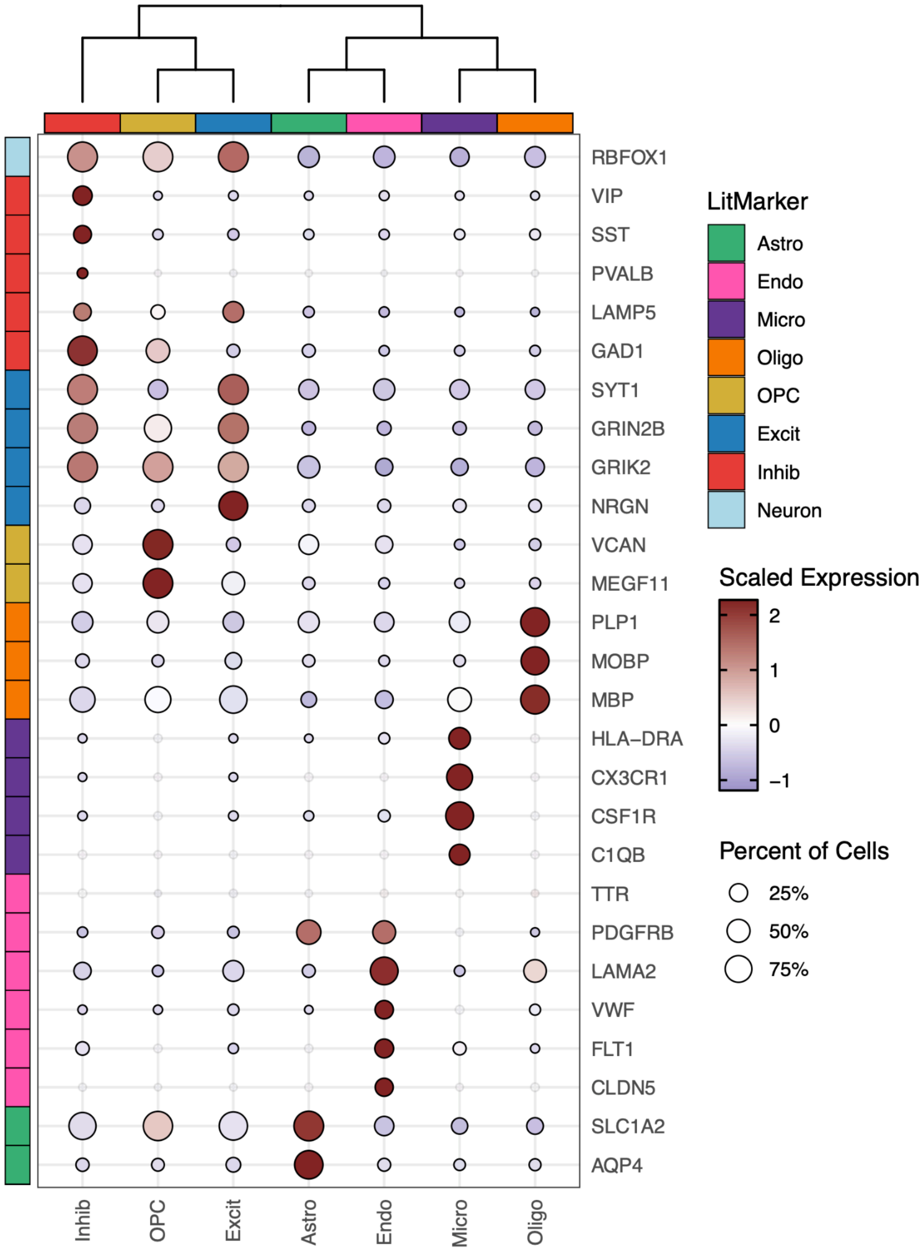


**Fig S22 Expression dotplot of literature marker genes in secondary optimal clustering broad cell types.** Columns are post preliminary-QC nuclei at broad cell type level. Log-normalized gene expression is scaled and centered. Marker genes sourced from other ERC snRNA-seq literature[19,38]. Related to **Figure 2**.

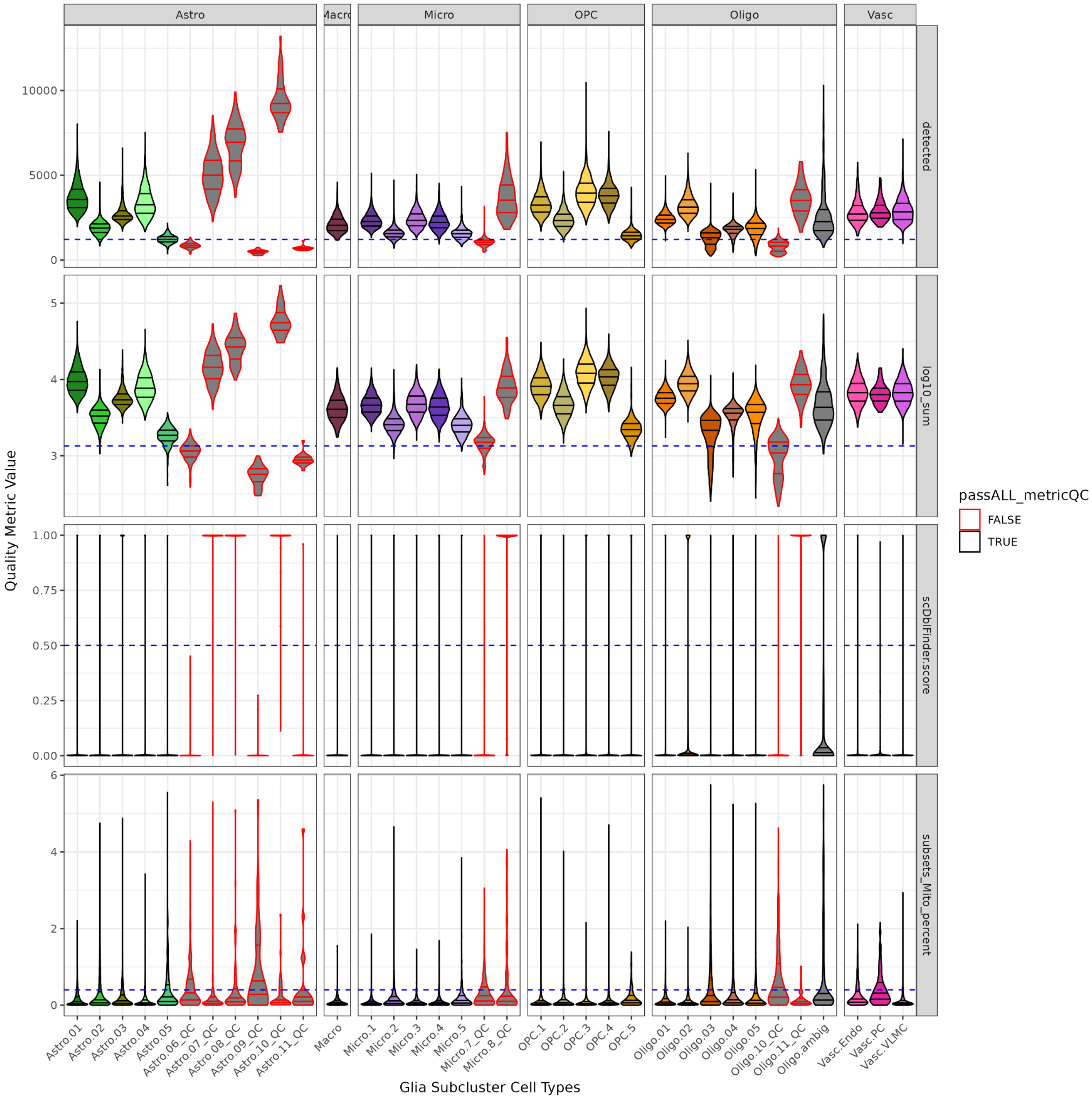


**Fig S23 Quality Control of subclustering of snRNA-seq non-neuronal data.** Post secondary cluster-QC, and non-neuronal subclustering, non-neuronal clusters were again evaluated for low total UMIs (`sum`), low number of detected genes (`detected`), high percent of mitochondria expression (`subsets_Mito_percent`), and high doublet scores (`scDblFinder.score`). The same cutoffs from the secondary clustering QC were applied, indicated by the dotted line. Clusters that failed one cutoff were outlined in red and were dropped due to their low quality. In this step 94,990/98,669 (96.3%) non-neuronal nuclei from 24/34 subclusters pass non-neuronal subclustering quality control. (**Methods: snRNA-seq non-neuronal subclustering and annotation**). Related to **Figure 2**.

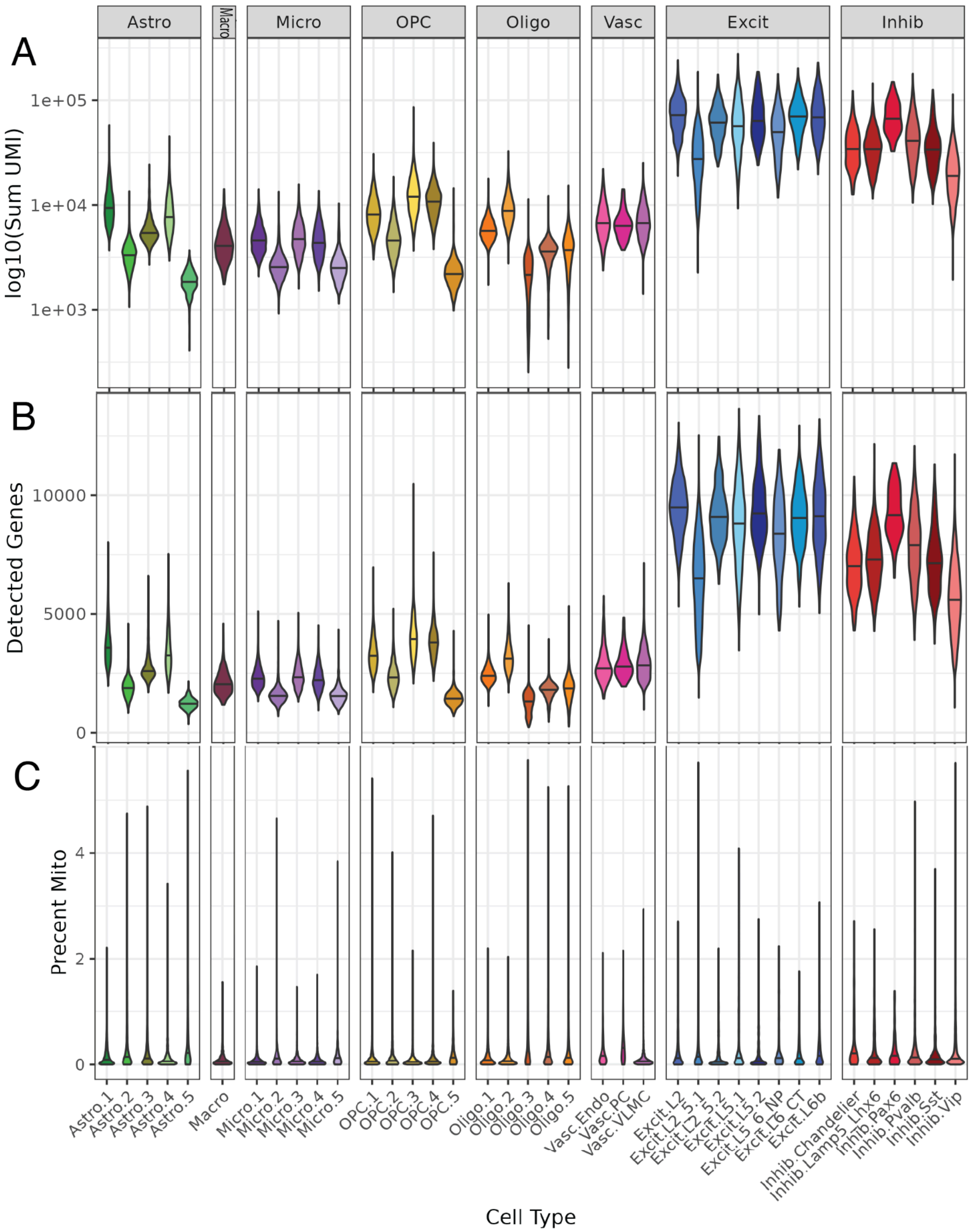


**Fig S24 snRNA-seq final fine subcluster QC metrics.** Violin plots of **A.** $\log_{10}$ sum UMI, **B.** Number of detected genes, and **C.** percent of mitochondria expression (percent Mito) for each fine subcluster. In total 122,004 nuclei in 38 subclusters pass all steps of quality control and clustering (**Methods: snRNA-seq non-neuronal subclustering and annotation**). Related to **Figure 2**.

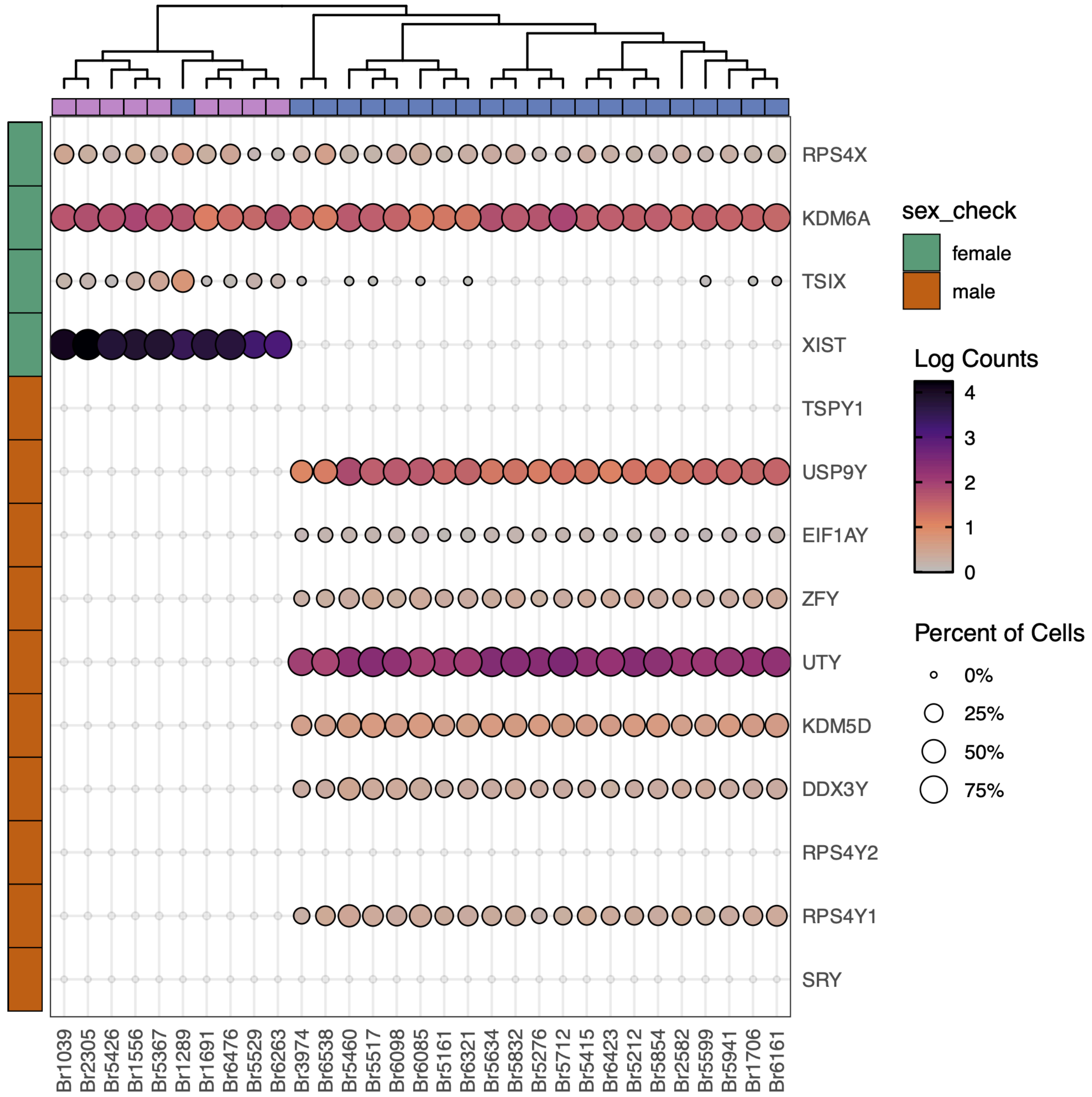


**Fig S25 Expression of sex genes in snRNA-seq data.** Centered and scaled log-normalized gene expression of select sex marker genes were visualized using *scDotPlot*[136]. Donor sex was annotated in the columns: pink for female, blue for male. This identified expression of female sex genes on donor Br1289 who was a male donor. Upon further verification, a swap in the case donor selection table was discovered: Br1285 is from a female African ancestry control *APOE* E3/E4 donor with age of death at 17.27 years. Given its outlier age Br1285 (labelled as Br1289) was not considered in differential expression analyses. (**Methods: Donor demographics check**). Related to **Figure 2**.

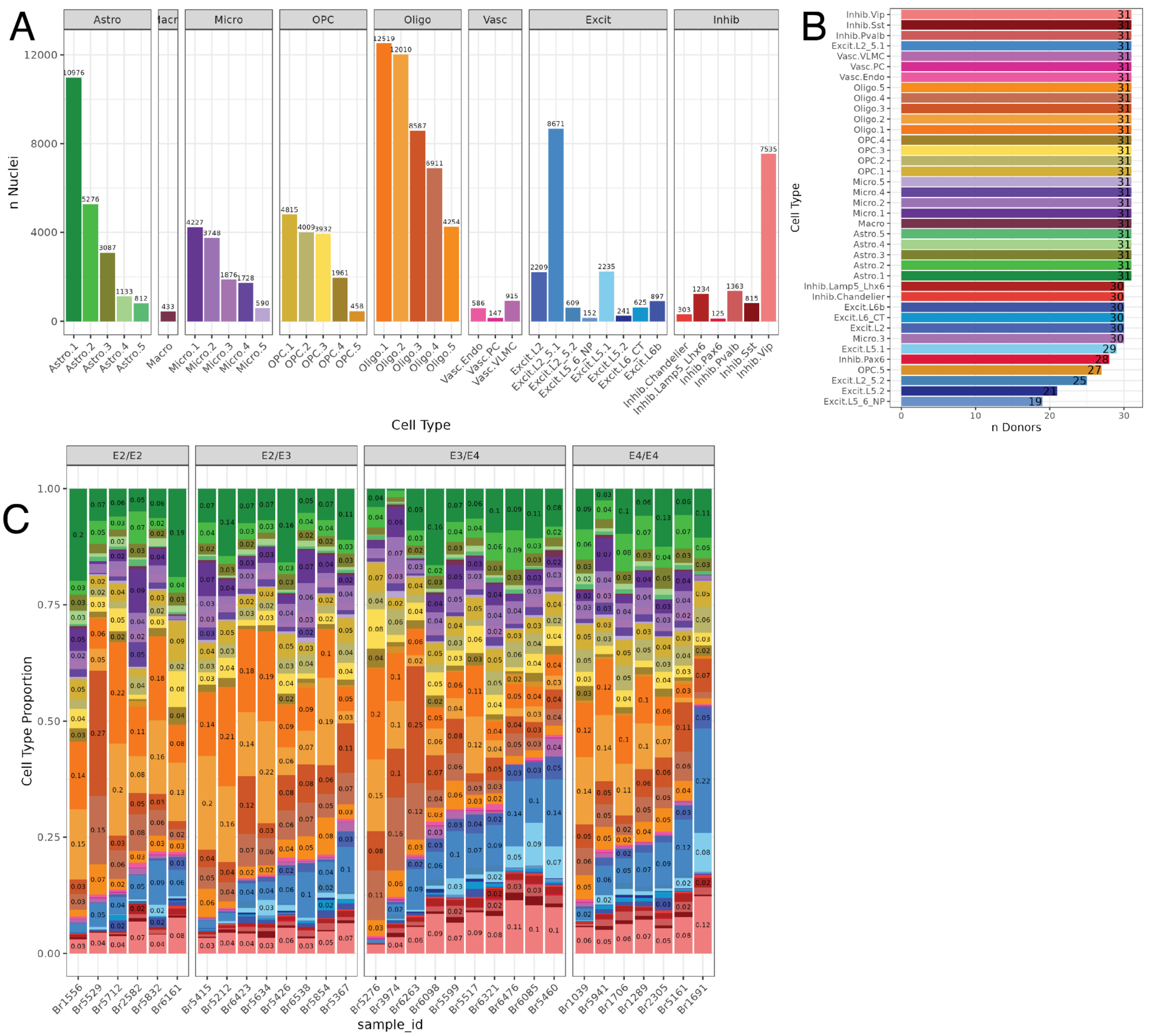


**Fig S26 Cell type composition of snRNA-seq dataset. A**. Barplot of the number of nuclei in each fine subcluster. Note that several subclusters had 620 nuclei (<20 nuclei/donor) which can limit power for pseudobulked DGE results[74]. **B.** Barplot of the number of donors contributing to each fine subcluster population. **C**. Fine subcluster composition barplots for the 31 samples, organized by *APOE* genotype, subcluster colors correspond to *A* (**Methods: snRNA-seq non-neuronal subclustering and annotation**). Related to **Figure 2**.

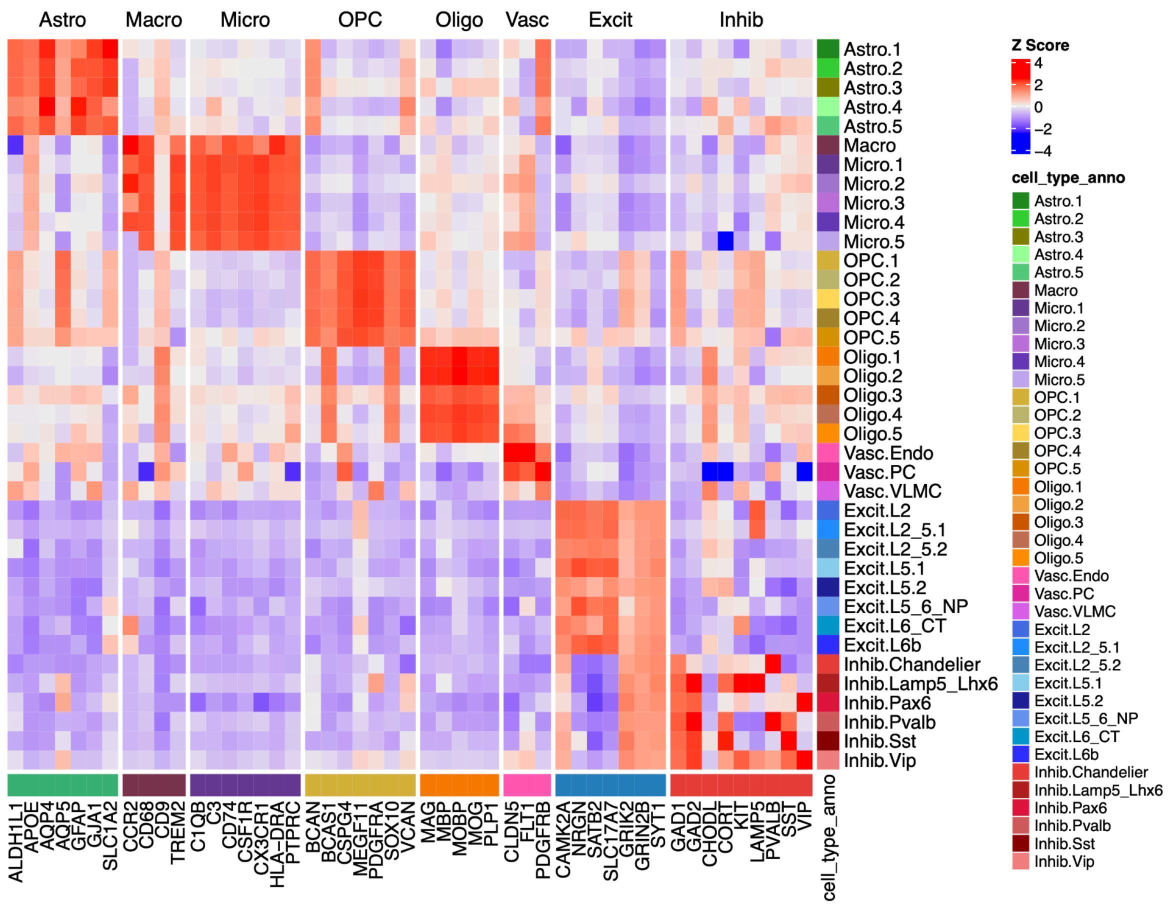


**Fig S27 Marker gene expression across fine subclusters.** Heatmaps of scaled and centered mean pseudobulked logcounts for select cell type marker genes from the literature (**Methods: snRNA-seq non-neuronal subclustering and annotation**). Related to **Figure 2**.

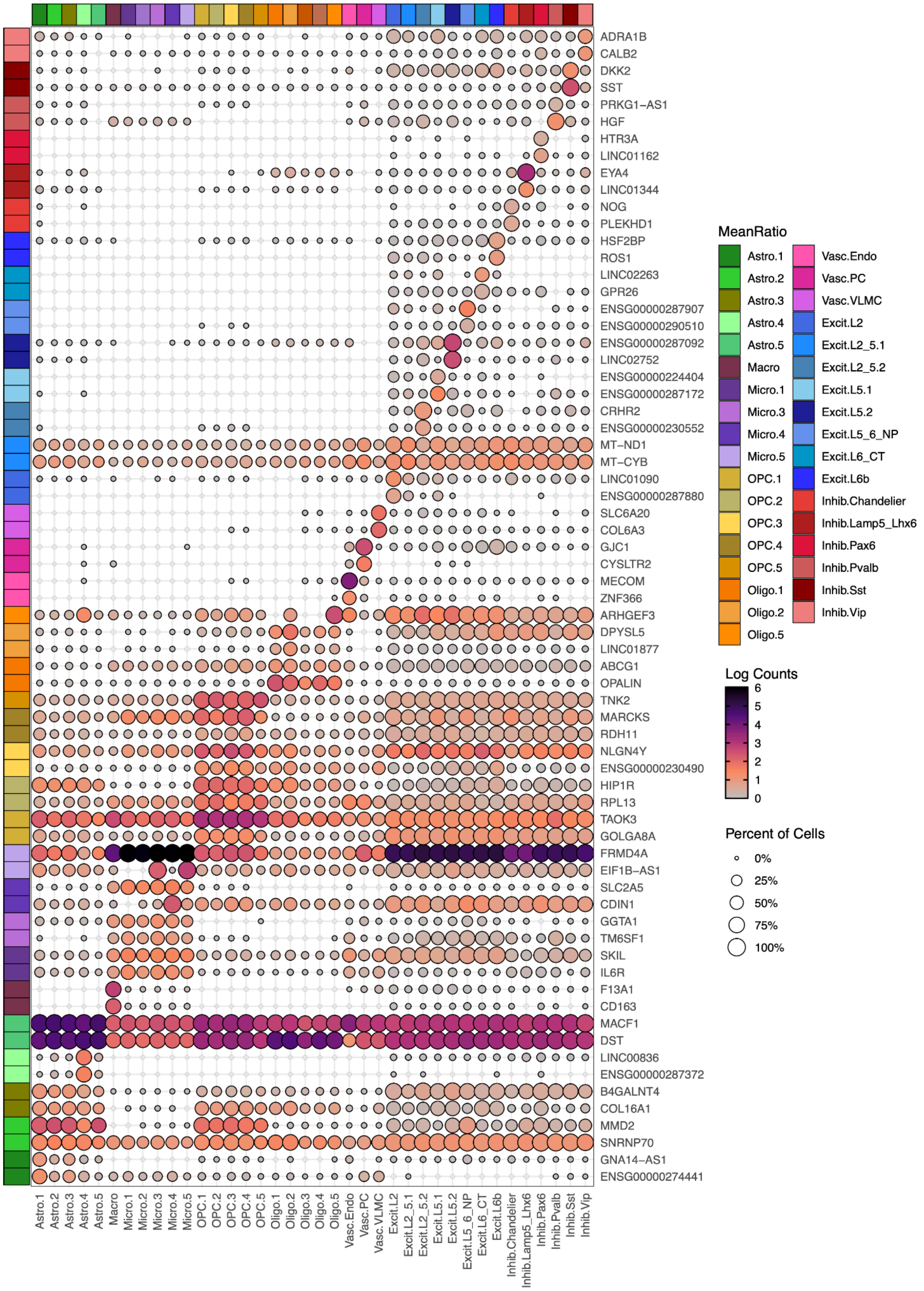


**Fig S28 Expression dotplot of *MeanRatio* marker genes for fine subclusters.** Rows pertain to the top two *MeanRatio* marker genes with `MeanRatio > 1`$^{40}$ for each fine subcluster (columns). Dot color is log-normalized gene expression, dot size is percent non-zero expression (**Methods: snRNA-seq non-neuronal subclustering and annotation**). Related to **Figure 2**, **Table S6**.

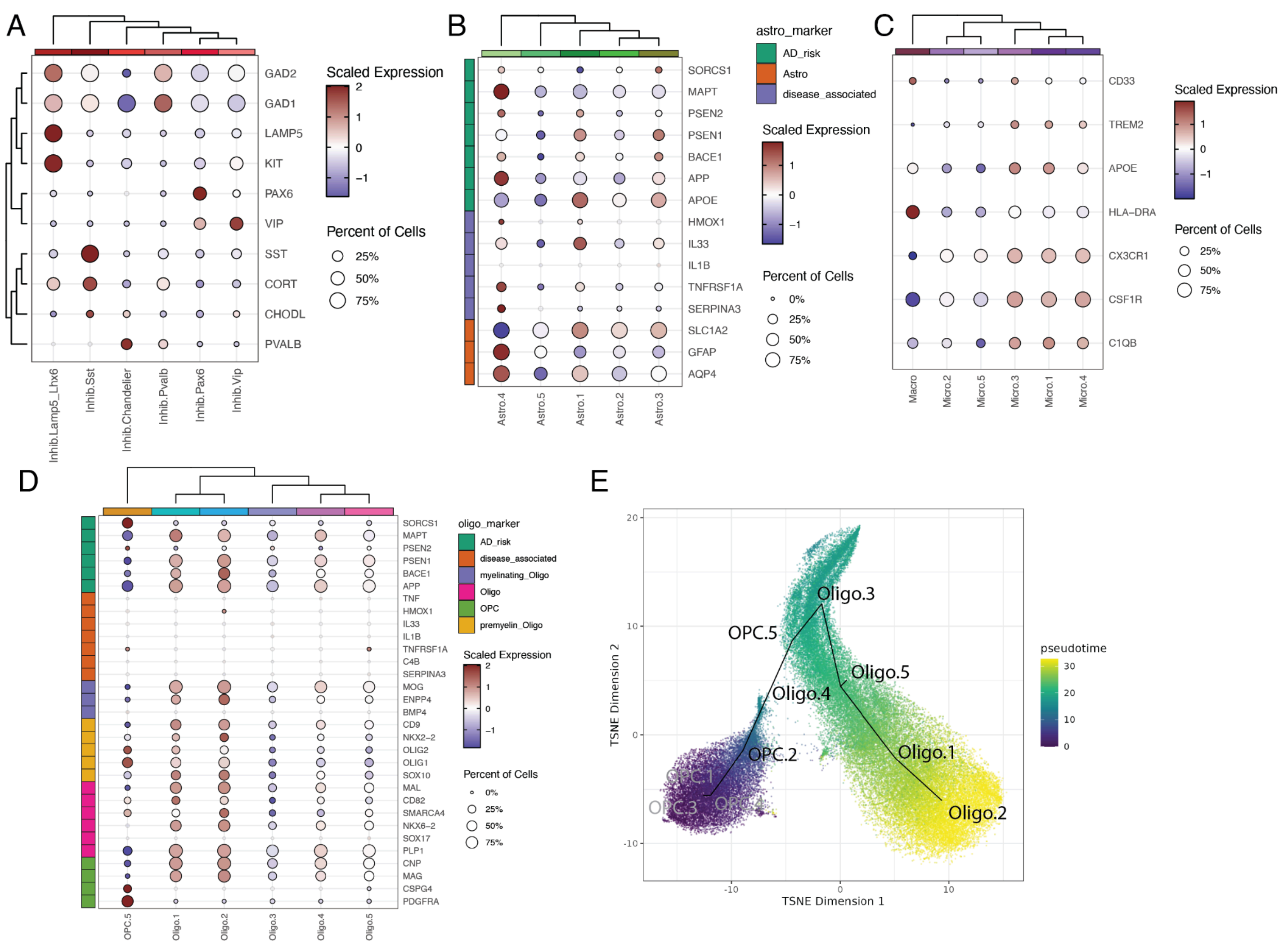


**Fig S29 snRNA-seq subcluster expression of subtype and disease state marker genes from literature for subcluster annotation. A-D**. Centered and scaled log-normalized gene expression of cell type markers, disease state associated markers genes[20] (**A-C**), and AD risk genes[138] (**D**), visualized using *scDotPlot*[136]. **A.** Subclusters of Inhib neurons over broad (*GAD1* and *GAD2*) and subcluster markers. **B.** Astrocyte subclusters with annotated genes associated with disease state. **C.** Microglial and Macro subclusters; *TREM2* has been associated with disease-activated microglia[16]. **D.** Oligodendrocyte subclusters and closely related OPC.5 over oligodendrocyte related marker genes, including disease state associated marker genes. **E.** tSNE plot of OPC and Oligo subclusters with trajectory analysis, shaded with pseudotime (**Methods: Oligodendrocyte subcluster trajectory analysis**). Related to **Figure 2**, **Figure 5.**

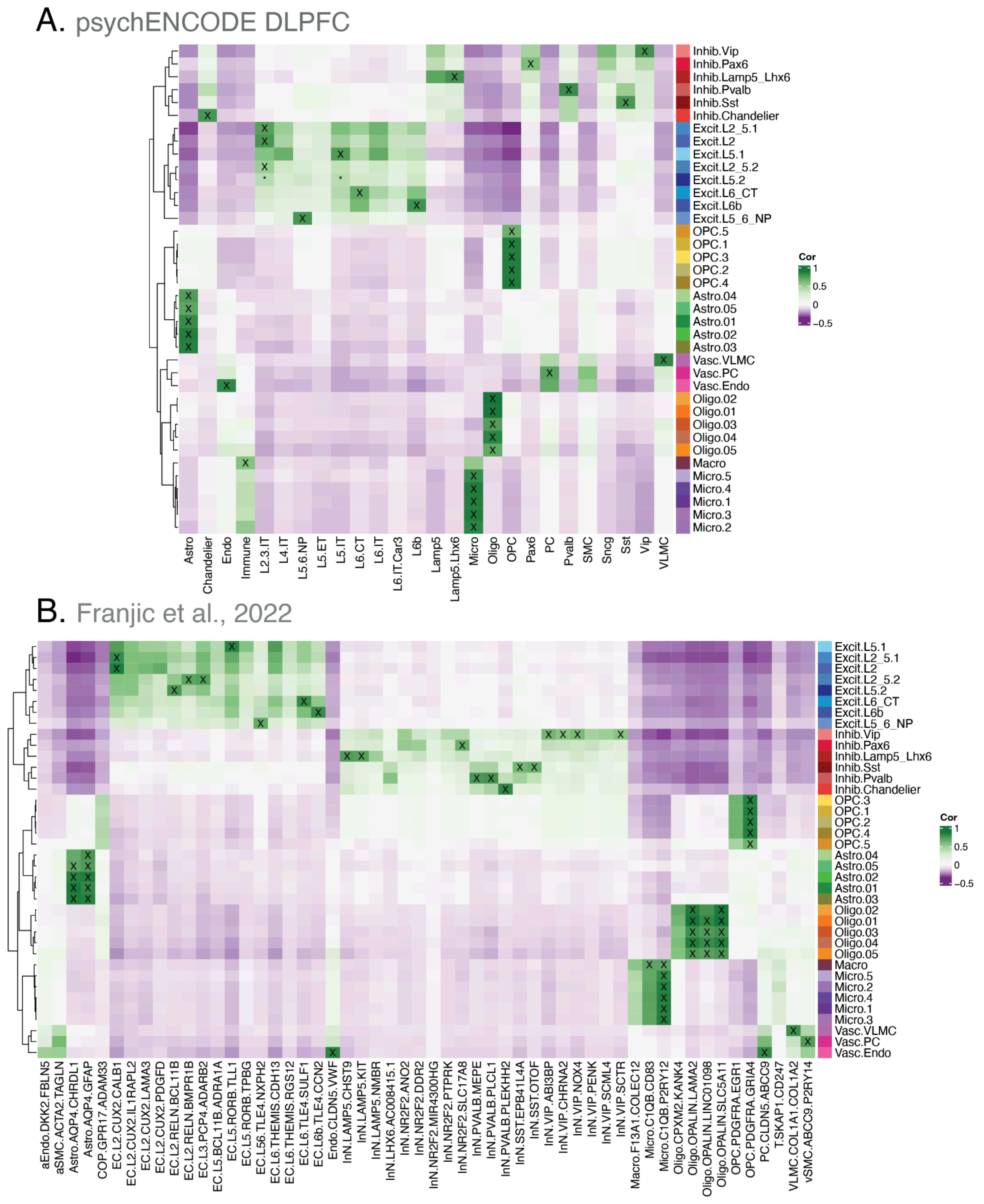


**Fig S30 snRNA-seq subcluster registration to external human cortex datasets. A**. *spatialLIBD* spatial registration heatmap[26–28] between our fine resolution snRNA-seq clusters and the psychENCODE LIBD DLPFC snRNA-seq data[39]. **B**. *spatialLIBD* spatial registration heatmap[26–28] against the Franjic et al. ERC fine subcluster populations[29] (**Methods: snRNA-seq non-neuronal subclustering and annotation**). High confidence matches (cor > 0.5, merge ratio = 0.1) are marked with an "X" low confidence matches are marked with "*". Cluster colors in all panels match. Related to **Figure 2**.

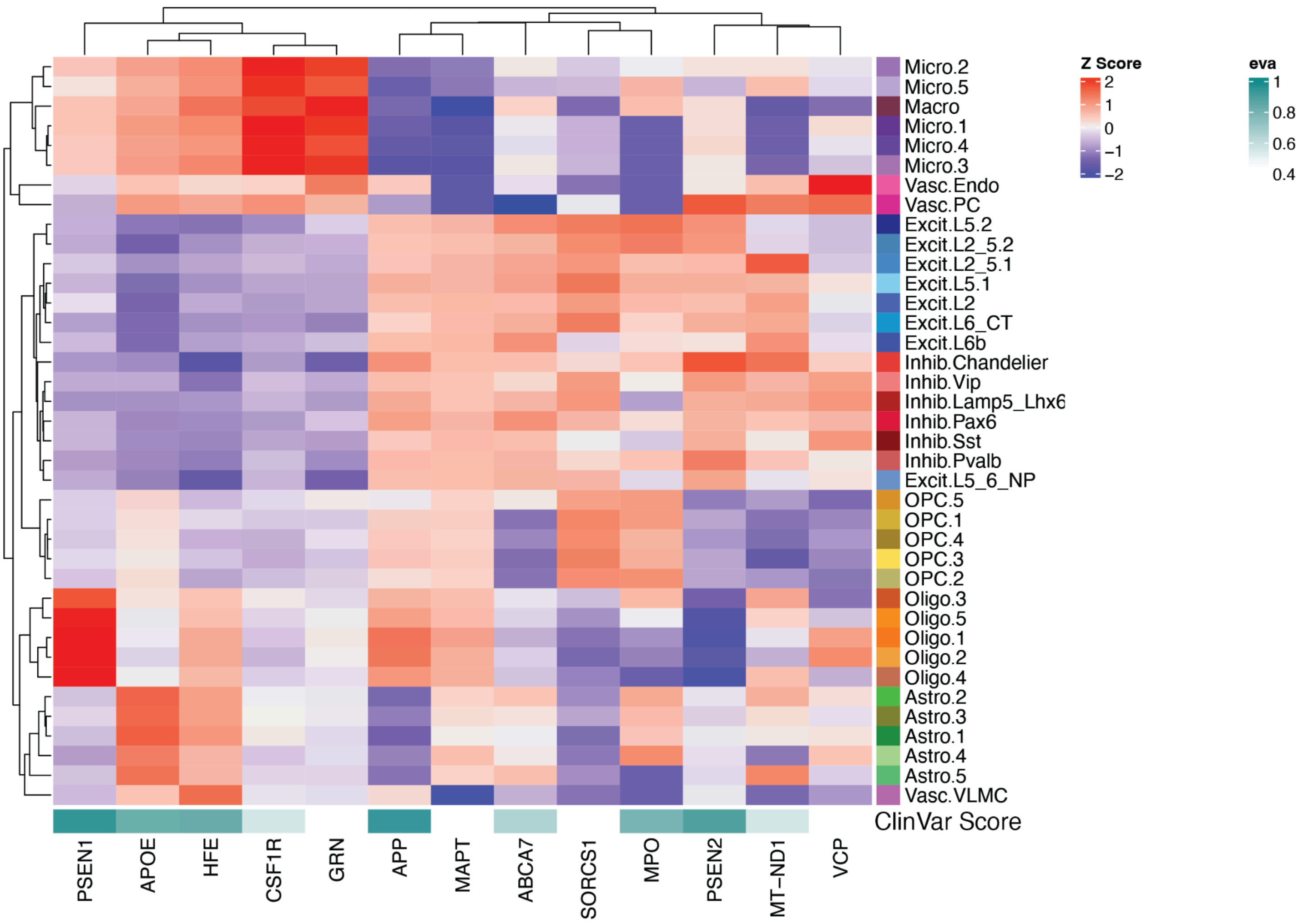


**Fig S31 Expression of AD risk genes across single cell clusters.** Heatmaps of scaled and centered mean pseudobulked logcounts expression of *ClinVar* AD risk genes across cell type subclusters. *ClinVar* genes sourced from *OpenTargets*, where `eva` stands for European Variation Archive and in this case is the association score for AD from EVA (available in *OpenTargets* through *ClinVar*)[110] (**Methods: AD risk gene set**). Related to **Figure 2**.

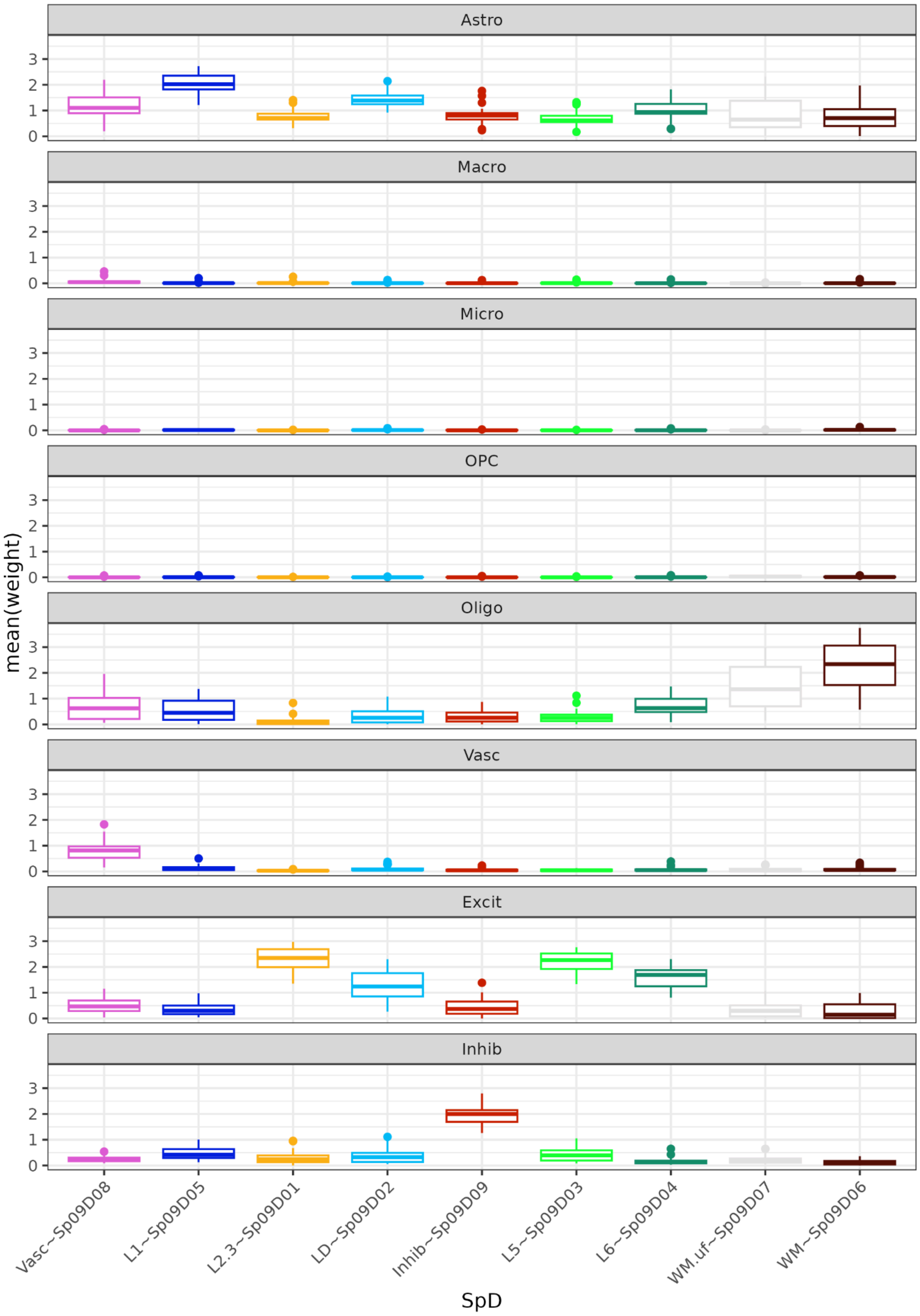


**Fig S32 Estimated spot cell type frequency with spot deconvolution**. Boxplots of Mean *RCTD* cell type proportions (weights) by donor (*Y*-axis) over SpDs (*X*-axis). Related to **Figure 2** and **Table S7**.

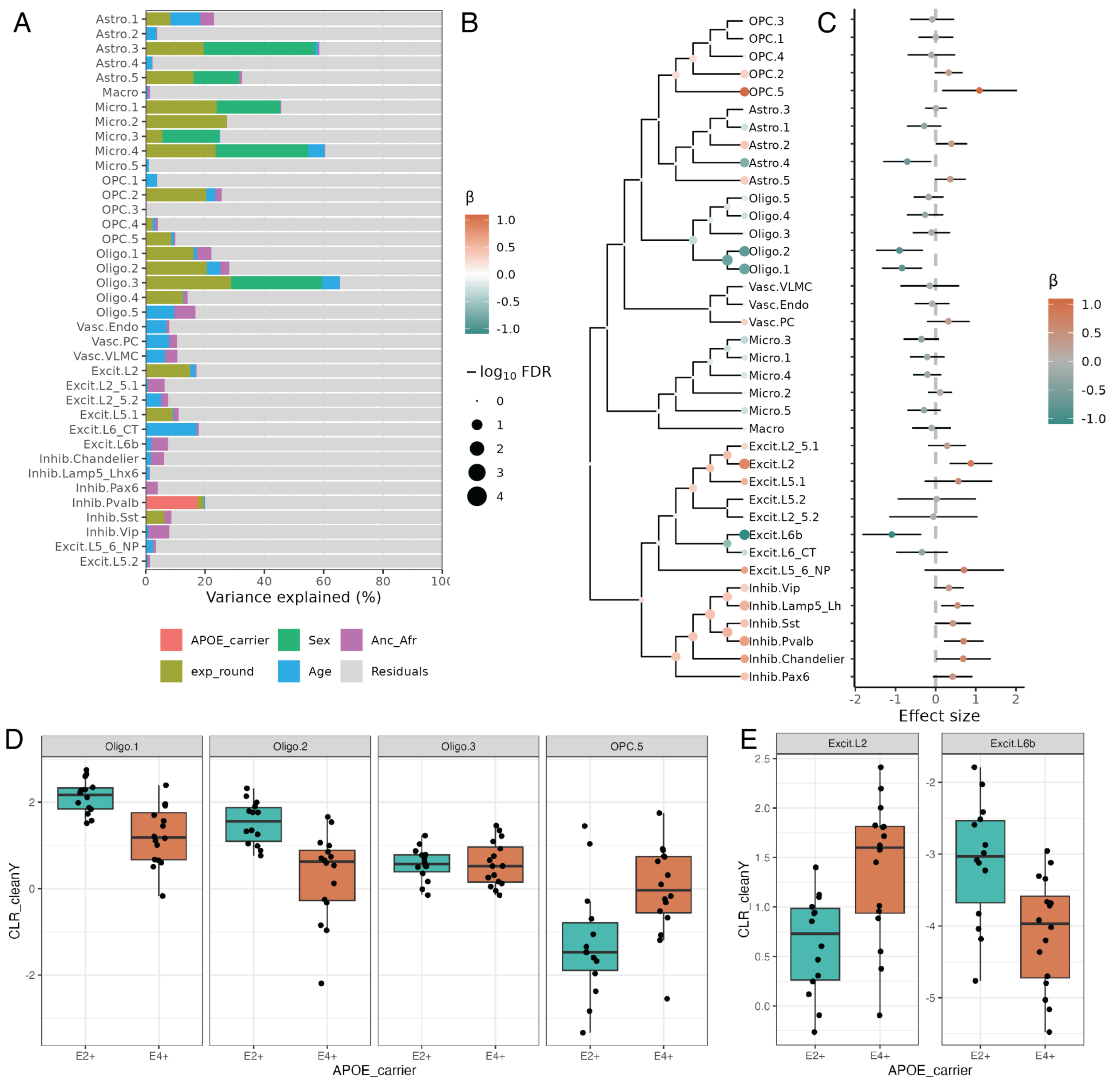


**Fig S33 Differential proportion analysis of snRNA-seq fine subclusters. A**. Barplot showing the percent of proportion variance explained by different variables. **B**. Hierarchical tree of snRNA-seq subclusters, node color indicates the estimated effect size from regression analysis of *APOE* carrier status (E2+ vs. E4+) at leaves (fine subclusters) and multivariate testing at internal nodes. Node point size represents FDR for each test. **C**. Forest plot of effect size from *APOE* carrier test, effect size > 0 represents higher frequency of the subcluster in E4+ donors, <0 higher frequency in E2+ donors. The color corresponds to the effect size (β) and bars indicate the 95% confidence interval. **D.** Box plots of the centered log ratio (CLR) of subcluster frequency for select Oligo and OPC populations across *APOE* carrier status, confounding variables regressed out with `cleaningY`, protecting `APOE_carrier`. **E.** `cleaningY` CLR boxplots for select Excit populations (**Methods: Differential proportion analysis**). Related to **Figure 2**, **Table S8**.

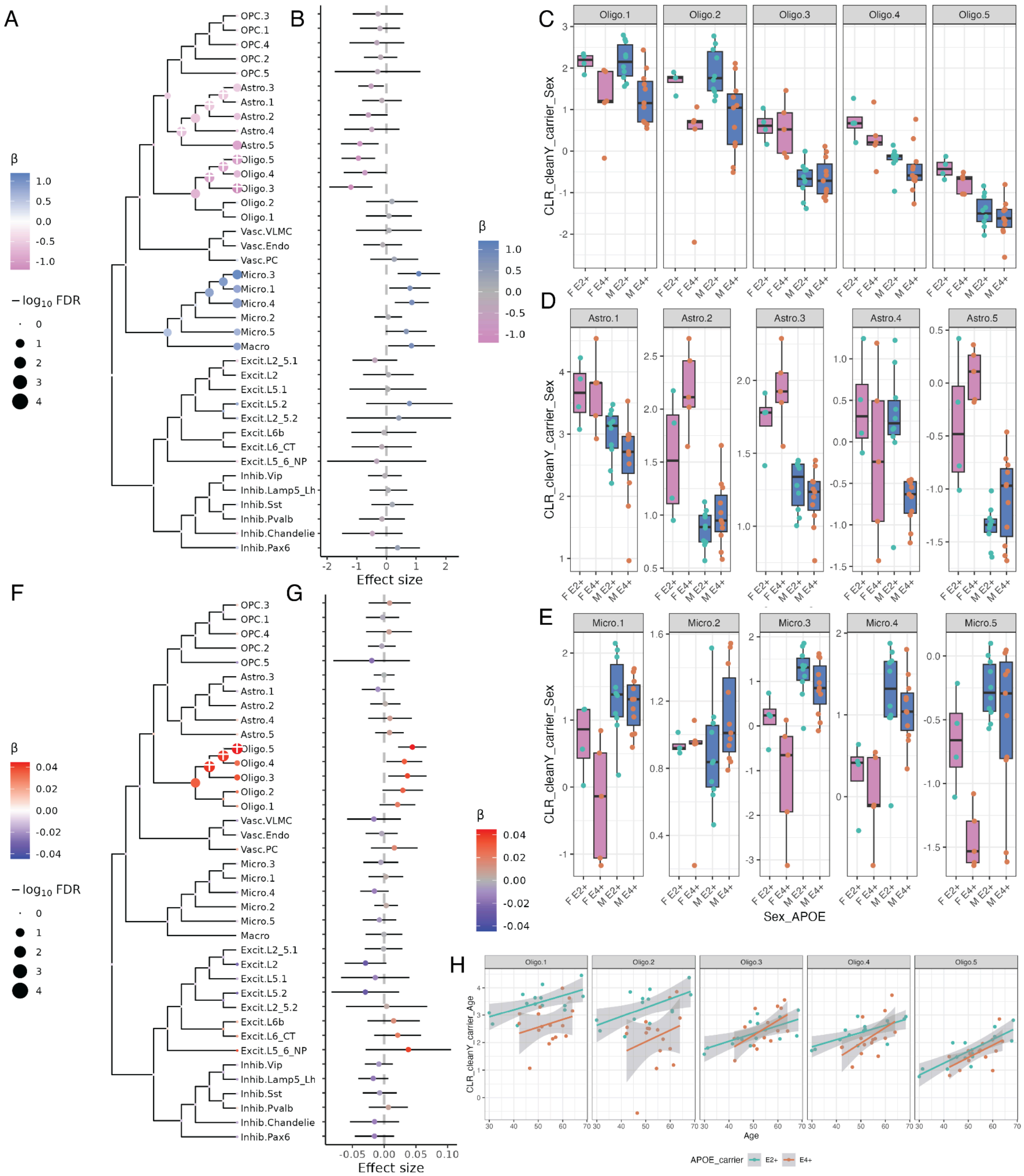


**Fig S34 Differential proportion analysis of snRNA-seq fine subclusters for sex and age. A**. Hierarchical tree of fine subclusters, node color indicates the estimated effect size from regression analysis of sex (female vs. male) at leaves (fine subclusters) and multivariate testing at internal nodes. node point size represents FDR for each test, FDR < 0.05 is marked with a "+". **B**. Forest plot of effect size from sex test, effect size > 0 represents higher frequency of the snRNA-seq subcluster in male donors, <0 higher frequency in female donors. The color corresponds to the effect size (β) and bars indicate the 95% confidence interval. **C.** Box plots of the centered log ratio (CLR) of subcluster frequency for select Oligo subclusters across sex and *APOE* carrier status, confounding variables regressed out with `cleaningY`, protecting `APOE_carrier` and `sex`. Oligo.1 and Oligo.2 proportions divided by carrier status and had higher proportions in E2+ carriers, Oligo.3 divided by sex with similar proportions between *APOE* carriers, and Oligo.4 and Oligo.5 were affected linearly by combinations between sex and *APOE* carrier. **D.** `cleaningY` CLR boxplots for Astro and **E.** Micro sub-populations. Astrocytes were often more frequent in females, but subcluster frequencies at times interacted with *APOE* carrier status, such as Astro.4 that had a lower frequency in E4+ carrier males (*D*). Nominal increases (FDR>0.05, p<0.05) in microglia proportions were observed in males compared to females for several microglia subclusters (*E*). **F**. Hierarchical tree of subclusters for regression analysis of Age. **G**.

Forest plot of effect size from Age. **H.** Scatter plot of age vs. `cleaningY` CLR (protecting `APOE_carrier` and `age`), trend lines from `geom_smooth(method = "lm")` (**Methods: Differential proportion analysis**). Related to **Figure 2**, **Table S8**.

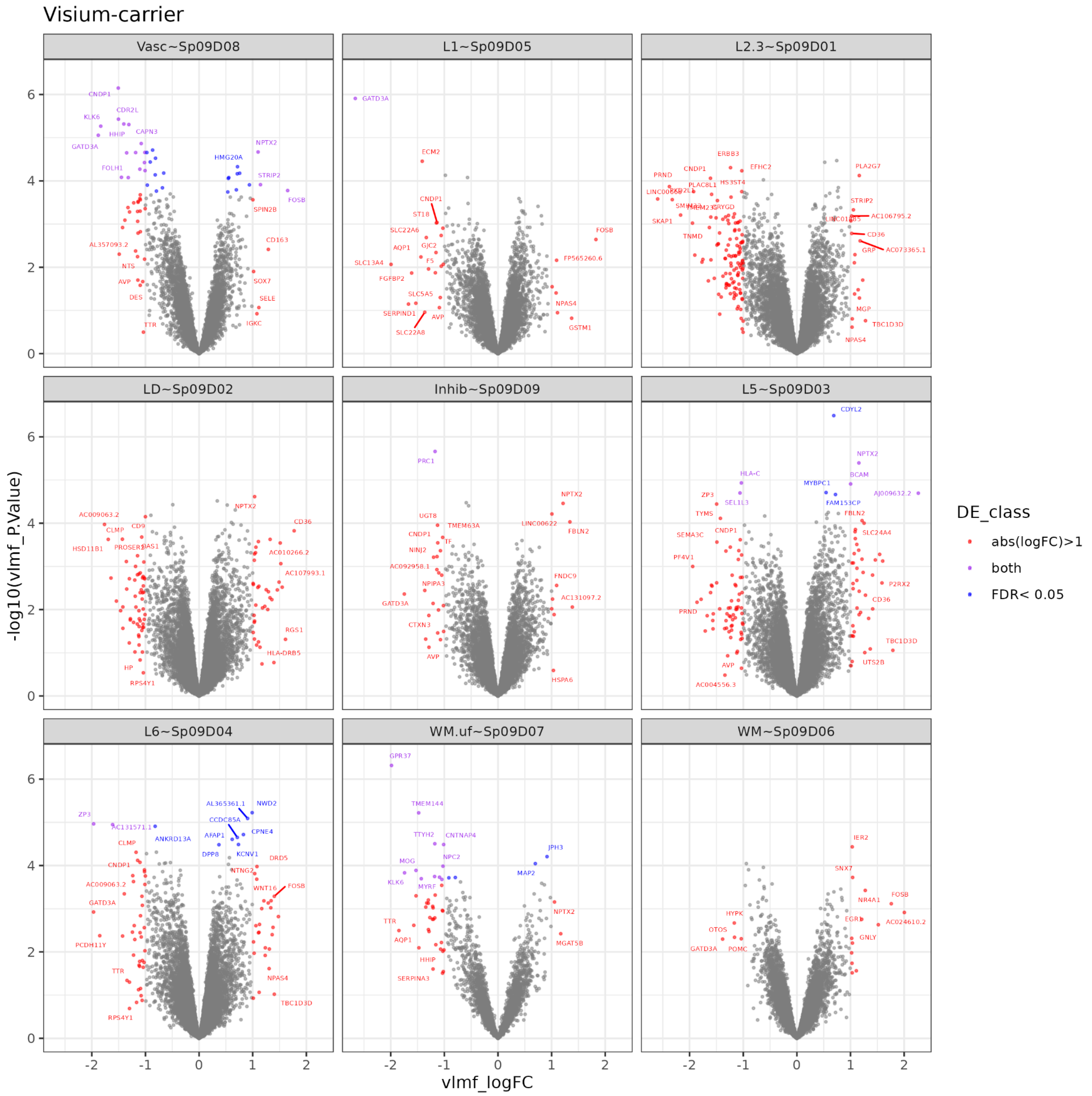


**Fig S35 Differential expression between *APOE* E2+ and E4+ carriers for SRT SpDs.** Volcano plots for carrier DE analysis for all spatial domains. Genes with FDR<0.05 or absolute $\log_2$ fold change (logFC) > 1 are indicated with point color. (**Methods: Differential expression analysis**). Related to **Figure 3, Table S9**.

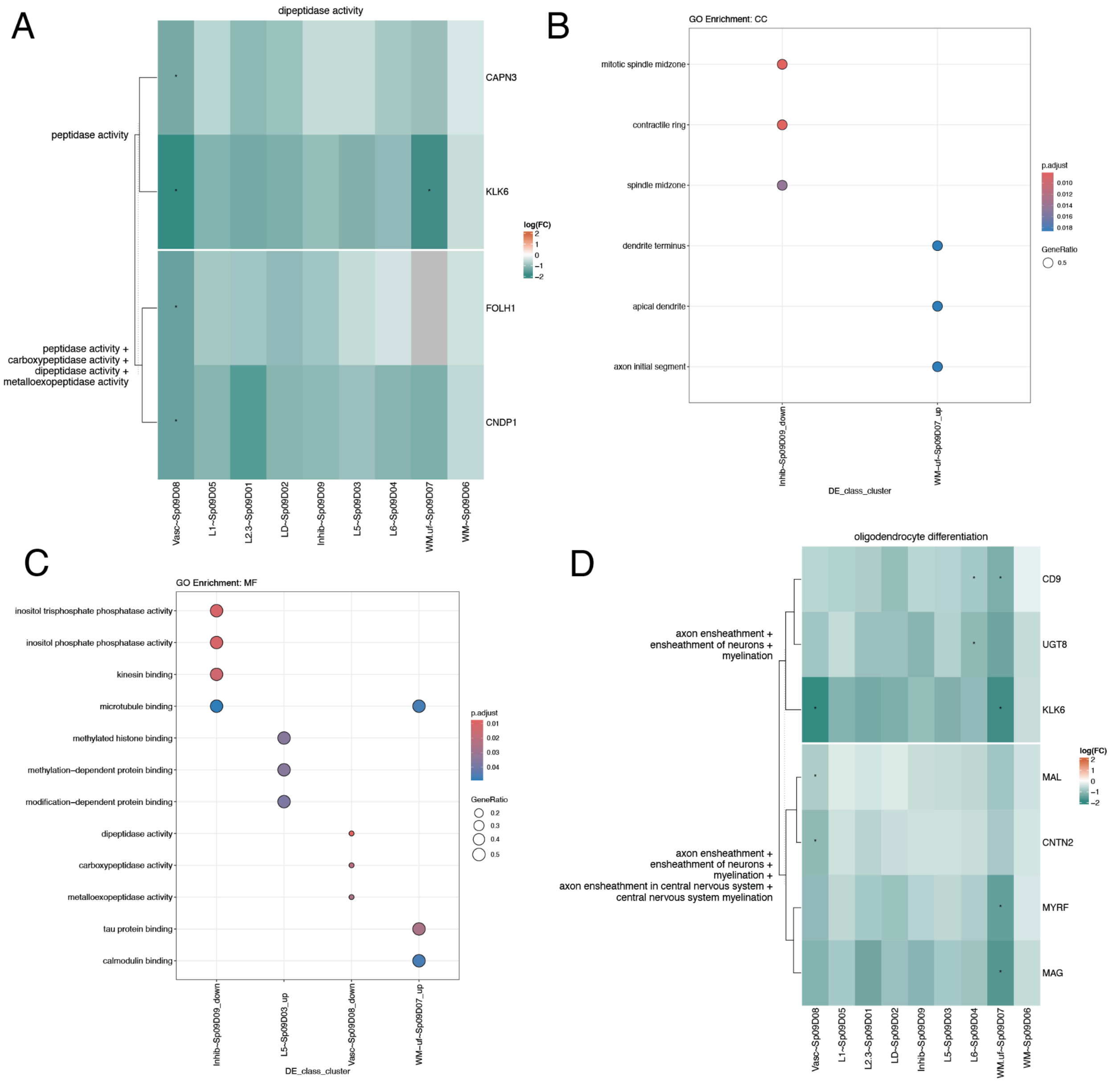


**Fig S36 Gene ontology analysis for SRT *APOE* carrier differential expression. A.** Differential expression logFC heatmap for genes associated with molecular function (MF) parent GO term “dipeptidase activity” (**Methods: Gene ontology overrepresentation analysis**). Related to **Figure 3**. **B-C.** *clusterProfiler*[127] GO dot plot for “Cellular Component” (**B**), and “Molecular Function” (**C**). Gene ratio is the number of genes in each gene set annotated in a term (y-axis) over the total in the respective gene set. p.adjust is the FDR-adjusted p-value. **D.** Differential expression logFC heatmap for genes associated with biological process (BP) parent term “oligodendrocyte differentiation" (**Methods: Gene ontology overrepresentation analysis**). Related to **Figure 3**, **Table S10**.

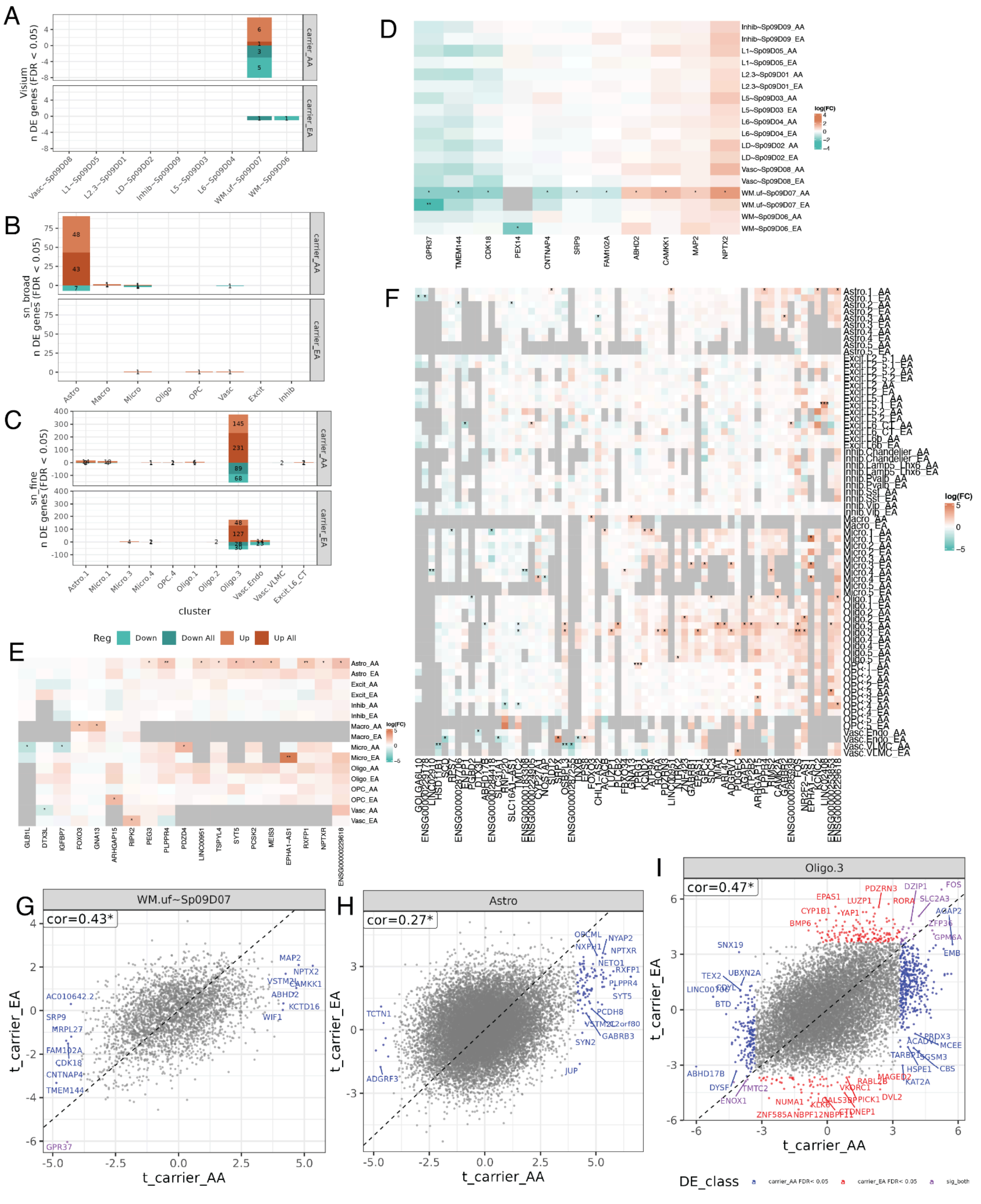

**Fig S37 Ancestry-specific *APOE* carrier differential expression analysis. A-C.** Barplots of number of differentially expressed genes in ancestry-specific analysis by *APOE* carrier (E2+ vs. E4+) for Visium SpDs (**A**), snRNA-seq cell types at broad resolution (**B**), and snRNA-seq fine subclusters (**C**). **D-F**.Heatmaps of log fold change values for top differentially expressed genes across Visium SpDs (**D**), broad snRNA-seq cell types (**E**), and fine snRNA-seq subclusters (**F**). **G-I**. Scatter plots comparing *t*-statistics from AA and EA specific DE analysis for WM.uf~$Sp_9D_7$ (**G**), road cell type Astro (**H**), and fine subcluster Oligo.3 (**I**, 24 DEGs were shared across both ancestry groups, 209 were EA-specific, and 509 were AA-specific). Related to **Figure 3**, **Figure 4**, **Figure 5, Table S9**, **Table S11**, **Table S13**.

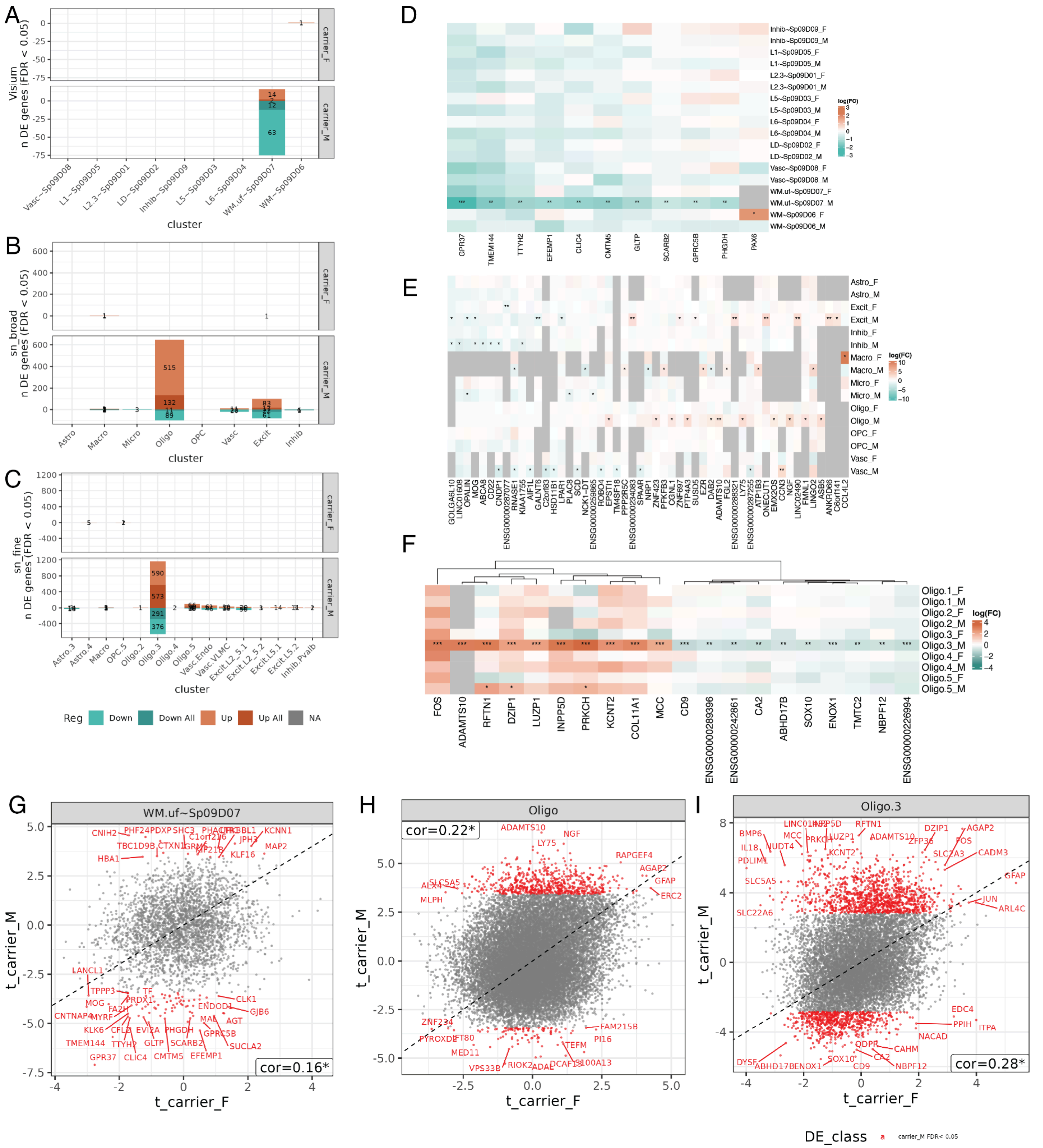

**Fig S38 Sex specific *APOE* carrier differential expression analysis. A-C.** Barplots of number of differentially expressed genes in ancestry-specific analysis by sex for Visium SpDs (**A**), cell types at broad resolution (**B**), and fine subclusters (**C**). **D-F**. Heatmaps of $\log_2$ fold change values for top differentially expressed genes between *APOE* E4+ and E2+ carriers across Visium SpDs (**D**), broad cell types (**E**), and fine Oligo subclusters (**F**). **G-I**. Scatter plots comparing *t*-statistics from AA and EA specific DE analysis for WM.uf~$Sp_9D_7$ (**G**), broad cell type Oligo (**H**), and fine subcluster Oligo.3 (**I**). Related to **Figure 3**, **Figure 4**, **Figure 5, Table S9**, **Table S11**, **Table S13**.

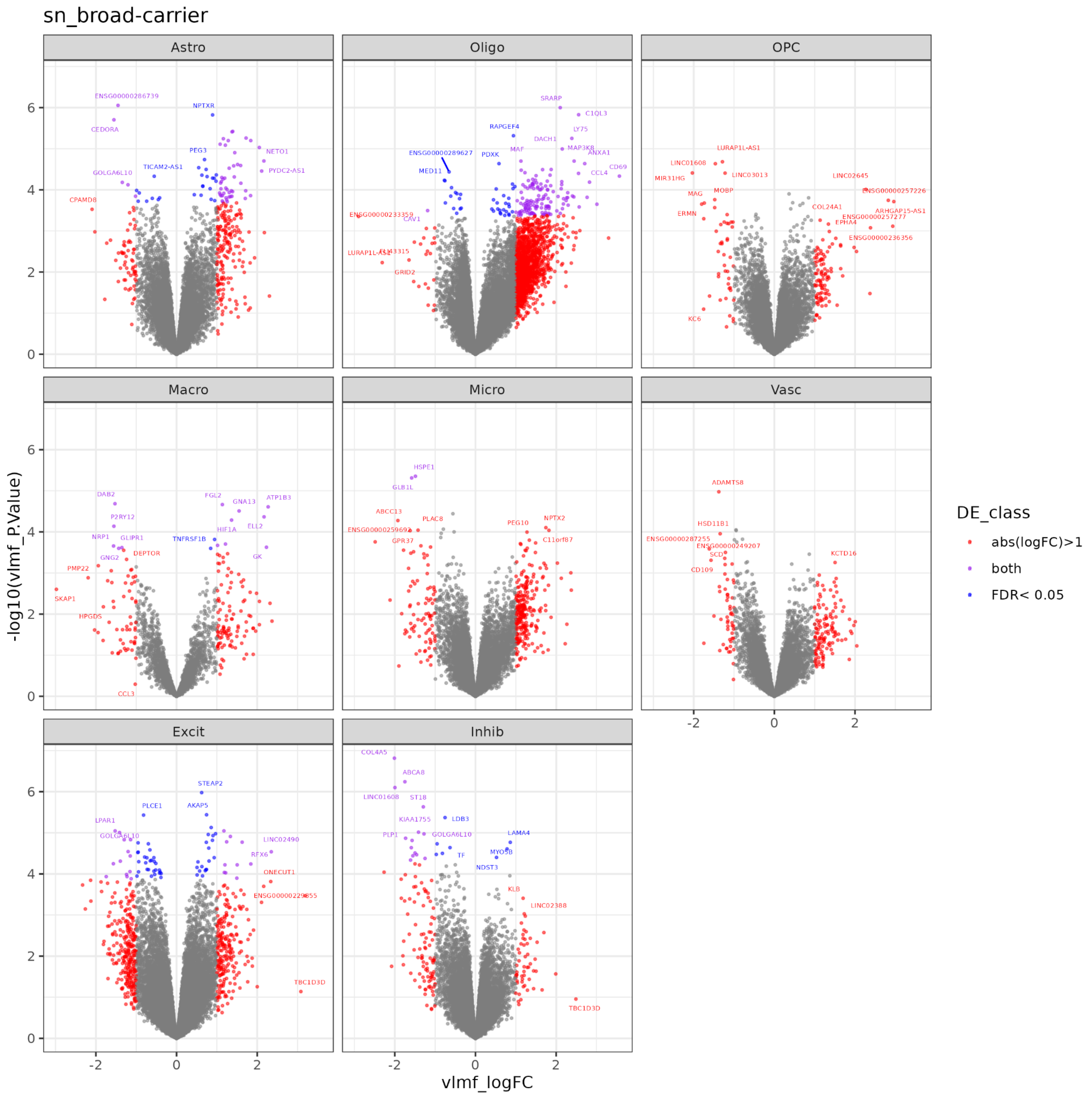


**Fig S39 Differential expression between *APOE* E4+ and E2+ carriers for snRNA-seq broad cell types.** Volcano plot for *APOE* carrier (E2+ vs. E4+) DE analysis for all broad cell types. Genes with FDR<0.05 or absolute $\log_2$ fold change (logFC) > 1 are indicated with point color. Related to **Figure 4, Table S11.**

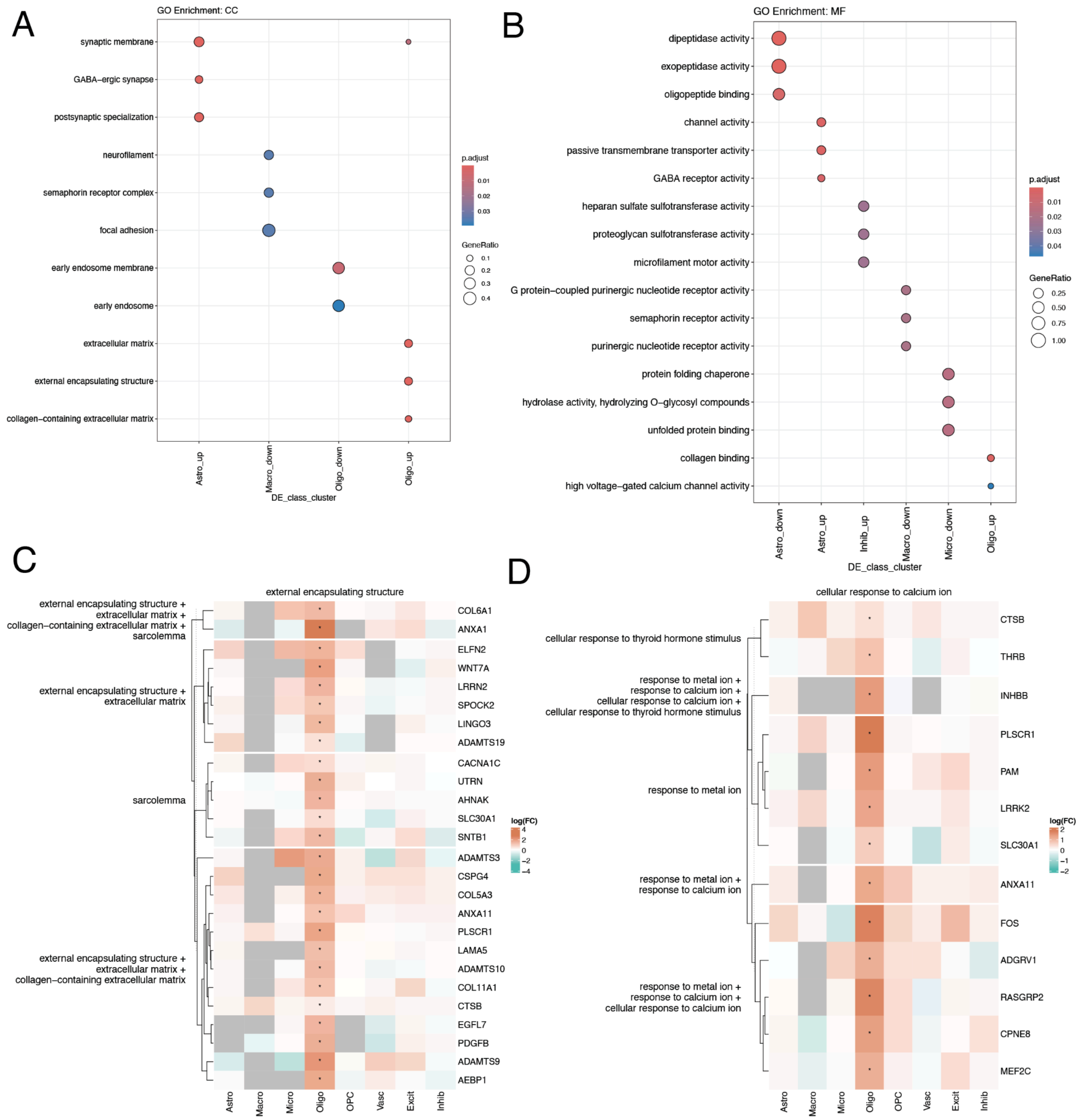


**Fig S40 Gene ontology analysis for broad cell type *APOE* carrier differential expression. A-B.** *clusterProfiler*[127] GO dot plot for "Cellular Component" (**A**) and "Molecular Function" (**B**). Gene ratio is the number of genes in each gene set annotated in a term (y-axis) over the total in the respective gene set. p.adjust is the FDR-adjusted p-value. **C-D.** Differential expression logFC heatmap for genes associated with "external encapsulating structure" (C) and "cellular response to calcium ion" (**D**) parent GO terms from the Biological Process ontology. Related to **Figure 4, Table S12.**

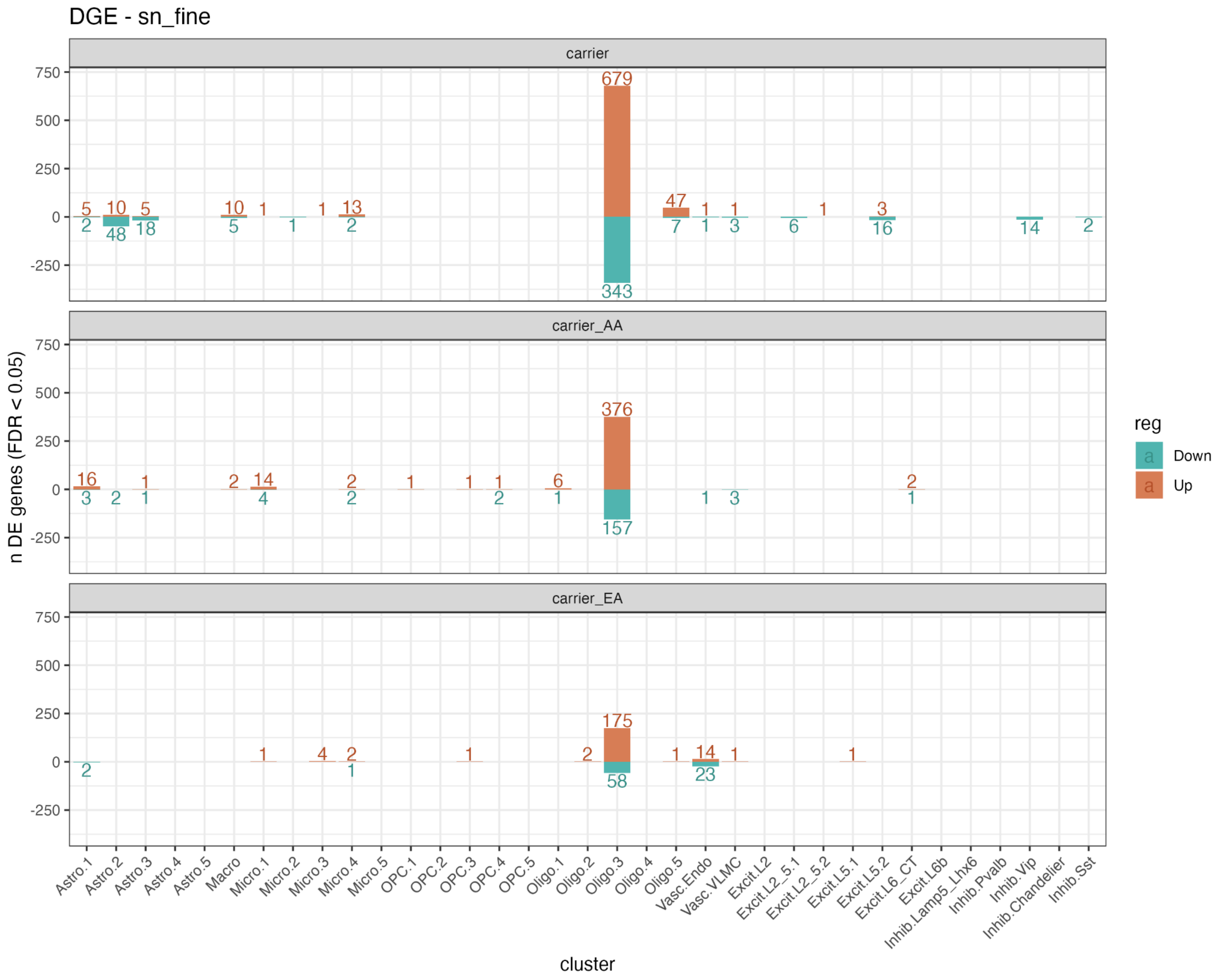


**Fig S41 Number of differentially expressed genes between *APOE* E2+ and E4+ carriers in snRNA-seq fine subclusters analysis.** Bar plots showing the number of *APOE* carrier (E2+ vs. E4+) DEG genes (FDR<0.05) identified for each snRNA-seq fine resolution subcluster either using all donors (top row), African Ancestry (AA) donors only, or European Ancestry (EA) donors only (**Methods: Differential expression analysis**). Related to **Figure 5**, **Table S13**.

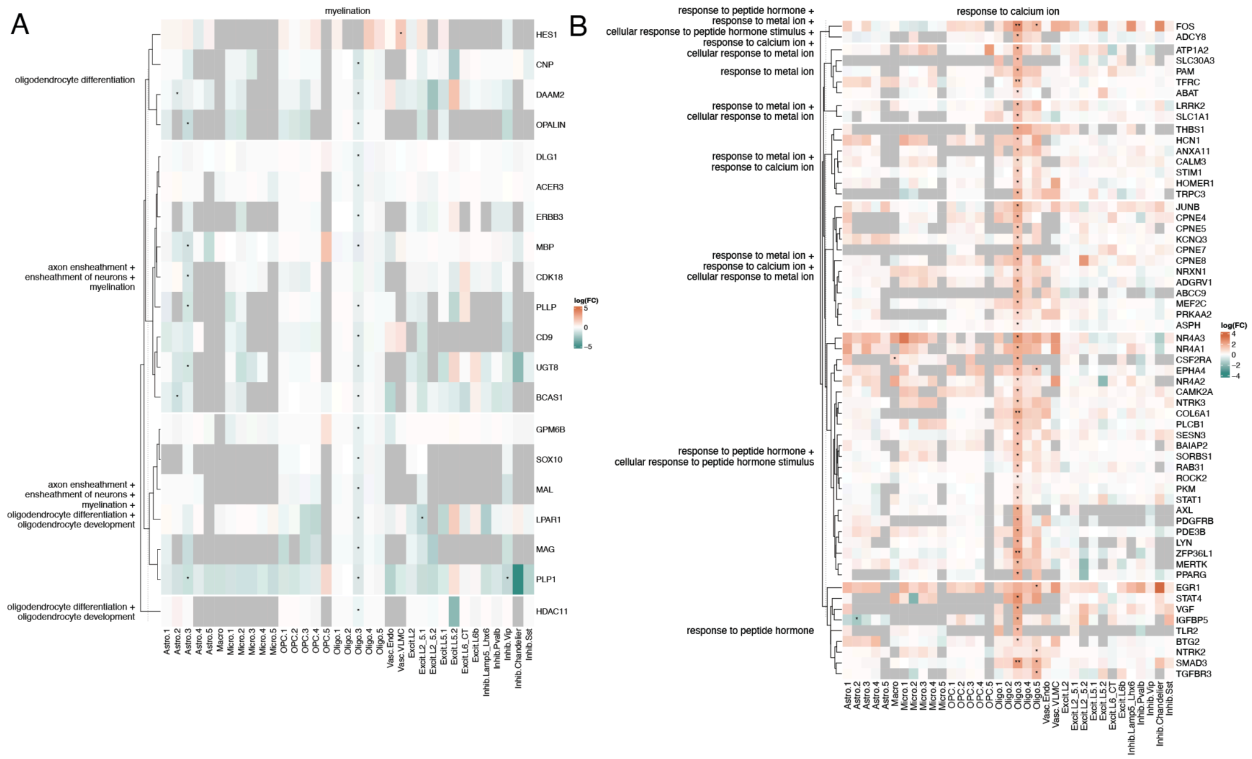


**Fig S42 Gene ontology analysis for fine subcluster *APOE* carrier differential expression. A.** logFC heatmap for genes associated with Biological Process (BP) GO terms with the parent term “myelination”. **B.** logFC heatmap for genes under “response to calcium ion” parent GO term from the BP ontology. Related to **Figure 5**, **Table S14**.

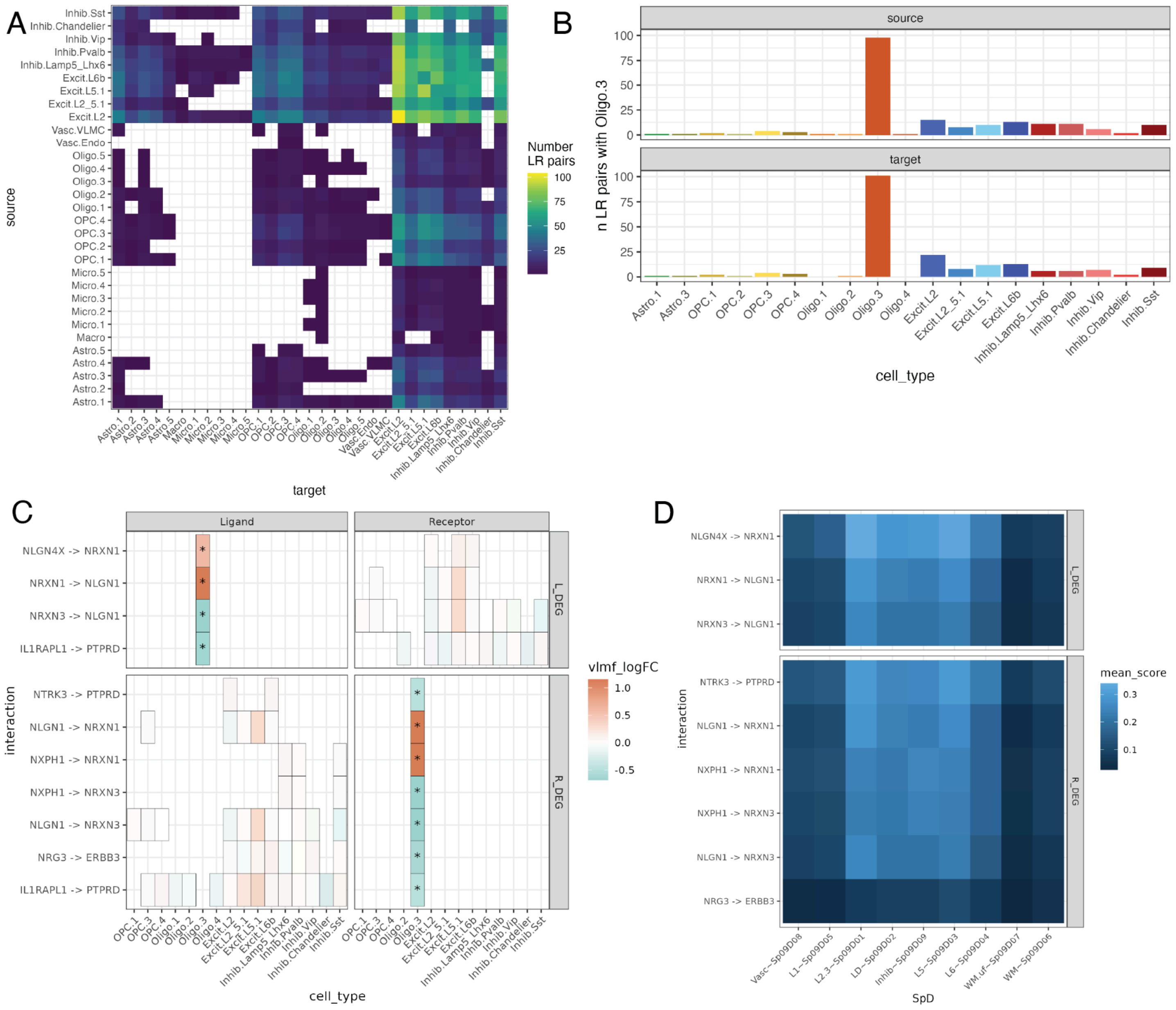


**Fig S43 Cell to Cell Communication identified recurrent ligand-receptor interactions between snRNA-seq subclusters. A.** Heatplot showing the number of commonly observed ligand-receptor (LR) pairs between source and target subclusters. Commonly observed LR pairs were defined as having a `magnitude_rank`< 0.05 in 20 of the 30 donors (**Methods: Cell-cell communication**). **B.** Barplot showing number of commonly observed LR pairs involving Oligo.3 as either source or receiver subcluster. **C.** Tile plot of LR pairs involving a differentially expressed gene as a ligand (L_DEG) or receptor (R_DEG). The color fill is given by the $\log_2$ fold change from differential expression for the source or target subclusters in the corresponding gene. Genes identified in both CCC and DE analysis only involved Oligo.3; these genes are annotated with an asterisk. There were no interactions where both ligand and receptor were DEGs. For example ligand *NRXN1* is upregulated in E4+ in source subcluster Oligo.3, the receptor *NLGN1* is not a DEG in any of the 5 target subclusters **D.** Heat plot of mean bivariate score, which measures tendency of each ligand and receptor to locally co-express throughout a given SpD, from *LIANA+*[59] for DEG-involved LR pairs from *C*. *IL1RAPL1* -> *PTPRD* failed bivariate evaluation. Related to **Figure 1**, **Table S16**.

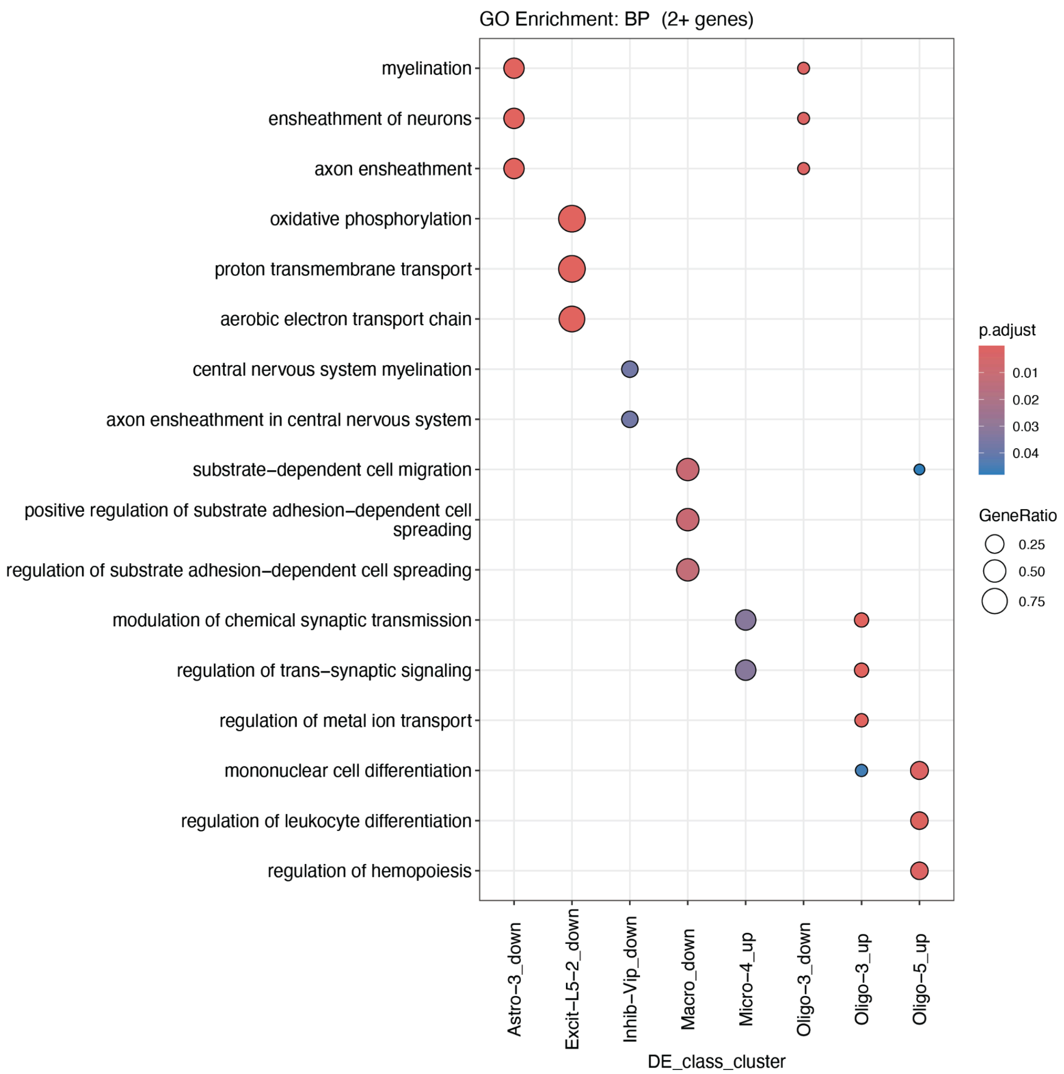


**Fig S44 Top GO terms for snRNA-seq subcluster DEGs.** *clusterProfiler*[127] GO dot plot for "Biological Process" showing the top 3 terms with 2 or more genes for each snRNA-seq subcluster. Gene ratio is the number of genes in each gene set annotated in a term (y-axis) over the total in the respective gene set. p.adjust is the FDR-adjusted p-value. Related to **Figure 5**.

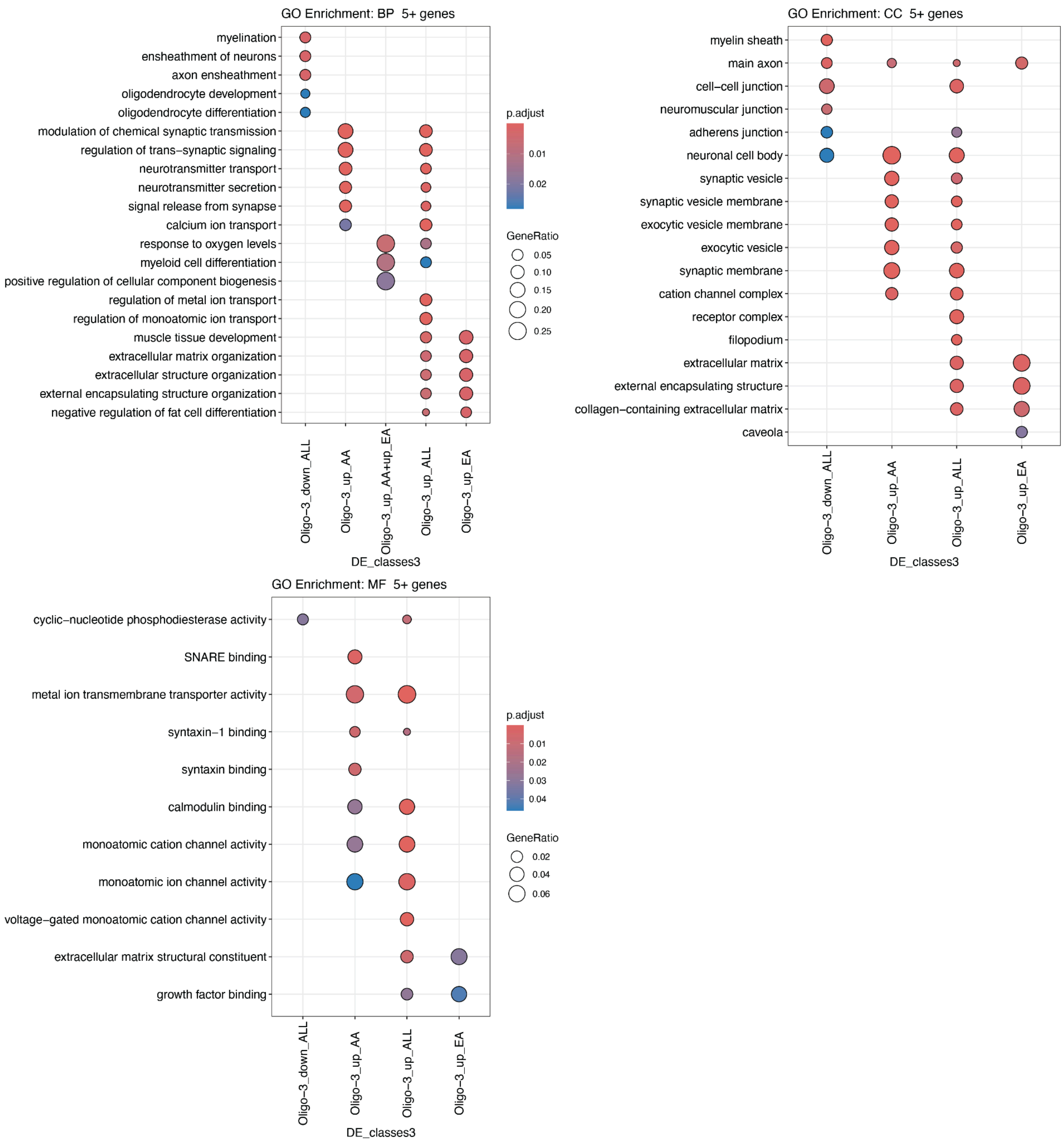


**Fig S45 Gene ontology analysis for Oligo.3 ancestry-specific DEGs.** *clusterProfiler*[127] GO dot plot for Biological Process (BP), Cellular Component (CC), and Molecular Function (MF) ontologies for top enrichment terms with 5 or more *APOE* carrier DEGs. Gene ratio is the number of genes in each gene set annotated in a term (y-axis) over the total in the respective gene set. p.adjust is the FDR-adjusted p-value. Related to **Figure 5F, Fig S37C+I, Table S14.**

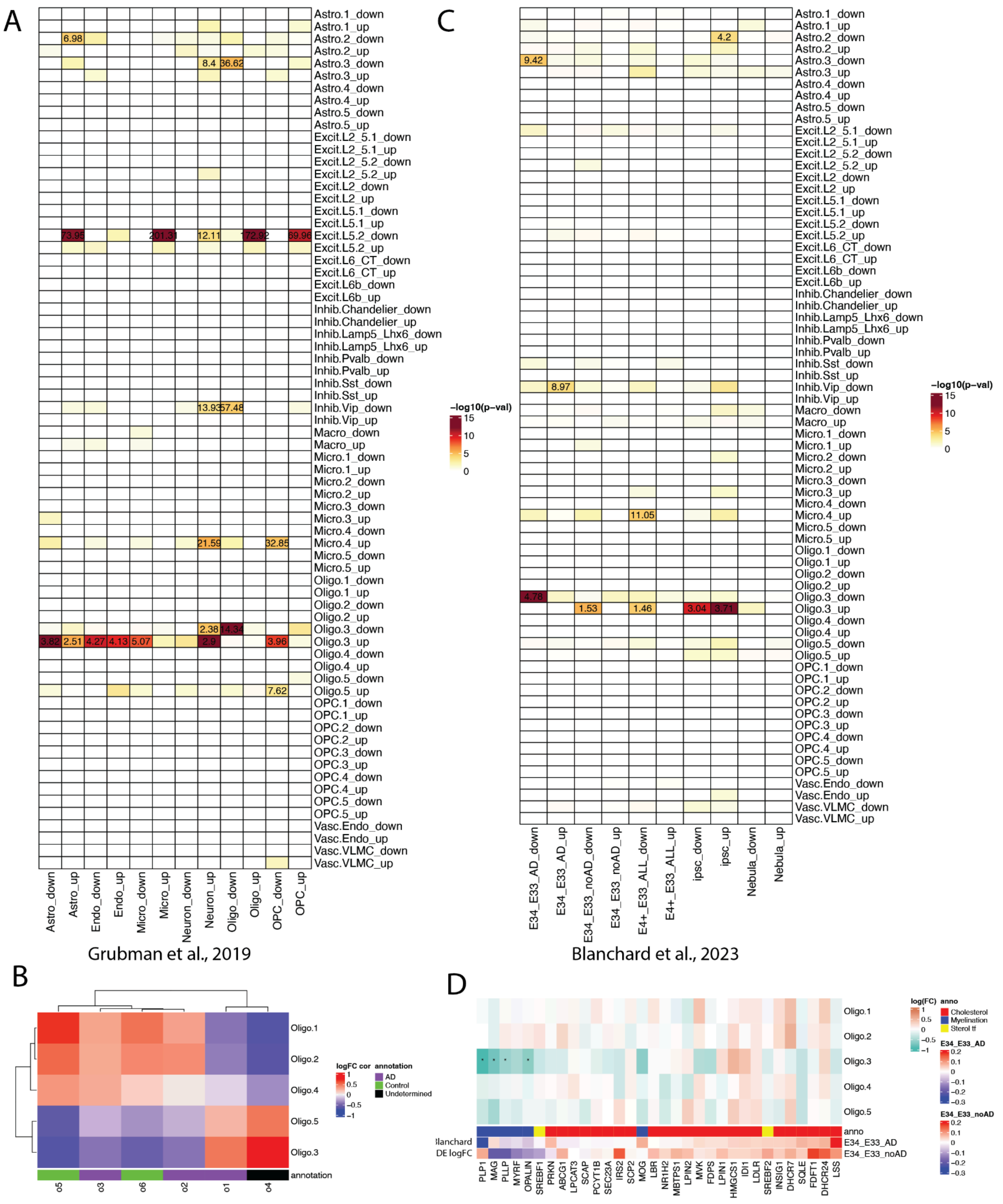


**Fig S46 Fine subcluster differential expression compared to previous studies. A.** Gene set enrichment heatmap of ERC fine subcluster DEGs compared to Grubman et al. DE analysis in postmortem ERC between AD vs control donors[19]. Downregulated genes in Excit.L5.2 in our dataset were enriched for upregulated DEGs in several cell types from Grubman et al.[19], which was mostly driven by mitochondrial genes. **B.** Correlation heatmap of logFC values from enrichment modeling of Oligo subclusters from Grubman et al.[19] compared to our ERC Oligo subclusters. Annotation bar is Grubman et al. classification of subclusters either mostly from AD or control donors, or undetermined. Oligo.3 had high correlation with Grubman et al. o4, and likewise between Oligo.1 and o5. **C.** Gene set enrichment with Oligo DEGs from Blanchard et al. comparing E3/E3 (E33) versus E3/E4 (E34) donors, and E4/E4 combined with E3/E4 (E4+) only among AD donors, among donors with no AD, or using all available donors using either a Wilcoxon rank-sum test or *Nebula*[17,139]. *Nebula* implements a fast negative binomial mixed model for DGE analysis[139] and which Blanchard et al. used for adjusting for donor as a random effect[17]. Blanchard et al. also generated data from iPSC-derived E4/E4 vs. E3/E3 oligodendrocytes[17]. **D.** logFC heatmap for Oligo subclusters for genes highlighted by Blanchard et al.[17] Annotation bars are gene function and Blanchard et al. reported logFC values from Wilcoxon rank-sum test between E3 and E4 homozygote *APOE* donors with and without AD. Related to **Fig S47**, **Figure 5**, **Table S15**.

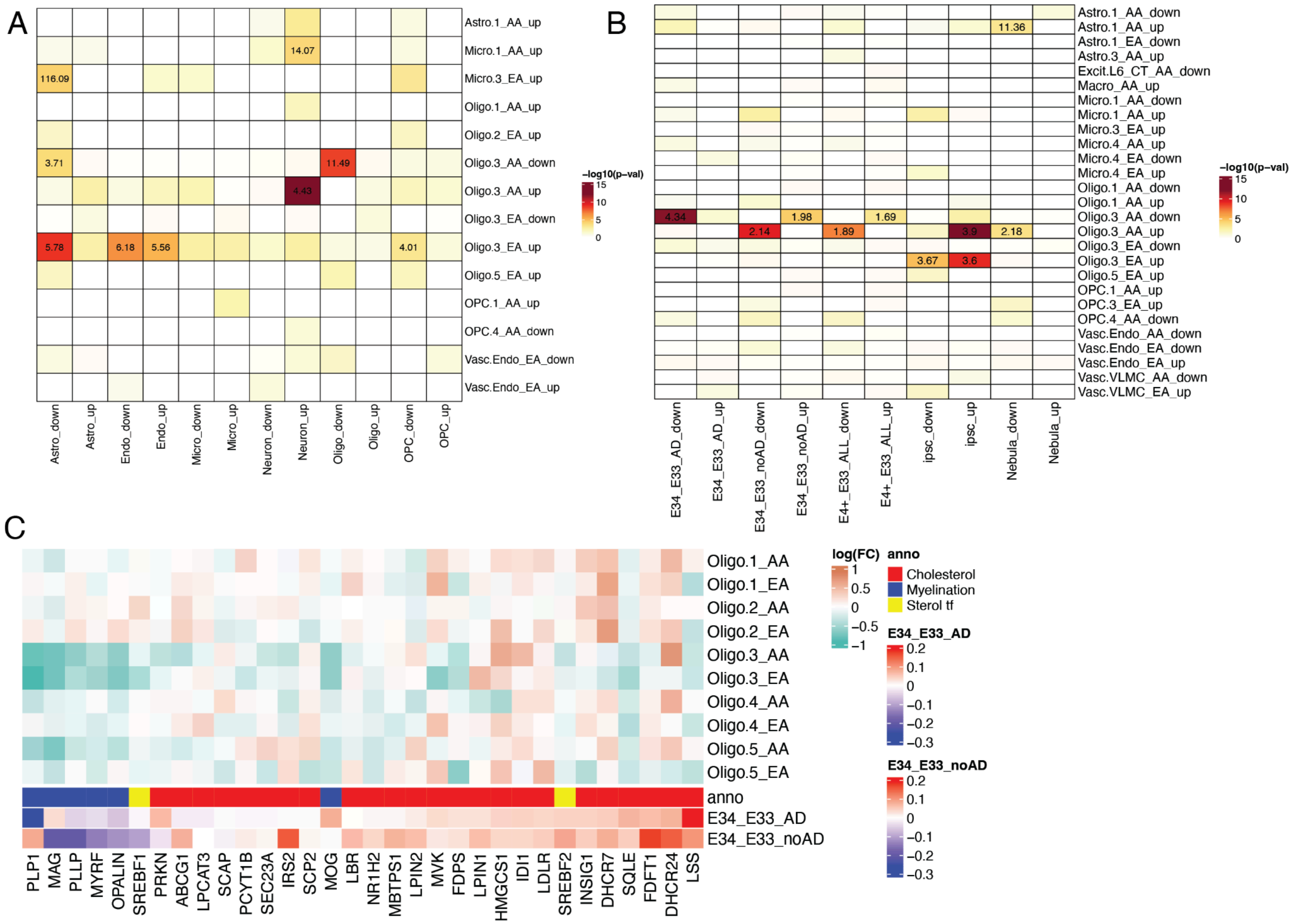


**Fig S47 Fine subcluster ancestry-specific differential expression compared to previous studies. A.** Gene set enrichment heatmap of ERC fine subcluster ancestry-specific DEGs compared to Grubman et al. DE analysis in postmortem ERC between AD vs control donors[19]. **B.** Gene set enrichment with Oligo DEGs from Blanchard et al. comparing E3/E3 (E33) versus E3/E4 (E34) donors, and E4/E4 combined with E3/E4 (E4+) only among AD donors, among donors with no AD, or using all available donors using either a Wilcoxon rank-sum test or *Nebula*[17,139]. **C.** Ancestry-specific logFC heatmap for Oligo subclusters for genes highlighted by Blanchard et al.[17] Annotation bars are gene function and Blanchard et al. reported logFC values from Wilcoxon rank-sum test between E3 and E4 homozygote *APOE* donors with and without AD. Related to **Fig S46**, **Figure 5**, **Table S15**.
